# Robust characterization of forest structure from airborne laser scanning – a systematic assessment and sample workflow for ecologists

**DOI:** 10.1101/2024.03.27.586702

**Authors:** Fabian Jörg Fischer, Toby Jackson, Grégoire Vincent, Tommaso Jucker

## Abstract

1. Forests display tremendous structural diversity, shaping carbon cycling, microclimates, and terrestrial habitats. One of the most common tools for forest structure assessments are canopy height models (CHMs): maps of canopy height obtained at high resolution and large scale from airborne laser scanning (ALS). CHMs can be computed in many ways, but little is known about the robustness of different CHM algorithms and how they affect ecological analyses.
2. Here, we used high-quality ALS data from nine sites in Australia, ranging from semi-arid shrublands to 90-m tall Mountain Ash canopies, to comprehensively assess CHM algorithms. This included testing their sensitivity to point cloud degradation and quantifying the propagation of errors to derived metrics of canopy structure.
3. We found that CHM algorithms varied widely both in their height predictions (differences up to 10 m, or 60% of canopy height) and in their sensitivity to point cloud characteristics (biases of up to ∼5 m or 40% of canopy height). Impacts of point cloud properties on CHM-derived metrics varied, from robust inference for height percentiles, to considerable errors in aboveground biomass estimates (∼50 Mg ha^−1^, or 10% of total), and high volatility in metrics that quantify spatial associations in canopies (e.g., gaps or spatial autocorrelation). In some cases, biases exceeded ecological variation across sites by a factor of 2. However, we also found that two CHM algorithms – a variation on a “spikefree” algorithm that adapts to local pulse densities and a simple Delaunay triangulation of first returns – allowed for robust canopy characterization and should thus create a secure foundation for ecological comparisons in space and time.
4. Canopy height models are a widely used tool in ecology, but their derivation is not trivial. Our study provides a best-practice guideline and a sample workflow to create robust CHMs and minimize biases and uncertainty in downstream analyses. In doing so we pave the way for global-scale comparisons of forest structural complexity from airborne laser scanning.

## 1. Introduction

Forests across the globe display tremendous structural diversity, ranging from open woodlands to tall, dense, multi-layered tropical forests. Quantifying structural differences across and within forest ecosystems is vital, as they shape climate at both small (De Frenne et al., 2021) and large scales (Alkama & Cescatti, 2016), are tightly coupled to carbon fluxes (R. Fischer et al., 2019), influence the resilience of forests to disturbance (Koontz et al., 2020) and create habitats for forest-dwelling organisms (Gouveia et al., 2014).

For a long time, the main challenge to quantifying the vertical and horizontal structure of forest canopies was data acquisition. It involved the painstaking delineation of canopy gaps from the ground (Meer & Bongers, 1996) or the manual measurements of tree crowns to generate canopy profile diagrams (Baker & Wilson, 2000). The advent of remote sensing has radically changed this, with ever-increasing opportunities to quantify forest structural complexity (Atkins et al., 2018; Disney, 2019; Duncanson et al., 2022; Ehbrecht et al., 2021; F. J. Fischer & Jucker, 2023). In particular, airborne laser scanning (ALS) has emerged as a key technology in this field, allowing researchers to probe forest canopies from above using high-frequency laser scanners mounted on airplanes, helicopters or even drones. Whenever a laser pulse emitted by the scanner hits an obstacle – a leaf, a branch, or the ground – the reflected energy spike can be converted into a geolocated point, capturing the forest as a 3D “point cloud”. The unique value of ALS is the ability to characterize forest structure and topography in high detail (e.g., 1-m resolution), while also covering areas large enough (e.g., 10s of km^2^) to map landscape patterns (Lines et al., 2022).

A particularly useful feature of ALS point clouds is that they can be used to generate high-resolution maps of canopy height, known as canopy height models (CHMs). CHMs rely mostly on top-canopy pulse returns and are thus expected to be more robust to instrumentation and acquisition parameters than the underlying point clouds (Asner, Mascaro, et al., 2013; Mielcarek et al., 2018). They are also smaller in data volume and easier to manipulate, share and interpret. As a result, they have found widespread application in ecology, including in carbon mapping (Asner, Mascaro, et al., 2013; Coomes et al., 2017; Labriere et al., 2018; Xu et al., 2017), tree delineation (Aubry-Kientz et al., 2019; Kaartinen et al., 2012), modelling of forest inventory attributes (Bottalico et al., 2017; Næsset, 2002), studies of canopy dynamics and gaps (Asner, Kellner, et al., 2013; Dalagnol et al., 2021; Goodbody et al., 2020; Huertas et al., 2022), the calibration of forest models (F. J. Fischer et al., 2020), and habitat assessments (Davies & Asner, 2014; Zellweger et al., 2013).

With large-scale, multi-temporal coverage available in many countries, ALS is now a tool ready for global ecological analysis (Jucker, 2022). However, to leverage its full potential for ecological research, standardised approaches to characterising canopy 3D structure are needed (Moudrý et al., 2023). Considerable effort has gone into optimising ALS-derived terrain models (Andrade et al., 2018; Chen et al., 2017) and quantifying canopy metrics from ALS point clouds (Almeida et al., 2019; Magnussen et al., 2010; Næsset, 2009; Pearse et al., 2019; Riofrío et al., 2022; Roussel et al., 2018; Vincent et al., 2023; Wilkes et al., 2015). In contrast, there is little guidance for ecologists on how to generate robust CHMs that are comparable across datasets acquired with different instruments and sampling designs. This is particularly problematic for studying canopy dynamics across time, as bias and uncertainty between multiple airborne lidar acquisitions can easily exceed the average canopy growth over a few years (Bruggisser et al., 2019; Huertas et al., 2022; Roussel et al., 2017).

The most frequent approach to obtaining CHMs is to rasterize the highest point at a target resolution and then subtract the resulting surface model from a topographic model. This is accurate when pulse densities are high (e.g., ≥20 pulses m^−2^), but will underestimate canopy height at low pulse densities, as laser shots miss branches and leaves or fail to produce ground returns (LaRue et al., 2022; Leitold et al., 2015). Pulse density is likely the strongest determinant of CHM quality, but bias can also result from variations in laser energy, scan angles and footprint size (Keränen et al., 2016; Næsset, 2009; Roussel et al., 2017, 2018). Several algorithms have been developed to improve CHM robustness (e.g., Khosravipour et al., 2014, 2016), some of which are available in open-source software such as the lidR R package (Roussel et al., 2020). However, systematic assessments of CHM algorithms are rare and have often been limited to single acquisitions or reference height points (Mielcarek et al., 2018; Quan et al., 2021). Only an in-depth understanding of how CHM derivation impacts canopy structural metrics across a diversity of forest types will allow us to mobilize the wealth of information that ALS provides for ecology.

Here we use ALS data from nine sites in Australia that cover a wide range of ecosystems – from open acacia woodlands to dense rain forests and 90-m tall Mountain Ash stands – and assess the robustness of six different CHM algorithms. Two of these are new takes on existing approaches, designed to be less error-prone. By systematically manipulating laser point clouds, we test: (1) What are the main sources of uncertainty and bias in CHMs? (2) Which algorithms are most robust to differences in acquisition parameters and point cloud degradation? And (3) how do errors in CHM construction propagate to canopy structural attributes such as canopy gaps, canopy heterogeneity and aboveground biomass estimates?

We summarize our results into 16 best-practice guidelines for ecologists and provide a versatile pipeline for producing robust CHMs in R. In doing so we aim to promote the uptake of ALS data across a wide range of ecological fields and facilitate efforts to leverage the growing access to ALS data for large-scale and multi-temporal analyses.

## 2. Methods

### 2.1 ALS data

ALS data were obtained from nine SuperSites belonging to Australia’s Terrestrial Ecosystem Research Network (TERN; Karan et al., 2016). These sites provide an ideal testbed for assessing the robustness of CHM algorithms. For one, they range widely in mean annual temperature (8.4 – 27.3° C), annual precipitation (260 – 1950 mm) and elevation (50–1000 m a.s.l.), resulting in a considerable variety of canopy structures: maximum heights between 10–90 m and 30–100% canopy cover (Table 1). Moreover, all sites were systematically scanned in 2012-15 over a consistent extent (5×5 km^2^) and with the same Riegl Q560 scanner. The scanner’s outgoing pulse rate was 240 kHz with an angular sweep of 45°. The vast majority of sites were scanned with 41 parallel flightlines (aligned either N-S or E-W), spaced 125 m apart. Exceptions were tall Eucalypt stands at Warra, Tasmania (50 flightlines), and Robson Creek, a dense, high-biomass rain forest in northern Queensland, which was flown across two days (112 flightlines, both N-S and E-W) to capture the site’s complex canopies and topography. Throughout, average flying height was 300 m, resulting in a maximum footprint diameter of 0.15 m and high sampling densities (19–31 pulses m^−2^).

**Table 1:** Forest and scan characteristics across the nine TERN SuperSites. Coordinates, elevation and canopy properties are derived from digital terrain models (DTMs) and canopy height models (CHMs) calculated from the original scans via the “locally adaptive spikefree” method (CHM_lspikefree_, see Section 2.2 for details). Maximum canopy height was calculated as the 99^th^ percentile of the CHM, while canopy cover is defined as the proportion of CHM pixels taller than 2 m above-ground. Sites are ordered by maximum canopy height.

| Site | Lon | Lat | Forest type | Elevation<br>(m a.s.l.) | Maximum<br>canopy<br>height (m) | Canopy<br>cover (%) | Pulse<br>density<br>(m<sup>-2</sup>) | Ratio<br>2<sup>nd</sup> / 1<sup>st</sup><br>returns |
| --- | --- | --- | --- | --- | --- | --- | --- | --- |
| Alice Mulga | 133.25 | -22.28 | Semi-arid acacia<br>woodland | 600 | 8.0 | 65 | 24.9 | 0.51 |
| Chowilla | 140.59 | -34.00 | Semi-arid mallee<br>shrubland | 50 | 10.1 | 43 | 20.6 | 0.33 |
| Credo | 120.66 | -30.19 | Semi-arid temperate<br>woodland | 450 | 22.3 | 29 | 21.5 | 0.24 |
| Litchfield | 131.79 | -13.18 | Tropical savanna | 220 | 25.6 | 49 | 20.8 | 0.31 |
| Rushworth<br>Forest | 144.96 | -36.75 | Dry sclerophyll forest | 200 | 25.6 | 71 | 19.1 | 0.54 |
| Robson<br>Creek | 145.62 | -17.11 | Seasonal tropical<br>rainforest | 870 | 53.6 | 100 | 31.0 | 0.45 |
| Zigzag<br>Creek | 148.34 | -37.47 | Dry/wet sclerophyll<br>forests | 280 | 55.0 | 92 | 19.2 | 0.34 |
| Warra | 146.66 | -43.10 | Wet sclerophyll forest | 250 | 75.1 | 97 | 20.2 | 0.57 |
| Watts<br>Creek | 145.69 | -37.69 | Wet sclerophyll forest | 990 | 87.0 | 97 | 19.0 | 0.66 |

### 2.2 A standardized workflow for generating CHMs from ALS point clouds

#### Point cloud processing

We developed a standardized pipeline to convert ALS point clouds into digital terrain models (DTMs) and canopy height models (CHMs). The pipeline is executed from R (R Core Team, 2023) and processes data with a combination of LAStools commands (Isenburg, 2023, called directly from R via “system()”) and custom functions written in R, mostly relying on the lidR, terra and data.table packages (Dowle & Srinivasan, 2023; Hijmans, 2023; Roussel et al., 2020). The pipeline was developed for the automatic processing of large volumes of data with variable structures and has been extensively tested across a global collection of ALS acquisitions (). Further details on the pipeline and R packages can be found in the Supplementary Information (S1), the code and a sample workflow are available on Zenodo (http://doi.org/10.5281/zenodo.10878070). More recent versions of the pipeline are available on github (). For the purposes of this analysis, ALS point clouds were processed in tiles of 500 m, with buffers of 25 m. All processing was carried out in parallel on an Intel® Core™ i9-10980XE CPU with 18 physical cores and 64 GB RAM. Processing times varied from less than a minute (single flightline) to several hours (scans at sites with dense point clouds, such as Robson Creek).

#### CHM algorithms

As part of the processing pipeline, we used six alternative CHM algorithms. Four are part of common software tools (LAStools, lidR) and run with recommended parameterizations to generate 1-m resolution CHMs (see Table S2 for details). They include a simple rasterization of the highest return per m^2^ (CHM_highest_), a TIN-interpolation of first returns (CHM_tin_), a “pitfree” algorithm that builds the canopy surface by merging multiple TIN-interpolations (CHM_pitfree_, Khosravipour et al., 2014) and the “spikefree” algorithm, a vertically constrained TIN-interpolation (CHMspikefree, Khosravipour et al., 2016). The other two CHM algorithms are custom variations conceived specifically for this study: a spikefree algorithm applied to a thinned-down point cloud (CHM_tspikefree_), and a spikefree algorithm that adapts its smoothing to local pulse density variation (CHM_lspikefree_, first returns only in both cases). The idea behind both modifications was that even within a single ALS acquisition, pulse densities are higher in areas closer to the scanner (centre of flightlines, tall trees or hill tops) and areas of flightline overlap, which can bias height estimates (Roussel et al., 2017). For example, the default spikefree algorithm might over-smooth in high and insufficiently remove spikes in low pulse density areas, resulting in over- and underestimation of height, respectively. By either removing local pulse density variation or adapting the spikefree parameters, we attempted to solve this problem.

We used all algorithms to first create digital surface models (DSMs) and then obtained CHMs by subtracting ground elevations (i.e., the DTM, based on LAStool’s “lasground_new”). The exception was CHM_pitfree_ which we computed directly from a height-normalized point cloud to reduce computation time. A DSM_pitfree_ was constructed post hoc by summing CHM_pitfree_ and DTM.

### 2.3 Robustness of CHM generation and forest structure assessment

#### Simulating point clouds of varying quality

Pulse density (the number of laser pulses per unit area) is usually the best first-order approximation of airborne laser scan quality, particularly for CHM construction. Pulse density depends on the frequency with which pulses are emitted, but also summarizes flightline properties, such as altitude, speed and overlap. It is highest when the laser is flown close to the canopy, at low speed and in narrowly spaced flightlines. As a general rule, the higher the pulse density, the less sensitive should a canopy reconstruction be to other laser properties. For example, if 1 m^2^ of canopy is hit by a single shot, it matters a lot where the shot came from, how wide its footprint was, how many returns it generated, and how the raw energy waveform was converted into a single point (Dayal et al., 2022; Keränen et al., 2016; Roussel et al., 2017, 2018). By contrast, if the same 1 m^2^ of canopy is saturated with shots from all directions (e.g., ≥ 20 pulses m^−2^), the canopy surface is likely well-approximated irrespective of these properties.

Pulse density is not only an important summary statistic of laser scans, but also easy to manipulate post hoc through a random (or targeted) thinning of laser shots. Here we used this approach to study how robust CHM algorithms and derived forest properties were to variations in point cloud quality. Starting from the original high-quality TERN ALS data, we created systematically degraded point clouds with pulse densities of 16, 8, 4, 2, 1 and 0.5 m^−2^ for each of the nine study sites. The thinning of laser shots was carried out using the homogenize() function in the lidR package, which simultaneously thins and homogenizes point clouds, so that every grid cell at a target resolution (here 25 m) has the same overall pulse density. We then ran the processing pipeline across the degraded point clouds and created standard DTM and CHM products for further analysis.

While pulse density is the best overall quality indicator, other scan properties can also affect CHMs, particularly at low pulse densities. To assess these effects, we carried out three companion analyses: (1) a custom degradation of pulse densities that, unlike homogenize(), removed every n^th^ shot, to assess the effects of within-scan heterogeneity, (2) a systematic degradation of laser penetration levels, i.e. the number of returns per shot, to investigate the effects of variation in higher-order returns on CHM algorithms, and (3) a splitting of the point clouds into flightlines, to study the effects of varying scan angles. For simplicity, we focus on pulse density variation throughout the main text and present details of the companion analyses in the Supplementary (Sections S2-S4).

#### Effects of point cloud quality on CHM generation

The way in which the algorithms described above interpolate laser returns to create DSMs is a major source of variability in CHMs. However, some variation is also expected due to errors in ground classification and DTM derivation. For example, at low pulse densities, canopy height could be underestimated because of insufficient sampling of treetops (negative bias in DSM) or insufficient sampling of the ground (positive bias in DTM). We here evaluated the two aspects separately: for each site and each pulse density level, we compared both the six algorithm-specific DSMs and the DTM to their reference products at the highest pulse density level (16 pulses m^−2^). Robustness of algorithms was assessed through the mean errors (ME, or bias, in m) and root mean square errors (RMSE, in m) of 1 m^2^ pixels. Throughout, we will visualize DSM/DTM stability across all pulse density levels, but for simplicity, only report RMSE/ME values for the most extreme but still acceptable point cloud differences (2 vs. 16 pulses m^−2^). To avoid edge effects, the outer 25 m of all DSMs and DTMs were removed.

#### Effects of point cloud quality on CHM-derived metrics

To assess how pulse density degradation propagated to CHM-derived forest structural metrics, we calculated 25 summary statistics from all CHMs at both 1 ha (100×100 m) and 1 km^2^ (1000×1000 m) resolution. Metrics were grouped into two categories: “vertical” ones that summarized the height distribution in each grid cell irrespective of spatial associations between the underlying 1 m^2^ pixels, and “horizontal” ones that explicitly assessed the connectivity of the 1 m^2^ pixels within each cell. The former included metrics such as the mean, standard deviation, and percentiles of canopy height, while the latter captured spatial autocorrelation (Moran’s I), height gradients (rumple index) and the clumping of canopy structures (gaps below and canopy clusters above height thresholds). We also assessed variation in the power law slopes of gap size frequency distributions measured at 1 km^2^ resolution (Silva et al., 2019). Moreover, for one site (Robson Creek) we estimated the impacts of CHM uncertainty and bias on above ground biomass (AGB) estimation. This was done by regressing field estimates of AGB from 25 1-ha plots against CHM-derived estimates of median canopy height (Labriere et al., 2018). A complete list of metrics can be found in Table S3, with a more detailed explanation of the AGB estimation in section S5.

For each metric, we again calculated MEs and RMSEs, but at 1 ha and 1 km^2^ resolution, between highly degraded point clouds and the reference point cloud (2 vs. 16 m^−2^), both within and across sites. To make errors comparable across metrics and relate them to ecological variation, we divided the MEs and RMSEs of all metrics by the standard deviations of the same metrics (hereafter rME and rRMSE). For example, the mean height of 1 ha grid cells is a robust metric for local structure assessments if its typical bias and uncertainty are much lower than the typical height variation within sites. To assess the practical effects of mixing high and low-quality point clouds in ecological analyses, we randomly swapped 50% of cells between 2 and 16 m^−2^ grids (for each site, both at 1 ha and 1 km^2^ resolution) and calculated Spearman’s R2 between the newly assembled grids (R2_rankswapped_). A high value (> 0.9) indicates that the ranking of grid cells is preserved and that forest structure assessments are unlikely to change even when low and high pulse density areas occur in the same analysis.

## 3. Results

### 3.1 Sources of uncertainty and bias in canopy height estimation

Canopy height varied strongly across sites, algorithms and pulse densities (Figs 1-2 and Tables S4-5). Mean canopy height ranged from 2.5 m at Alice Mulga (average across algorithms) to 36.1 m at Watts Creek (Δh = 33.6 m). However, values depended strongly on the CHM algorithm: the same sites ranged from 3.4 m to 39.7 m for CHM_highest_ (Δh = 36.3 m) and from 1.7 m to 28.9 m for CHM_tin_ (Δh = 27.2 m). The largest relative difference between algorithms at a single site was observed at the savanna site Litchfield where many low canopy returns resulted in a mean height of 2.9 m for CHM_tin_ (60% difference), but 7.3 m for CHM_highest_.

**Fig 1:**
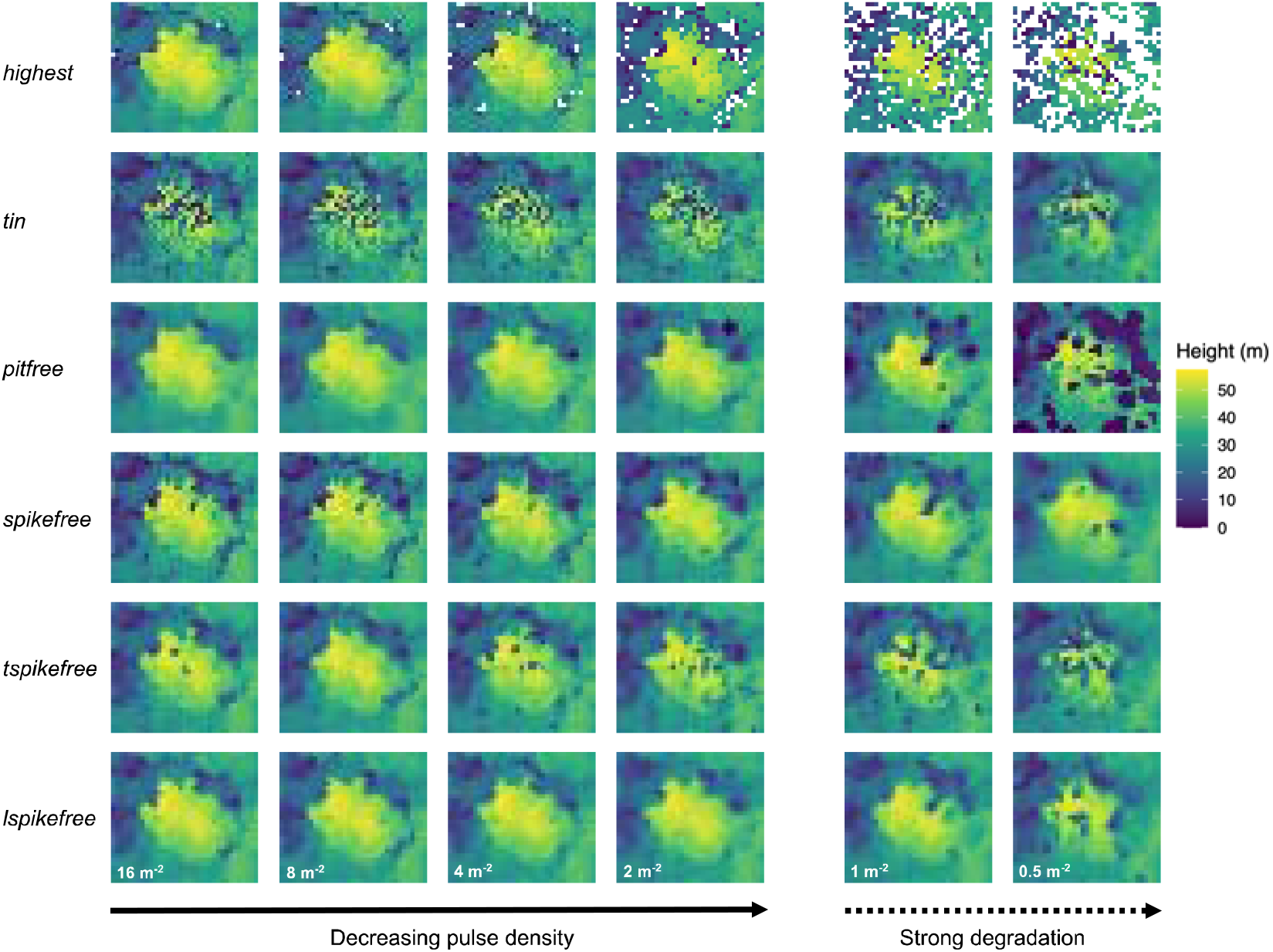
Canopy height models across pulse densities for a single tree at Robson Creek. Shown are canopy height models (CHMs, 1 m resolution) for a tree at Robson Creek (crown diameter = 23 m) from 16 down to 0.5 pulses m^−2^. Each row corresponds to a CHM algorithm. Algorithms degrade severely below 2 pulses m^−2^, which is why robustness was assessed only above this threshold (see Figs S1-4 for a visualization of CHMs for different types of degradations and sites).

**Fig 2:**
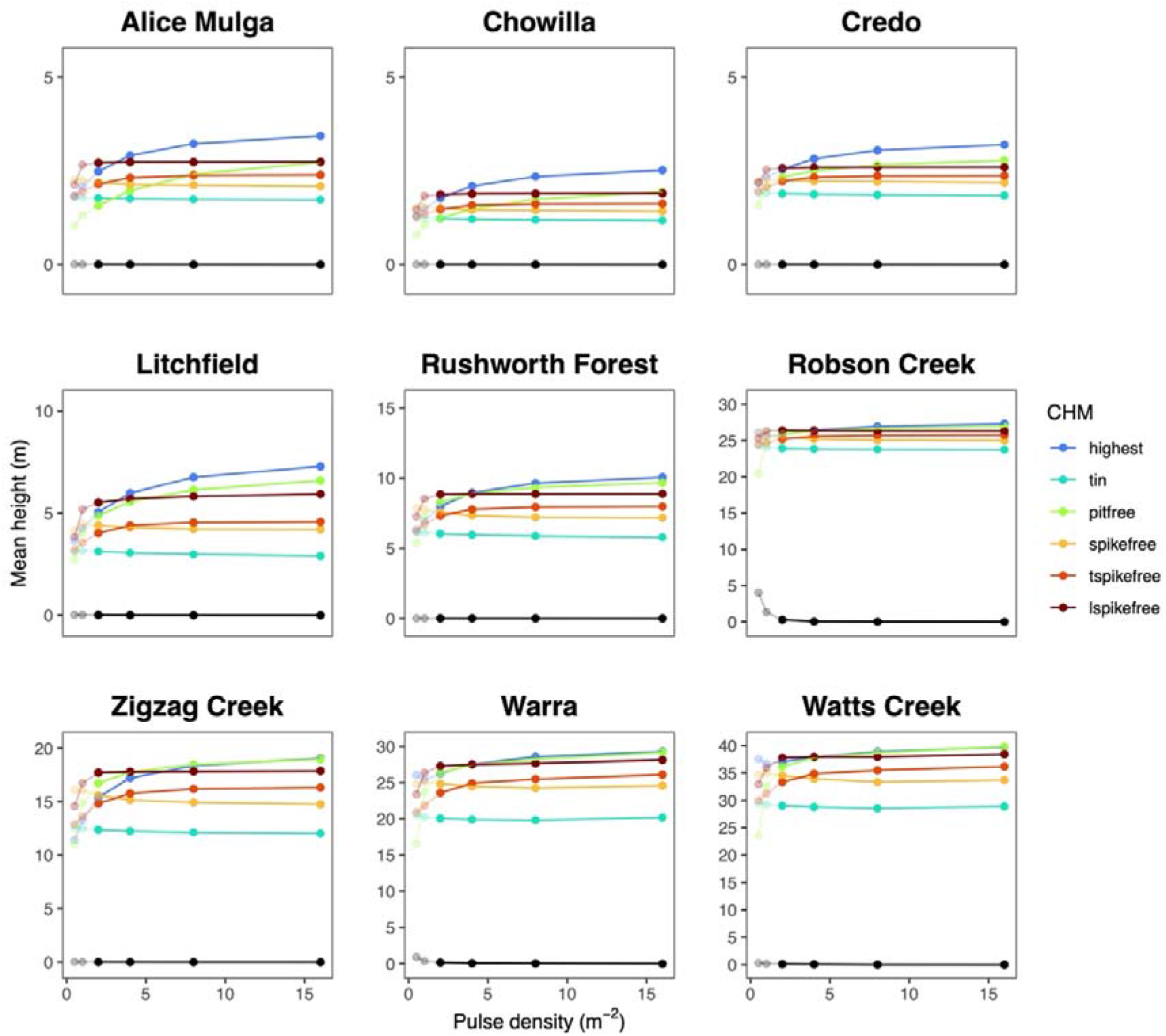
Changes in mean canopy height across pulse densities and algorithms for all TERN SuperSites. Shown is mean canopy height from 0.5 to 16 pulses m^−2^ across canopy height model (CHM) algorithms and study sites. To separate out effects on topography from effects on canopy surfaces, CHMs are computed by subtracting a reference digital terrain model (DTM) generated using high-quality point clouds (16 pulses m^−2^) from digital surface models (DSMs) generated using increasingly degraded point clouds. Also shown is a pulse-density specific DTM (black line), normalized by the reference DTM of each site. Changes below 2 pulses m^−2^ are plotted with transparency, as they were not used for robustness assessments. Figs S7-8 show the equivalent for laser penetration degradation and scan angle differences.

Variation in pulse density had a strong effect on canopy height estimates, with decreases from 16 m^−2^ to 2 m^−2^ resulting in a mean absolute bias of up to −5.64 m for tall canopies (CHM_highest_ at Watts Creek) and relative biases of up to −42 % for short canopies (CHM_pitfree_ at Alice Mulga). Below 2 pulses m^−2^, deviations became even more extreme (Figs 1-2 and Table S5). Pixel-level RMSEs reached up to 14 m between 16 and 2 pulses m^−2^ (Watts Creek, CHM_highest_) or >100% of mean canopy height (Litchfield, CHM_tin_), with comparable uncertainties due to within-scan heterogeneity (Figure S5). Most of these errors and uncertainties were due to the CHM algorithms, with little additional effects due to DTM generation (bias ≤ 0.5 m for most sites). The tropical rain forest at Robson Creek was an exception, with smaller relative changes in canopy height than the other sites, dropping by only 3.6 m (13%) from CHM_highest_ to CHM_tin_ and 1.91 m (7%) from 16 to 2 pulses m^−2^ for CHM_highest._ At the same time, Robson Creek also exhibited the strongest DTM degradation, with positive biases of 2-5 m at low pulse density, low laser penetration and oblique scan angles (Fig 2, Fig S9, Tables S5).

### 3.2 Robustness of CHM algorithms

Among the six algorithms we tested, CHM_lspikefree_ was the most robust, followed by CHM_tin_, with both showing little degradation between 16 and 2 pulses m^−2^, even in tall canopies (maximum bias of −0.52 m and +0.57 m respectively, Table S5). However, between the two, CHM_tin_ was much noisier than CHM_lspikefree_ which had the overall lowest RMSE (Fig 1 and Table S5). Deviations in the other four algorithms were variable, but usually much larger (Table S5). For example, at 2 pulses m^−2^, biases in canopy height estimated from CHM_highest_ and CHM_pitfree_ routinely reached several metres. In addition, CHM_highest_ degraded rapidly in coverage (∼4% NA values at 2 pulses m^−2^, ∼42% at 0.5 m^−2^). A visual assessment of algorithms confirmed the highest consistency in CHM_lspikefree_ and the clearest degradation in CHM_highest_ (Fig 1, Figs S1-S4). However, it also showed that, at high pulse densities, contours were sharper in CHM_highest_ and CHM_spikefree_ than in CHM_tin_ (too noisy) and CHM_lspikefree_ (too smooth). Results were broadly consistent with companion analyses of scan angle and laser penetration, where CHM_lspikefree_-based estimates also came out as the most robust, followed by CHM_tin_ (Table S5). However, we did observe that even for CHM_lspikefree_ and CHM_tin_, canopy height was systematically underestimated at larger scan angles (15-20° difference, Table S5 and Fig S4).

### 3.3 Robustness of CHM-derived forest structure analysis

Downstream analyses were strongly affected by errors in CHM creation. We compared the calculation of a total of 25 metrics between 2 and 16 pulses m^−2^, and at both 1 ha and 1 km^2^ scales (see Fig 3 for examples, and Figs S10-23 for a full overview), and RMSEs and MEs often reached or exceeded standard deviations within sites (rRMSE of 50-100% or more; Tables S6-7). In a few cases, errors and biases even exceeded cross-site variation by a factor of 2 or more, and some metrics never reached R2_rankswapped_ > 0.9 within sites even in comparisons between high-quality scans, indicating an inherent instability in those metrics (Tables S12).

**Fig 3:**
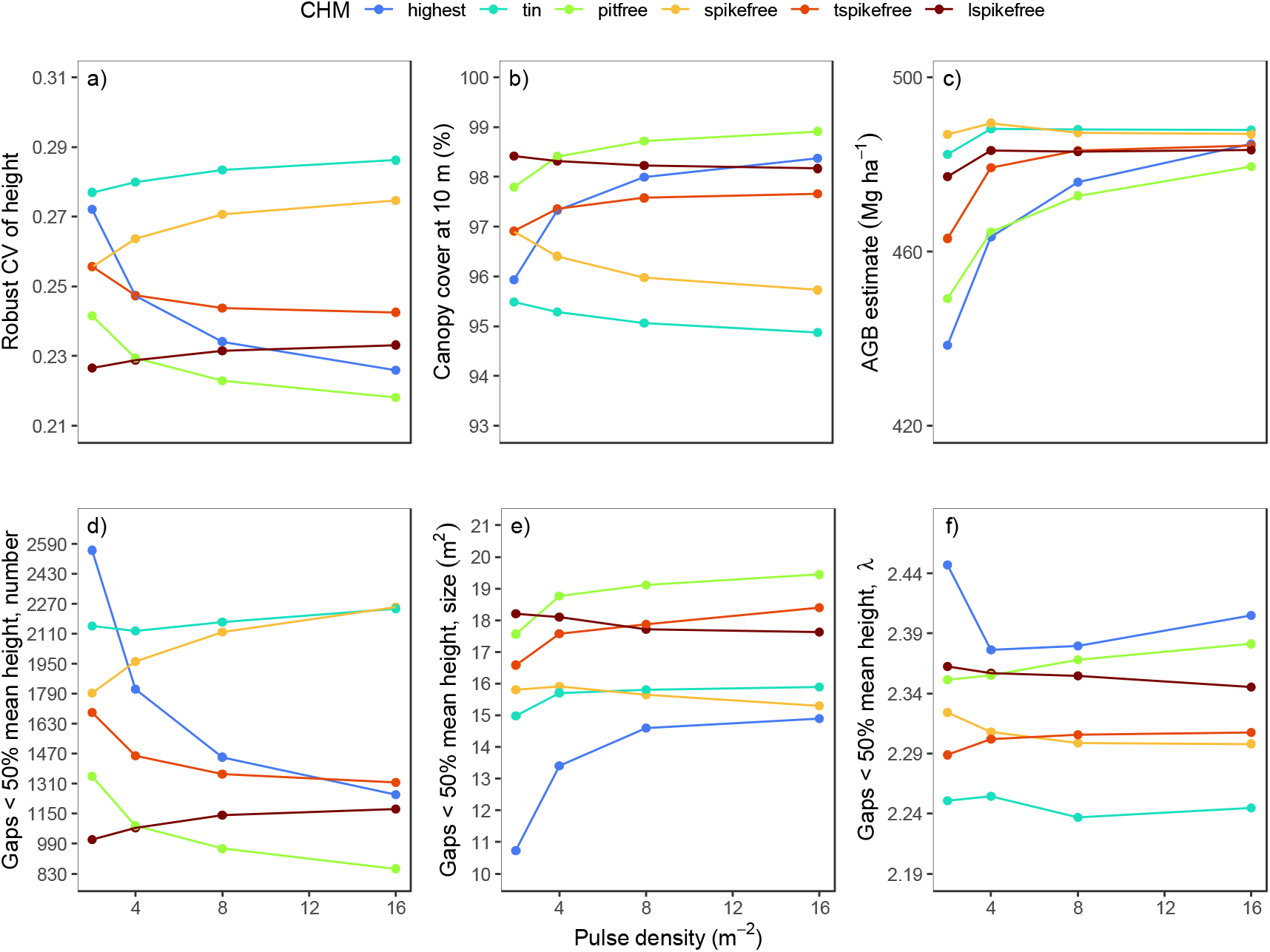
Robustness of CHM-derived structural metrics at Robson Creek under point cloud degradation. Shown are a selection of six measures of forest structure. The top row shows vertical metrics and those derived from them, including (**a**) the robust coefficient of variation of height, (**b**) canopy cover at 10 m aboveground, and (**c**) aboveground biomass (AGB), all estimated at 1 ha resolution. The bottom row shows horizontal metrics calculated at 1 km^2^ resolution, including the number (**d**), average size (**e**) and power law slope (**f**) of gaps, where gaps are defined as clusters >25 m^2^ in size and <50% of mean canopy height. We rescaled tick marks on the y-axis to correspond to 50% of each metric’s typical standard deviation at 16 pulses m^−2^. Biases smaller than inter-tick mark distance thus indicate robustness, while those exceeding it denote instability.

However, there were clear differences between CHM algorithms and between vertical and horizontal metrics. CHM_lspikefree_ and CHM_tin_ were the most robust, with CHM_lspikefree_ performing slightly better. It had low rRMSEs and rMEs for all vertical metrics (< 10% across sites and 10-20% within sites, Tables S6-9) and highly conserved rankings when randomly combining measurements at low and high pulse densities (R2_rankswapped_ ∼1.00 across and R2_rankswapped_ ∼0.90-0.95 within sites, Tables S10-11, Figs. S10-23). The consistency of CHM_lspikefree_-inferred vertical metrics translated to derived products, such as aboveground biomass (AGB, Fig 3, Table S14). Errors due to pulse density degradation (16.11 Mg ha^−1^) were small compared to those from algorithms such as CHM_highest_ and CHM_pitfree_ (49.15 Mg ha^−1^ and 34.24 Mg ha^−1^, respectively), model calibration error (41.94 Mg ha^−1^) and absolute estimates (483.3 Mg ha^−1^). Horizontal metrics were also the most robust when derived from CHM_lspikefree_, but showed considerable errors and uncertainties within sites (50-100% rRMSE and rME; R2_rankswapped_ of 0.3-0.7). An exception were canopy gaps (defined as areas < 50% mean canopy height) extracted at 1 km^2^ resolution. For CHM_lspikefree_, their numbers, mean size and power law slopes were well-preserved across (rRMSEs ∼ 10% and rMEs ∼5%) and within sites (∼25% and 10%).

CHM_tin_-derived metrics were also robust, and for some metrics (e.g., upper percentiles of height), all algorithms performed well. But overall CHM_lspikefree_ was generally more consistent and the only algorithm for which at least some horizontal metrics had within-site rRMSEs below 40% and R2_rankswapped_ > 0.9 (Tables S7 and S11). Two commonly used algorithms, CHM_highest_ and CHM_pitfree_, on the other hand, were consistently the worst-performing, with errors regularly exceeding 100% and rank-consistency < 0.9 for most metrics (Tables S10-11).

## 4. Discussion

### Towards robust forest structure comparisons in time and space

Laser scanning technology is advancing fast, and airborne laser scans (ALS) are increasingly available in huge, open archives (e.g., https://opentopography.org/). This presents a unique opportunity for ecologists to characterize and monitor ecosystems in time (Huertas et al., 2022) and in space (Jucker, 2022). In particular, ALS-derived canopy height models (CHMs, high-resolution grids of canopy height) hold a lot of promise, as they are easy to create, share and interpret and a core part of most point cloud processing tools (Mielcarek et al., 2018; Roussel et al., 2020).

Here, we used high-quality ALS data from nine TERN SuperSites in Australia and systematically assessed the quality of six CHM algorithms across a wide range of ecosystems. We show that estimates of forest structure vary widely both between algorithms and for the same algorithm under different degrees of point cloud degradation. However, robust assessments of canopy structure are possible through careful methodological choices. As a way forward, we present a new type of CHM that increases the robustness of an existing algorithm (Khosravipour et al., 2016), an openly available pipeline to derive it in R, and a set of 16 best-practice guidelines that will allow ecologists to make the most of growing access to ALS data (Fig 4).

**Fig 4:**
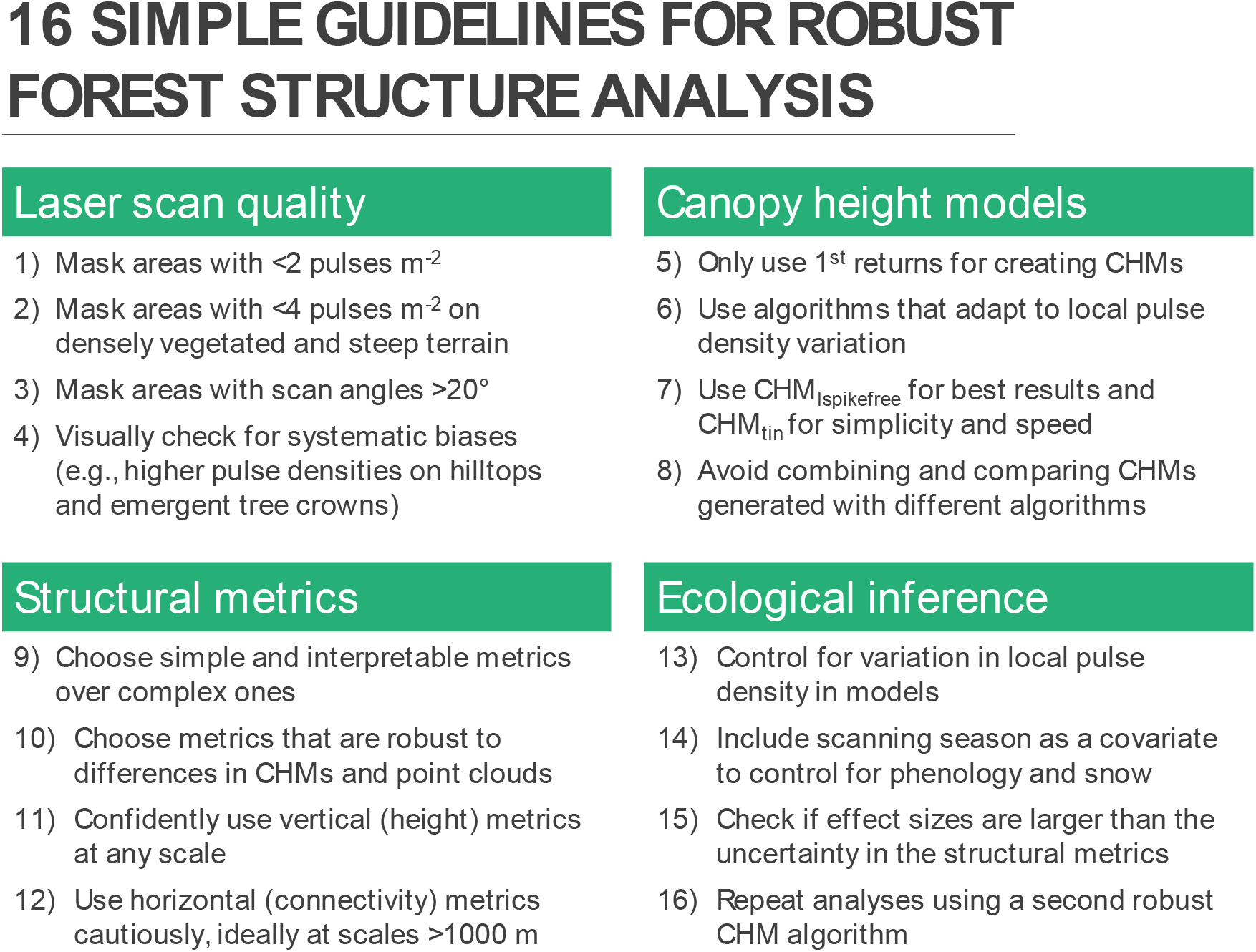
16 simple guidelines for robust forest structure analysis. Shown are best-practice guidelines for robust assessments of forest structure from airborne laser scanning (ALS) via canopy height models (CHMs). We assume that a) scans have been acquired at pulse densities high enough to produce accurate ground models, b) the target resolution for CHMs is 1 m, and c) that the aim is to compare forest structural properties across scans and sites. For some goals, such as tree crown delineation across a single high-quality scan, other algorithms such as CHM_highest_ or CHM_spikefree_ may prove useful.

### Not all canopy height models are created equal…

A clear takeaway from our analysis is that canopy height cannot be compared across CHM algorithms, and that variation in point cloud properties adds large errors. Mean canopy height estimates could differ by up to 10 m between algorithms and up to 60% in relative height, and point cloud degradation could add additional biases of up to ∼5 m in absolute differences, or up to ∼40% in relative terms. This is true even when staying within reasonable quality ranges (e.g., densities ≥ 2 pulses m^−2^) and despite the general robustness of CHMs to differences in light extinction and good retrieval of top canopy height (Mielcarek et al., 2018). To make matters worse, errors were inconsistent both across sites – higher absolute errors in tall, higher relative errors in short forests (Table S5) – and within scans (Fig S5, Table S5). Large scan angle differences (15-20°) also introduced a negative bias in height estimates, but across-scan pulse density variation was the single most important factor in determining CHM quality, in agreement with previous studies (Huertas et al., 2022; LaRue et al., 2022).

Errors and uncertainties in CHM algorithms also had disproportionately large effects on derived structural metrics, often exceeding the ecological variation within sites. For extremely sensitive metrics (e.g., Moran’s I or gap statistics from CHM_highest_), errors and uncertainties due to pulse density even exceeded the variation across the entire gradient from Acacia woodlands (below 10 m) to Mountain Ash forests (up to 90 m). This is beyond most ranges seen in ecological studies, and, barring stand-replacing disturbance, exceeds forest dynamics by orders of magnitude. Even at a single site (Robson Creek) and for a comparatively robust way of quantifying canopy gaps (< 50% mean height and > 25 m^2^ in size), values ranged from ∼800 to ∼2,200 gaps km^−2^ between algorithms, and could go from ∼1,200 to ∼2,600 gaps km^−2^ across pulse densities for a single algorithm (CHM_highest_). In comparison, estimates of aboveground biomass were more stable, but degradation in pulse density could still lead to biases in excess of model calibration errors (∼50 Mg ha^−1^ for CHM_highest_) and >10% of the landscape-level estimates (∼485 Mg ha^−1^).

### … but we can make them more equal

There is, however, a clear way forward when it comes to implementing robust high-resolution (1 m^2^) forest structure assessments (see Fig 4: *16 simple guidelines for robust forest structure analysis* for a condensed version). First, any ALS data should be carefully inspected for potential issues. We recommend masking areas below 2 pulses m^−2^ (averaged at 5-10 m), and below 4 m^−2^ in rugged terrain with dense vegetation (see degradation in Figs 1-2 and Fig S9). The effects of scan angle variation and laser penetration were small relative to those of pulse density (Fig S4), but became noticeable at 15-20°, so we also recommend 20° as an upper limit (see Dayal et al., 2022). Masking of areas is also recommended when DTMs cannot be reliably inferred due to a lack of ground returns.

Second, forest structural estimates should be derived from a single, consistent CHM algorithm, ideally one that uses only first returns and adapts to pulse density variation. Two of our algorithms fulfilled these criteria (CHM_lspikefree_ and CHM_tin_). CHM_lspikefree_, a newly proposed, locally-adaptive variation on the “spikefree” algorithm (Khosravipour et al., 2016) had low errors across metrics (<5% of across-site variation). It also preserved many canopy properties when mixing scans from high and low pulse densities (R2_rankswapped_ = 0.95-1.0 across and within sites) and was the only algorithm that estimated gap statistics with reasonable accuracy (rRMSE ∼25% within sites). CHM_tin_, a much simpler Delaunay-triangulation of first returns, matched the low biases in CHM_lspikefree_, but created porous canopy surfaces and was more susceptible to scan angle differences. Application of CHM_tin_ are thus context-dependent – better for estimating light-penetration, worse for estimating top height – but it should be a useful complement to other robust algorithms. In contrast, two of the most commonly used algorithms, CHM_highest_ and CHM_pitfree_, performed badly throughout. We recommend their application only in specific cases, e.g., when sample areas are scanned with the same instruments and at high densities (> 20 pulses m^−2^).

Third, for most applications we recommend using statistics that summarize the vertical distribution of canopy elements, such as the mean, standard deviation or percentiles of canopy height. Vertical metrics are usually simple to calculate and interpret, robust, and have been successfully used for the modelling and mapping of forest plot attributes (Pearse et al., 2019; Tompalski et al., 2019; Wilkes et al., 2015). In contrast, horizontal metrics that quantify the connectivity of CHM pixels – spatial autocorrelation, height gradients and clusters of canopy features such as gaps – are more problematic. They are attractive, because they open up many ecological questions around canopy dynamics and biodiversity (Asner, Kellner, et al., 2013; Jucker, 2022). However, they are difficult to interpret and proved extremely volatile, with only a single gap type (<50% of mean canopy height) producing moderately reliable statistics, and only for the most robust algorithm at the coarsest scale (CHM_lspikefree_ at 1 km^2^).

Finally, any measurement will involve some form of error. Even the most robust algorithms, such as CHM_lspikefree_ and CHM_tin_, showed small biases in this study, and some features of ALS data (e.g., footprint size, or acquisition timing and its relation to phenology or snow cover) are difficult to correct for. We therefore recommend including summaries of point cloud quality (e.g., pulse density, measured at the resolution of forest structure metrics) and ecological confounders of structure (e.g., vegetation indices that reflect phenology) into statistical models. Crucially, effect sizes should always be put into context of the typical noise and biases of CHMs and their derived metrics. As a rule of thumb, differences < 1.0 m in mean canopy height should be interpreted with great caution. For any type of analysis, a repetition with a second type of CHM may greatly increase the robustness of findings.

### Towards a robust analysis of forest canopies at global scales

There are limitations to our findings. First, we ignored interactions between pulse density degradation and other sources of uncertainty (e.g., an operator flying at higher altitudes can compensate lower point densities by increasing laser power) and focussed on sites with subtle changes in topography (except for Robson Creek; Fig S9). Furthermore, some sources of point cloud variation (rotational patterns of scanners, varying footprint sizes or waveform-to-discrete-point conversion) were beyond the scope of our analysis and would need ray tracing simulations or careful experiments with different instrumentations (Boucher et al., 2023; Brede et al., 2022; Næsset, 2009). Footprint size, for example, has been found to introduce moderate, but highly variable biases into height estimates (0.1-0.5 m), including both positive (Næsset, 2009), negative (Morsdorf et al., 2008; Roussel et al., 2017) and site-dependent (Goodwin et al., 2006) differences between larger and smaller footprints. Overall, even if we knew all error sources and how to correct for them, there would still be practical limitations, as metadata for lidar scans are inconsistently (and often incompletely) reported. In contrast, pulse density is an inherent feature of ALS point clouds, so, together with acquisition timing, it should provide a good first-order correction for forest structure analysis.

Second, if a study relies on a single scan or a set of scans with consistent acquisition parameters, some of our rules-of-thumb could be relaxed. For example, to estimate tree top heights (which we expect to be robust, S10-11), it might be advantageous to retain pulse densities <2 m^−2^ and maximize information by using CHMs based on all returns instead of first returns only (Mielcarek et al., 2018). Similarly, one could use the sharper contours of CHM_highest_ or CHM_spikefree_ for tree crown delineation and apply targeted height corrections (Roussel et al., 2017). In a few cases, it may even make sense to use instable metrics such as rumple indices or gap statistics, as there is an accuracy-robustness trade-off: many metrics are instable precisely because they are sensitive to fine-scale variation in point cloud distributions. For high-quality scans – modern, large-scale surveys easily reach densities of 50-100 pulses m^−2^ (McNicol et al., 2021) – such metrics may provide a lot of ecological information in return for only small biases. However, this comes with a number of caveats: first, most scans found in open archives (e.g., https://opentopography.org/) are neither consistent with each other nor obtained at high pulse densities. Second, even high-quality scans can have highly variable pulse density coverage (cf. biases in Fig S5), and some metrics may be inherently noisy or bias-prone (errors even between 8 and 16 pulses m^−2^; Table S12). Finally, any use of non-robust metrics limits comparability across studies and precludes the investigation of temporal change, unless flightlines and instrumentation are consistent.

Finally, our study does not provide an exhaustive picture of all potential metrics that can be derived from ALS. For example, we might be able to derive more robust alternatives to the horizontal metrics tested here, or develop more stable CHM algorithms. Crucially, our analysis also does not peek below the canopy surface, but this is where a lot of important ecological processes are happening. Expanding our robustness analysis to point cloud or voxel-derived metrics (Pearse et al., 2019; Vincent et al., 2023) is a priority, and the current developments are exciting. Several recent studies systematically assessed how point cloud and instrument properties propagate to the measurement of ecologically interesting features (Dayal et al., 2022; LaRue et al., 2022; Roussel et al., 2017, 2018; Tompalski et al., 2019; Vincent et al., 2023). It is through studies such as these that ecology will soon be ready for global scale analysis of fine-scale forest structure, as well as its impacts on carbon cycles, microclimates, and animal habitats.

## Acknowledgments

We acknowledge TERN and Airborne Research Australia (ARA), who collected the airborne laser scanning data and made them openly available under a CC-BY license, as well as the Traditional Owners and Custodians throughout Australia. TJ was supported by a UK NERC Independent Research Fellowship (grant: NE/S01537X/1) and through a Research Project Grant from the Leverhulme Trust which also funded FJF (grant code: RPG-2020-341).

## Author contributions

FJF and TJu conceived of the study. TJa and GV contributed critically to the analysis and to the writing of the draft. FJF led the development of algorithms, writing and analysis of the data, with substantial inputs from all co-authors.

## Data availability

All data used in this study are openly available on the Terrestrial Ecosystem Research Network’s homepage: https://portal.tern.org.au/metadata/TERN%2F4ff0b4c9-cfa0-4d09-9520-b5402adc583f (ALS data) and https://portal.tern.org.au/metadata/supersite.174 (field data for Robson Creek). All R scripts, including the ALS processing pipeline and functions used for analysis and visualization, as well as derived products that support the results in the article are available on Zenodo (http://doi.org/10.5281/zenodo.10878070).

## Conflict of Interest Statement

The authors declare no conflict of interest.

## Notes

### Competing Interest Statement

The authors have declared no competing interest.

