## Supplementary Materials for "Robust characterization of forest structure from airborne laser scanning – a systematic assessment and sample workflow for ecologists"

### A. Supplementary Text

#### S1: Processing pipeline and CHM analysis

*Overview*

The airborne laser scanning (ALS) processing pipeline is a custom pipeline to clean and process ALS point cloud data. It derives raster products, such as digital terrain models (DTMs), digital surface models (DSMs) and canopy height models (CHMs), and provides ancillary information on pulse densities and scan angles, also in raster format. The final output are summary statistics across the processing workflow as well as metadata (coordinate reference systems, minimum and maximum GPS times, etc.). The pipeline is run from the R environment (R Core Team, 2023), calling LAStools functions (Isenburg, 2023), and has been tested across a wide variety of data sets with minimal tuning. It solely requires the user to specify relevant directories – a data folder, a folder to store temporary files, and a folder for the results – as well as a few basic parameters, such as the resolution for retiling, buffer size, and the number of cores for parallel processing. The pipeline is available on github (~~masked for review~~) and continuously updated. A priority for future development is the inclusion of LAStools-equivalent *lidR* routines (Roussel et al., 2020) as an open-source alternative and aspects of this are being tested (robust and LAStools-equivalanet DTM derivation, in particular).

*Workflow*

The pipeline first searches the data directory for subdirectories that contain point cloud data in a user-defined format (i.e., .las or .laz files). Each of the subdirectories that contains point cloud data is then processed separately. This means that a single command can be used to process many different data sets (e.g., scans from different sites or different acquisition times). Before the actual processing, the outline of all point clouds in a directory is delimited via *LAStools* (lasboundary). If the directory contains scan lines or point clouds that do not intersect with each other, these will be further split into clusters that are processed separately. This accelerates processing, reduces memory demands and avoids errors when lidar scans from separate locations are stored in the same folder. Splitting into data subsets can also be enforced directly by the user to reduce temporary storage demands, but this should only be necessary for very large data (typically >= 100 GB).

After the initial data preparation step, each subset of the data is processed with a standard sequence of commands described in Table 2, each time followed by a reindexing procedure (lasindex). If errors occur, they are captured via *R*'s tryCatch() command and result in the abortion of the processing of the current directory's or cluster's data. The pipeline then jumps to the next directory or point cloud cluster and continue processing normally. Processing can also be restarted. It then targets only directories that have not yet been fully processed (unless recomputing is enforced by the user). Each time, the pipeline generates processing reports with associated timestamps and other point cloud characteristics to assess the overall processing workflow and potential errors.

*Other R packages and routines used as part of the processing pipeline and for analysis*

Apart from external LAStools commands, the pipeline relies mainly on *terra* (Hijmans, 2023), *sf* (Pebesma, 2018; Pebesma & Bivand, 2023), *data.table* (Dowle & Srinivasan, 2023) *lidR* (Roussel et al., 2020), as well as *igraph* (Csárdi et al., 2023), *parallel*, and *Hmisc* (Harrell Jr, 2023).

For some further analysis, we used *futures* (Bengtsson, 2021) and *poweRlaw* (Gillespie, 2015), and for plotting, we used *ggplot2* (Wickham, 2011), *tidyterra* (Hernangómez, 2023), *patchwork* (Pedersen, 2023), *viridis* and *viridisLite* (Garnier et al., 2023).

#### S2: Companion analysis – within-point cloud pulse density variation

*Setup*

Within-scan variation in pulse or point densities is common due a wide range of features, from multiple scan lines that overlap to imperfect tracking of the topographic profile by the airplane (Petras et al., 2023; Roussel et al., 2017). To assess the effect of within-point cloud heterogeneities, we repeated the pulse density thinning applied in the main analysis, but used an approach that kept the original scans' heterogeneities intact. Instead of homogenizing pulse densities within spatial subsamples (e.g., 25 m x 25 m windows), we ordered pulses by their acquisition time and removed every $n^{th}$ pulse, with $n$ chosen to obtain the desired pulse density levels of 16, 8, 4, 2, 1, and 0.5 pulses $m^{-2}$. To assess local heterogeneities, we compared the inhomogeneous and homogenized point clouds at 4 pulses $m^{-2}$. We chose the latter pulse density, as full homogenization of heterogeneous scans can usually only be achieved at pulse densities well below the original sampling density.

*Results*

As expected, within-scan heterogeneities in pulse density  did not lead to systematic bias across scans. However, RMSEs between homogeneous and inhomogeneous 4 pulses $m^{-2}$ were comparable in size to RMSEs between 16 and 2 pulses $m^{-2}$ of homogeneous scans, suggesting large local instabilities in height estimates (Table S5). Furthermore, deviations revealed clear spatial biases, with deviations in canopy height mirroring deviations in pulse density and devatiations still clearly visible even after aggregation at 25 m scale (Figure S5).

#### S3: Companion analysis – laser penetration

*Setup*

A common difference between laser scanners is their power, i.e., the amount of energy that is transmitted from the laser and reflected after hits. Increases in power typically translate into increases in the chances of obtaining higher-order returns and thus improve within-canopy sampling (Brede et al., 2022). To test the effect of laser penetration, we simulated a decrease in return probabilities by removing percentages of higher-order returns from the original TERN point clouds (0%, 25%, 50%, 75%, 100%). As the effects of low power are expected to have stronger effects at low pulse density, we first thinned the point cloud to 4 pulses $m^{-2}$ and then removed a fixed percentage of randomly selected $2^{nd}$ and associated higher-order returns (e.g., 25%). This procedure was repeated for the $3^{rd}$ and all subsequent higher-order returns ($4^{th}$, $5^{th}$ etc.). At 0%, nothing is removed. At 100%, a $1^{st}$ return-only point cloud remains. We note that we would expect no systematic changes in CHM algorithms that rely only on first returns for CHM construction – as these are not affected by the thinning –, but there may still be low levels of noise due to the inherent stochasticity in CHM algorithms.

*Results*

As expected, algorithms that did not rely on first returns ($CHM_{tin}$, $CHM_{lspikefree}$ and $CHM_{tspikefre}$) were trivially unaffected by laser penetration, although there was some background noise (RMSEs < 1 m, Table S5, Figure S7). Of the remaining algorithms, $CHM_{highest}$ showed a small negative bias, reflecting the removal of higher-order returns from the upper canopy. The only strong effects were found in $CHM_{spikefree}$, with positive biases of up to 3 m (Table S5), and in the DTM at Robson Creek, which also had a clear upward bias (Figure S7).

#### S4: Companion analysis – scan angles

*Setup*

Scan angle, the angle at which a laser shot reaches the canopy, depends on the scanning instrument (e.g., maximum angle, rotational patterns) and the distance from the flightline. Canopy far away from the scanner is typically hit at more oblique angles than canopy directly underneath the scanner, changing its transmittance and the likelihood of ground returns (Dayal et al., 2022; Keränen et al., 2016). Variation in scan angles is difficult to simulate from existing point clouds, but all TERN scans consist of multiple, overlapping flight lines. As a result, many areas have been scanned from a different angle. Therefore, to test the robustness of CHM algorithms to differences in scan angle, we reconstructed separate point clouds for individual flightlines using the "lassplit" function in LAStools. We then extracted areas that were scanned by overlapping flightlines and compared the CHMs of the two flightlines with each other. For each pixel covered by two flightlines, we determined the absolute scan angle in each and took the flightline with the lower absolute scan angle as reference. To control for variation in pulse density, we only kept overlapping areas with the same pulse densities in both flightlines.

*Results*

Scan angle had clear directional effects on canopy height estimation, with CHMs uniformly, but weakly decreasing when scan angle differences were between 15 and 20 degrees (Figure S8, Table S5). Differences were the largest for tall canopies, but still small in comparison to pulse density effects (average bias around -1 m for Warra, compared to biases of up to -5 m with pulse density). $CHM_{lspikefree}$ was generally among the best algorithms with regard to scan angles, but still showed the same negative bias as all other CHM algorithms.

#### S5: AGB estimation at Robson Creek

To estimate above ground biomass (AGB, Mg $ha^{-1}$) at Robson Creek, we downloaded data for 25 1-ha-plots from TERN Australia (https://portal.tern.org.au/metadata/supersite.174). We obtained data both at plot and tree level. Since we discovered that some of the provided plot coordinates were not aligned with the plot grid, we recalculated them: we used the minimum and maximum extent of tree-level coordinates to define plot extents and linked the resulting quadrats with provided AGB estimates at 1 ha scale. We then developed reference AGB models following standard protocols (Labriere et al., 2018).

For all six algorithms, we selected the CHM from the highest-quality scan with homogeneous pulse density as reference (16 pulses $m^{-2}$). For each of the six CHMs, we then predicted AGB from the median canopy height ($H_{50}$) of the 25 plots by fitting a simple power law model: AGB = A x $H_{50}^b$, where A and b are inferred coefficients (for details cf. Labriere et al., 2018). For each CHM algorithm, we used its specific AGB model to predict AGB at different (homogeneous) pulse density levels. Details on the results from model fitting can be found in Table S14. Note that this is a best-case scenario, as we use locally fitted and algorithm-specific AGB models. One could also imagine a scenario where an AGB model is calibrated with a CHM derived from one algorithm and then applied to a CHM calculated with a different algorithm. For some algorithm combinations (e.g. $CHM_{tin}$ and $CHM_{highest}$), this could result in massive biases, beyond anything shown in Table S14.

### B. Tables

| Step | Functions (LAStool/custom) | Key parameters and flags | Short summary |
|---|---|---|---|
| splitting into clusters | lasboundary | number of forced clusters | clusters are delineated automatically, so enforcement is only recommended in special cases |
| retiling | lastile | tile size<br>buffer size | typically, tiles of 250-500 m and buffers of 25-50 m |
| duplicate removal | lasduplicate | unique_xyz | only clear duplicates (same x, y, z coordinates) are removed, should remove artefacts from previous tiling if present |
| denoising | lasnoise | step size<br>isolation distance | typically, a step size of 3, with isolation parameter automatically chosen based on mean point density |
| ground classification | lasground_new | step size | default parameters are kept (25 m initial search window, no specific refinement options picked) |
| height cutoff | lasheight | maximum height | removal of any points above maximum height threshold (typically 125m) to cut off high noise from clouds |
| pulse density map | lasgrid | step size | based on number of first returns per grid cell |
| scan angle map | lasgrid | step size | based on absolute scan angle |
| pulse density masks | custom | step size<br>pulse density levels | aggregation into 5 x 5 $m^2$ tiles<br>levels of 0.5 pulses $m^{-2}$ and 2.0 pulses $m^{-2}$ |
| DTM/DSM/CHM derivation | las2dem<br>custom functions | step size | creation of DTMs, DSMs and CHMs as described in main text (all except $CHM_{pitfree}$) |
| height cutoff | lasheight | | normalization of point cloud |
| $CHM_{pitfree}$ derivation | las2dem<br>custom functions | step size | creation of $CHM_{pitfree}$ |

**Table S1: ALS point cloud processing workflow.** Steps of the point cloud processing workflow according to the order in which they are executed. All processing is carried out with LAStools, some smaller intermediate steps, such as reindexing of tiles with *lasindex* or calculation of tile-level summary statistics via *lasinfo* have been omitted.

| Algorithm | Abbreviation | Description |
|---|---|---|
| First-return TIN | $DSM/CHM_{tin}$ | Delaunay triangulation over ALS first returns, as implemented in the *LAStools* blast2dem function. Performed on the non-normalized point cloud to create a digital surface model (DSM). Gridded at 1 m resolution and normalized via DTM-subtraction. |
| Highest return | $DSM/CHM_{highest}$ | Gridding of highest returns at 1 m, using lasgrid, and CHM creation via DTM subtraction. |
| Pitfree algorithm | $DSM/CHM_{pitfree}$ | Pitfree algorithm (Khosravipour et al., 2014), as described in *LAStools* batch procedures (https://github.com/LAStools/LAStools/blob/master/example_batch_scripts, accessed on 7 February 2023). Moves through the normalized point cloud vertically and creates intermediate CHMs that combine for a final "pitfree" CHM at 1 m resolution. Carried out with standard parameters, i.e., thinning at half the target resolution (0.5 m), "splatting out" of returns (radius = 0.05 m) and removal of Delaunay triangles in intermediate CHMs with edge lengths larger than 3 times the target resolution (3 m). Intermediate CHMs were computed at 5 m vertical steps. A DSM was obtained by summing DTM and CHM. |
| Spikefree algorithm | $DSM/CHM_{spikefree}$ | Constrained Delaunay triangulation (Khosravipour et al., 2016). Triangulates points that are nearby both horizontally and vertically and omits non-relevant returns below already triangulated areas. Conceived as update to the pitfree algorithm and implemented on non-normalized point clouds via las2dem. The so-called "freeze constraint" parameter is set to three times the average pulse spacing, i.e., sqrt(1/pulse density). Note that this is not the original implementation, but a common heuristic described in the *LAStools* documentation (https://rapidlasso.com/2016/02/03/generating-spike-free-digital-surface-models-from-lidar/, accessed on 7 February 2023). Used to create a DSM from non-normalized returns, then converted to CHM via DTM-subtraction at 1 m resolution. |
| Spikefree algorithm, (thinned point cloud) | $DSM/CHM_{tspikefree}$ | Same as spikefree algorithm, but based on a systematically thinned and homogenized point cloud (2 points per $m^2$). Thinning is performed with the lasthin function at 1 m resolution, retaining only first returns to increase robustness across instruments. The freeze constraint is set to 3 x sqrt(1/2) m or ~ 2.1 m. |
| Spikefree algorithm (locally adaptive) | $DSM/CHM_{lspikefree}$ | Same as the spikefree algorithm, but using only first returns and a freeze-constraint adapted to local pulse densities. The algorithm first computes pulse densities across 5 x 5 $m^2$ patches and divides the point cloud into pulse density classes, in steps of 0.5 up to 3 pulses $m^{-2}$, in steps of 1 from 3 to 10 pulses $m^{-2}$, 2 to 20 pulses $m^{-2}$, 5 to 50 pulses $m^{-2}$, 25 up to 100 pulses $m^{-2}$. Point cloud areas with pulse densities > 100 $m^{-2}$ are processed as if their pulse density was 100 $m^{-2}$. For each pulse density class, we create a buffer zone of 5 x 5 $m^2$ and compute a DSM with a pulse-density adapted freeze constraint. The freeze constraint is determined as: $freeze = m_{freeze}/(\ln(s_{freeze} \times pd + o_{freeze}) + 1.0)$, where *pd* is the pulse density of the current patch, and $m_{freeze}$, $s_{freeze}$ and $o_{freeze}$ are free parameters (multiplicator, slope and offset, respectively). The formula was developed empirically and calibrated through a grid search of parameters on three 1 km x 1 km canopy extents at Alice Mulga, Robson Creek and Watts Creek. A robust parameter set was found to be $m_{freeze}$ = 3.0, $s_{freeze}$ = 1.75 and $o_{freeze}$ = 2.1, which we use here. |

**Table S2: Canopy height model algorithms.** Description of the six CHM-derivation algorithms tested in the study.

| Type of metric | Metric | Calculation |
|---|---|---|
| Vertical (not considering between-pixel spatial connections) | Mean height (m) | Mean canopy height |
| | Median of height (m) | Median canopy height |
| | 25th perc. of height (m) | 25th percentile of canopy height |
| | 75th perc. of height (m) | 75th percentile of canopy height |
| | 95th perc. of height (m) | 95th percentile of canopy height |
| | 99th perc. of height (m) | 99th percentile of canopy height |
| | SD of height (m) | Standard deviation of canopy height |
| | CV of height | Coefficient of variation of canopy height (SD / mean) |
| | Robust CV of height | Robust coefficient of variation of canopy height (Lobry et al., 2023) |
| | IQR of height (m) | Interquartile range of canopy height |
| | Canopy cover at 2 m (%) | Percentage of pixels >= 2 m in canopy height |
| | Canopy cover at 10 m (%) | Percentage of pixels >= 10 m in canopy height |
| Horizontal (connectivity of pixels with their neighbours) | Rumple index | Surface area divided by planar area |
| | Normalized rumple index | Rumple index calculated for a surface that has been normalized by maximum canopy height (99th percentile) |
| | Moran's I | Spatial autocorrelation as measured by Moran's I (3 x 3 raster neighbourhood) |
| | clusters < 25th perc., number | Number of clusters below the 25th percentile of canopy height (and area >= 25 $m^2$); a type of "gaps" |
| | clusters < 25th perc., size ($m^2$) | Area of clusters below the 25th percentile of canopy height (and area >= 25 $m^2$); a type of "gaps" |
| | clusters > 75th perc., number | Number of clusters above the 75th percentile of canopy height (and area >= 25 $m^2$); a type of "crown cluster" |
| | clusters > 75th perc., size ($m^2$) | Area of clusters above the 75th percentile of canopy height (and area >= 25 $m^2$); a type of "crown cluster" |
| | gaps < 10 m, number | Number of absolute gaps; defined with regard to an absolute height threshold (10 m, and area >= 25 $m^2$) |
| | gaps < 10 m, size ($m^2$) | Area of absolute gaps |
| | gaps < 10 m, lambda | Power law slope of the size distribution of absolute gaps, using poweRlaw package and xmin = 25 $m^2$ (Gillespie, 2015) |
| | gaps < 50% mean height, number | Number of relative gaps, defined with regard to a relative height threshold (50% of mean height, and area >= 25 $m^2$) |
| | gaps < 50% mean height, size ($m^2$) | Area of relative gaps |
| | gaps < 50% mean height, lambda | Power law slope of the size distribution of relative gaps, using poweRlaw package and xmin = 25 $m^2$ (Gillespie, 2015) |

**Table S3: Forest structure metrics tested in the study.** Shown are all 25 metrics tested in this study. Aggregation extents are 100 and 1000 m. We note that we use the power law slope of gap size frequency distributions as an indicator of forest structure, but we did not test whether gap patterns actually conformed to power laws.

| Model | Alice Mulga | Chowilla | Credo | Litchfield | Rushworth | Robson Creek | Zigzag Creek | Warra | Watts Creek |
|---|---|---|---|---|---|---|---|---|---|
| highest | 3.43 | 2.52 | 3.2 | 7.29 | 10.06 | 27.33 | 19.05 | 29.3 | 39.73 |
| tin | 1.73 | 1.17 | 1.85 | 2.9 | 5.78 | 23.74 | 12.02 | 20.18 | 28.88 |
| pitfree | 2.72 | 1.94 | 2.78 | 6.59 | 9.66 | 26.92 | 18.96 | 29.19 | 39.83 |
| spikefree | 2.09 | 1.42 | 2.19 | 4.21 | 7.17 | 25 | 14.75 | 24.6 | 33.71 |
| tspikefree | 2.39 | 1.62 | 2.38 | 4.58 | 7.98 | 25.78 | 16.32 | 26.13 | 36.19 |
| lspikefree | 2.74 | 1.9 | 2.6 | 5.94 | 8.88 | 26.3 | 17.88 | 28.15 | 38.44 |
| DTM | 601 | 50 | 445 | 217 | 202 | 856 | 277 | 246 | 988 |

**Table S4: Mean canopy height and mean elevation by algorithm and site.** Shown are reference heights for all nine SuperSites and all CHM algorithms, based on point clouds with 16 pulses $m^{-2}$. Units are metres throughout.

| **RMSE/ME (m)** at 1 $m^2$ pixel level | **Model** | **Alice Mulga** RMSE / ME (m) | **Chowilla** RMSE / ME (m) | **Credo** RMSE / ME (m) | **Litchfield** RMSE / ME (m) | **Rushworth** RMSE / ME (m) | **Robson Creek** RMSE / ME (m) | **Zigzag Creek** RMSE / ME (m) | **Warra** RMSE / ME (m) | **Watts Creek** RMSE / ME (m) |
|---|---|---|---|---|---|---|---|---|---|---|
| **Deviations at low pulse density** | highest | 1.71 / -0.99 | 1.58 / -0.79 | 2.21 / -0.76 | 5.13 / -2.39 | 4.70 / -2.25 | 4.33 / -1.91 | 7.49 / -3.72 | 9.48 / -4.33 | 13.93 / -5.64 |
| | tin | 1.20 / 0.04 | 1.09 / 0.05 | 1.78 / 0.05 | 3.75 / 0.22 | 4.03 / 0.26 | 4.20 / 0.16 | 6.75 / 0.35 | 9.13 / 0.33 | 13.23 / 0.57 |
| **(2 $m^{-2}$ vs. 16 $m^{-2}$)** | pitfree | 1.79 / -1.15 | 1.39 / -0.70 | 1.49 / -0.44 | 4.08 / -1.72 | 3.35 / -1.39 | 2.66 / -1.04 | 5.10 / -2.24 | 5.95 / -2.35 | 9.42 / -3.17 |
| | spikefree | 1.24 / 0.10 | 1.13 / 0.08 | 1.58 / 0.07 | 3.64 / 0.19 | 3.57 / 0.39 | 3.04 / 0.44 | 5.54 / 0.85 | 6.81 / 0.71 | 10.31 / 1.29 |
| | tspikefree | 1.15 / -0.24 | 1.04 / -0.15 | 1.51 / -0.15 | 3.81 / -0.54 | 3.45 / -0.66 | 2.78 / -0.58 | 5.53 / -1.49 | 7.28 / -2.06 | 10.58 / -2.38 |
| | lspikefree | 0.93 / -0.03 | 0.88 / -0.03 | 1.17 / -0.03 | 3.19 / -0.41 | 2.58 / -0.03 | 1.98 / 0.10 | 3.76 / -0.16 | 4.52 / -0.52 | 7.13 / -0.20 |
| | DTM | 0.02 / 0.00 | 0.02 / 0.00 | 0.02 / 0.00 | 0.03 / 0.01 | 0.03 / 0.00 | 1.35 / 0.28 | 0.10 / 0.01 | 0.56 / 0.16 | 0.29 / 0.10 |
| **Deviations due to within-site pulse density variation** | highest | 1.09 / 0.05 | 1.03 / 0.04 | 1.41 / 0.05 | 3.51 / 0.15 | 2.98 / 0.21 | 2.76 / 0.07 | 4.77 / 0.35 | 6.41 / 0.21 | 9.36 / 0.56 |
| | tin | 1.15 / -0.01 | 1.04 / -0.01 | 1.70 / 0.00 | 3.60 / -0.02 | 3.84 / -0.01 | 4.06 / 0.01 | 6.47 / 0.00 | 8.86 / -0.02 | 12.63 / 0.06 |
| **(heterogeneous vs. homogeneous, at 4 $m^{-2}$)** | pitfree | 1.16 / 0.07 | 0.95 / 0.03 | 1.04 / 0.02 | 2.76 / 0.09 | 2.24 / 0.11 | 1.93 / 0.02 | 3.31 / 0.17 | 3.88 / 0.06 | 6.10 / 0.23 |
| | spikefree | 1.04 / 0.06 | 0.93 / 0.04 | 1.29 / 0.06 | 3.09 / 0.11 | 2.87 / 0.19 | 2.56 / 0.02 | 4.50 / 0.21 | 5.95 / 0.14 | 8.52 / 0.24 |
| | tspikefree | 1.03 / 0.02 | 0.95 / 0.01 | 1.33 / 0.02 | 3.49 / 0.06 | 3.01 / 0.09 | 2.34 / 0.03 | 4.67 / 0.22 | 6.11 / 0.15 | 8.85 / 0.38 |
| | lspikefree | 0.82 / 0.02 | 0.76 / 0.01 | 0.99 / 0.01 | 2.73 / 0.08 | 2.17 / 0.05 | 1.70 / 0.02 | 3.20 / 0.10 | 3.86 / 0.09 | 5.95 / 0.15 |
| | DTM | 0.02 / -0.00 | 0.02 / -0.00 | 0.02 / -0.00 | 0.03 / -0.00 | 0.02 / -0.00 | 0.77 / 0.03 | 0.08 / -0.00 | 0.43 / 0.01 | 0.23 / -0.00 |
| **Deviations at large scan angles** | highest | 0.64 / -0.02 | 0.68 / -0.01 | 0.63 / 0.01 | 2.73 / -0.02 | 2.03 / -0.05 | 2.81 / 0.08 | 4.05 / -0.19 | 8.11 / -0.22 | 10.05 / -0.81 |
| | tin | 1.09 / -0.08 | 0.94 / -0.09 | 1.08 / -0.09 | 3.58 , / -0.54 | 3.95 / -0.66 | 3.66 / -0.11 | 7.36 / -1.05 | 10.30 / -0.42 | 16.60 / -0.94 |
| **(> 15° difference)** | pitfree | 0.65 / -0.03 | 0.61 / -0.02 | 0.48 / 0.00 | 2.27 / -0.03 | 1.65 / -0.05 | 2.99 / -0.17 | 3.06 / -0.19 | 6.07 / -0.53 | 8.09 / -1.02 |
| | spikefree | 0.77 / -0.03 | 0.68 / -0.02 | 0.65 / -0.01 | 2.69 / -0.10 | 2.46 / -0.12 | 2.19 / -0.10 | 5.00 / -0.36 | 7.18 / -0.58 | 12.34 / -0.54 |
| | tspikefree | 0.87 / -0.02 | 0.81 / -0.04 | 0.79 / -0.02 | 3.26 / -0.28 | 2.78 / -0.21 | 2.52 / -0.04 | 5.03 / -0.57 | 7.64 / -0.61 | 11.55 / -1.24 |
| | lspikefree | 0.63 / -0.01 | 0.58 / -0.01 | 0.47 / 0.00 | 2.45 / -0.09 | 1.81 / -0.05 | 1.77 / -0.04 | 3.52 / -0.29 | 5.17 / -0.43 | 8.78 / -1.12 |
| | DTM | 0.07 / 0.01 | 0.03 / 0.00 | 0.03 / 0.01 | 0.06 / 0.01 | 0.03 / 0.01 | 4.02 / 1.54 | 0.09 / 0.01 | 1.27 / 0.36 | 0.39 / 0.05 |
| **Deviations at low laser penetration** | highest | 0.23 / -0.02 | 0.13 / -0.01 | 0.26 / -0.01 | 0.77 / -0.07 | 0.79 / -0.08 | 0.71 / -0.06 | 1.58 / -0.21 | 1.62 / -0.16 | 3.22 / -0.35 |
| | tin | 0.00 / 0.00 | 0.00 / 0.00 | 0.01 / 0.00 | 0.06 / 0.00 | 0.04 / 0.00 | 0.99 / 0.08 | 0.21 / 0.00 | 0.73 / 0.03 | 1.47 / 0.09 |
| **(no higher-order returns)** | pitfree | 0.70 / 0.18 | 0.50 / 0.11 | 0.42 / 0.05 | 0.94 / 0.08 | 0.90 / 0.11 | 1.99 / 0.10 | 1.54 / 0.09 | 1.64 / 0.02 | 3.77 / -0.11 |
| | spikefree | 1.03 / 0.46 | 0.71 / 0.21 | 0.92 / 0.17 | 2.21 / 0.59 | 3.16 / 1.28 | 2.26 / 0.76 | 5.83 / 2.91 | 4.30 / 1.36 | 9.45 / 3.63 |
| | tspikefree | 0.23 / 0.00 | 0.28 / 0.00 | 0.74 / 0.00 | 2.76 / 0.00 | 2.13 / 0.00 | 1.75 / 0.01 | 3.57 / 0.00 | 4.69 / 0.01 | 6.68 / 0.01 |
| | lspikefree | 0.00 / 0.00 | 0.00 / 0.00 | 0.02 / 0.00 | 0.10 / 0.00 | 0.03 / 0.00 | 0.13 / 0.00 | 0.12 / 0.00 | 0.25 / 0.00 | 0.85 / 0.00 |
| | DTM | 0.02 / 0.00 | 0.02 / 0.00 | 0.02 / 0.00 | 0.03 / 0.01 | 0.03 / 0.00 | 6.27 / 3.95 | 0.18 / 0.02 | 1.00 / 0.31 | 0.79 / 0.26 |

**Table S5: Height deviations from reference models under point cloud degeneration.** Shown are root mean squared error (RMSE) and mean error (or bias, ME) for digital terrain and surface models (DTMs, DSMs) under four scenarios: low pulse densities (2 vs. 16 pulses $m^{-2}$), heterogeneous pulse densities (vs. inhomogeneous, both 4 pulses $m^{-2}$), large scan angle deviations (>= 15° from reference) and low laser penetration (no vs. all higher-order returns, at 4 pulses $m^{-2}$). All effects are in m. Combinations of sites/algorithms with absolute biases >= 1.0 m are shown in red.

| **rRMSE** across sites | **Metric** | **$CHM_{highest}$** | | **$CHM_{tin}$** | | **$CHM_{pitfree}$** | | **$CHM_{spikefree}$** | | **$CHM_{tspikefree}$** | | **$CHM_{lspikefree}$** | |
|---|---|---|---|---|---|---|---|---|---|---|---|---|---|
| | | **100 m** | **1000 m** | **100 m** | **1000 m** | **100 m** | **1000 m** | **100 m** | **1000 m** | **100 m** | **1000 m** | **100 m** | **1000 m** |
| **Vertical** | Mean height (m) | 0.21 | 0.21 | 0.03 | 0.03 | 0.14 | 0.14 | 0.05 | 0.05 | 0.11 | 0.11 | 0.03 | 0.03 |
| | Median of height (m) | 0.25 | 0.25 | 0.05 | 0.05 | 0.18 | 0.17 | 0.06 | 0.07 | 0.15 | 0.16 | 0.05 | 0.04 |
| | 25th perc. of height (m) | 0.50 | 0.47 | 0.09 | 0.08 | 0.30 | 0.26 | 0.17 | 0.16 | 0.20 | 0.18 | 0.06 | 0.05 |
| | 75th perc. of height (m) | 0.14 | 0.13 | 0.03 | 0.02 | 0.12 | 0.11 | 0.03 | 0.02 | 0.08 | 0.09 | 0.04 | 0.04 |
| | 95th perc. of height (m) | 0.06 | 0.07 | 0.02 | 0.02 | 0.05 | 0.05 | 0.02 | 0.01 | 0.04 | 0.05 | 0.02 | 0.02 |
| | 99th perc. of height (m) | 0.05 | 0.05 | 0.02 | 0.02 | 0.05 | 0.04 | 0.02 | 0.01 | 0.03 | 0.03 | 0.03 | 0.02 |
| | SD of height (m) | 0.27 | 0.16 | 0.06 | 0.05 | 0.18 | 0.11 | 0.10 | 0.07 | 0.09 | 0.05 | 0.06 | 0.05 |
| | CV of height | 0.60 | 0.65 | 0.19 | 0.14 | 0.58 | 0.52 | 0.15 | 0.13 | 0.17 | 0.09 | 0.15 | 0.04 |
| | Robust CV of height | 0.55 | 0.56 | 0.11 | 0.11 | 0.36 | 0.38 | 0.14 | 0.14 | 0.12 | 0.11 | 0.04 | 0.04 |
| | IQR of height (m) | 0.63 | 0.40 | 0.09 | 0.07 | 0.44 | 0.27 | 0.15 | 0.12 | 0.21 | 0.14 | 0.10 | 0.07 |
| | Canopy cover at 2 m (%) | 0.50 | 0.51 | 0.09 | 0.08 | 0.45 | 0.46 | 0.11 | 0.11 | 0.09 | 0.09 | 0.05 | 0.04 |
| | Canopy cover at 10 m (%) | 0.22 | 0.22 | 0.03 | 0.03 | 0.15 | 0.15 | 0.06 | 0.06 | 0.10 | 0.10 | 0.04 | 0.03 |
| **Horizontal** | Rumple index | 1.45 | 1.51 | 0.15 | 0.16 | 0.60 | 0.62 | 0.44 | 0.46 | 0.46 | 0.46 | 0.15 | 0.15 |
| | Normalized rumple index | 2.17 | 2.02 | 0.41 | 0.42 | 0.76 | 0.72 | 0.81 | 0.79 | 0.36 | 0.29 | 0.37 | 0.36 |
| | Moran's I | 2.08 | 2.45 | 0.37 | 0.44 | 1.18 | 1.17 | 0.88 | 1.01 | 0.52 | 0.55 | 0.43 | 0.42 |
| | clusters < $25^{th}$ perc., number | 1.87 | 1.91 | 0.47 | 0.41 | 0.83 | 0.75 | 0.50 | 0.44 | 0.43 | 0.34 | 0.42 | 0.36 |
| | clusters < $25^{th}$ perc., size ($m^2$) | 1.17 | 1.26 | 0.72 | 0.83 | 3.45 | 1.15 | 0.80 | 0.47 | 0.77 | 0.52 | 0.89 | 0.31 |
| | clusters > $75^{th}$ perc., number | 1.40 | 1.62 | 0.42 | 0.36 | 1.46 | 1.82 | 0.67 | 0.67 | 0.49 | 0.45 | 0.41 | 0.37 |
| | clusters > $75^{th}$ perc., size ($m^2$) | 0.92 | 0.96 | 0.61 | 0.48 | 0.69 | 0.53 | 0.94 | 1.14 | 0.64 | 0.61 | 0.61 | 0.31 |
| | gaps < 10 m, number | 1.71 | 1.78 | 0.22 | 0.09 | 0.90 | 0.84 | 0.31 | 0.22 | 0.53 | 0.46 | 0.26 | 0.08 |
| | gaps < 10 m, size ($m^2$) | 0.56 | 0.37 | 0.48 | 0.08 | 0.43 | 0.15 | 0.45 | 0.17 | 0.50 | 0.17 | 0.40 | 0.15 |
| | gaps < 10 m, lambda | | 0.30 | | 0.28 | | 0.28 | | 0.31 | | 0.29 | | 0.16 |
| | gaps < 50% mean height, number | 2.17 | 2.45 | 0.28 | 0.12 | 0.93 | 0.92 | 0.35 | 0.23 | 0.60 | 0.53 | 0.33 | 0.09 |
| | gaps < 50% mean height, size ($m^2$) | 0.74 | 0.75 | 0.64 | 0.12 | 0.64 | 0.34 | 0.66 | 0.15 | 0.57 | 0.21 | 0.55 | 0.13 |
| | gaps < 50% mean height, lambda | | 0.34 | | 0.16 | | 0.27 | | 0.21 | | 0.19 | | 0.12 |

**Table S6: Relative RMSE across sites for forest structure metrics (100 / 1000 m).** Shown is the RMSE (2 vs. 16 pulses $m^2$) of key forest structure metrics across all nine TERN sites, separated by algorithm and normalized by the standard deviation across the nine sites.  Minimum size of gaps and clusters is 25 $m^2$. Combinations of metrics and algorithms with a lot of noise (rRMSEs >= 20%) are shown in red.

| **rRMSE** **within sites** | **Metric** | **$CHM_{highest}$** | | **$CHM_{tin}$** | | **$CHM_{pitfree}$** | | **$CHM_{spikefree}$** | | **$CHM_{tspikefree}$** | | **$CHM_{lspikefree}$** | |
|---|---|---|---|---|---|---|---|---|---|---|---|---|---|
| | | **100 m** | **1000 m** | **100 m** | **1000 m** | **100 m** | **1000 m** | **100 m** | **1000 m** | **100 m** | **1000 m** | **100 m** | **1000 m** |
| **Vertical** | Mean height (m) | 1.05 | 1.83 | 0.15 | 0.22 | 0.90 | 1.57 | 0.20 | 0.32 | 0.37 | 0.65 | 0.10 | 0.14 |
| | Median of height (m) | 1.16 | 1.98 | 0.57 | 1.55 | 0.92 | 1.64 | 0.54 | 1.37 | 0.42 | 0.61 | 0.15 | 0.21 |
| | 25th perc. of height (m) | 1.36 | 2.26 | 1.50 | 3.86 | 1.00 | 1.58 | 1.82 | 24.54 | 0.44 | 0.74 | 0.35 | 0.47 |
| | 75th perc. of height (m) | 0.68 | 1.16 | 0.13 | 0.18 | 0.85 | 1.39 | 0.09 | 0.12 | 0.33 | 0.58 | 0.13 | 0.20 |
| | 95th perc. of height (m) | 0.36 | 0.74 | 0.11 | 0.17 | 0.49 | 0.84 | 0.07 | 0.08 | 0.20 | 0.41 | 0.10 | 0.18 |
| | 99th perc. of height (m) | 0.28 | 0.60 | 0.12 | 0.18 | 0.37 | 0.68 | 0.08 | 0.10 | 0.16 | 0.31 | 0.10 | 0.17 |
| | SD of height (m) | 0.48 | 0.73 | 0.21 | 0.35 | 0.59 | 0.92 | 0.21 | 0.34 | 0.26 | 0.42 | 0.16 | 0.28 |
| | CV of height | 1.55 | 2.94 | 0.36 | 0.69 | 1.25 | 2.35 | 0.37 | 0.76 | 0.32 | 0.52 | 0.14 | 0.18 |
| | Robust CV of height | 1.49 | 2.58 | 0.43 | 0.75 | 1.09 | 2.03 | 0.48 | 0.83 | 0.32 | 0.53 | 0.12 | 0.18 |
| | IQR of height (m) | 0.96 | 1.36 | 0.25 | 0.34 | 0.91 | 1.43 | 0.27 | 0.32 | 0.37 | 0.57 | 0.20 | 0.34 |
| | Canopy cover at 2 m (%) | 1.54 | 3.79 | 0.29 | 0.58 | 1.27 | 2.76 | 0.40 | 0.87 | 0.27 | 0.55 | 0.15 | 0.29 |
| | Canopy cover at 10 m (%) | 0.89 | 1.71 | 0.18 | 0.23 | 0.63 | 1.16 | 0.24 | 0.39 | 0.38 | 0.62 | 0.15 | 0.21 |
| **Horizontal** | Rumple index | 3.41 | 5.92 | 0.45 | 0.78 | 1.09 | 1.87 | 1.19 | 2.05 | 0.67 | 1.04 | 0.60 | 1.01 |
| | Normalized rumple index | 5.03 | 8.46 | 0.73 | 1.24 | 1.79 | 3.34 | 1.55 | 2.75 | 0.74 | 1.14 | 0.82 | 1.46 |
| | Moran's I | 3.30 | 6.20 | 0.64 | 1.21 | 1.68 | 2.98 | 1.52 | 3.06 | 0.82 | 1.56 | 0.71 | 1.26 |
| | clusters < $25^{th}$ perc., number | 4.30 | 10.40 | 0.75 | 1.16 | 1.61 | 3.06 | 0.94 | 1.24 | 1.06 | 1.55 | 0.79 | 1.00 |
| | clusters < $25^{th}$ perc., size ($m^2$) | 1.64 | 2.96 | 0.79 | 1.09 | 3.37 | 4.15 | 1.17 | 1.44 | 0.99 | 1.03 | 1.00 | 1.08 |
| | clusters > $75^{th}$ perc., number | 1.95 | 5.57 | 0.60 | 0.83 | 1.36 | 2.19 | 0.97 | 2.15 | 0.77 | 1.51 | 0.62 | 1.10 |
| | clusters > $75^{th}$ perc., size ($m^2$) | 1.22 | 2.61 | 0.79 | 0.95 | 0.91 | 1.08 | 1.47 | 2.96 | 0.78 | 1.09 | 0.87 | 1.17 |
| | gaps < 10 m, number | 3.31 | 15.60 | 0.53 | 0.37 | 1.30 | 2.02 | 0.60 | 0.74 | 0.88 | 1.32 | 0.52 | 0.30 |
| | gaps < 10 m, size ($m^2$) | 7.98 | 12.20 | 4.79 | 11.37 | 0.80 | 0.75 | 0.62 | 0.43 | 0.76 | 0.58 | 0.60 | 0.38 |
| | gaps < 10 m, lambda | | 1.00 | | 0.44 | | 0.78 | | 0.65 | | 0.75 | | 0.40 |
| | gaps < 50% mean height, number | 3.12 | 6.30 | 0.51 | 0.42 | 1.45 | 2.68 | 0.56 | 0.55 | 0.88 | 1.25 | 0.52 | 0.26 |
| | gaps < 50% mean height, size ($m^2$) | 0.74 | 1.18 | 0.64 | 0.56 | 0.89 | 0.95 | 0.64 | 0.54 | 0.60 | 0.56 | 0.60 | 0.26 |
| | gaps < 50% mean height, lambda | | 0.81 | | 0.39 | | 0.65 | | 0.5 | | 0.42 | | 0.26 |

**Table S7: Average relative RMSE within sites for forest structure metrics (100 / 1000 m).** Shown is the average site-level RMSE of key forest structure metrics separated by algorithm and normalized by site-level standard deviation. Minimum size of gaps/clusters is 25 $m^2$. Combinations of metrics and algorithms with a lot of noise (rRMSEs >= 20%) are shown in red.

| **rME** across sites | **Metric** | **CHM$_{highest}$** 100 m | **CHM$_{highest}$** 1000 m | **CHM$_{tin}$** 100 m | **CHM$_{tin}$** 1000 m | **CHM$_{pitfree}$** 100 m | **CHM$_{pitfree}$** 1000 m | **CHM$_{spikefree}$** 100 m | **CHM$_{spikefree}$** 1000 m | **CHM$_{tspikefree}$** 100 m | **CHM$_{tspikefree}$** 1000 m | **CHM$_{lspikefree}$** 100 m | **CHM$_{lspikefree}$** 1000 m |
|---|---|---|---|---|---|---|---|---|---|---|---|---|---|
| **Vertical** | Mean height (m) | -0.17 | -0.18 | 0.02 | 0.02 | -0.12 | -0.12 | 0.03 | 0.03 | -0.08 | -0.08 | -0.02 | -0.02 |
| | Median of height (m) | -0.17 | -0.19 | 0.02 | 0.03 | -0.11 | -0.12 | 0.03 | 0.03 | -0.09 | -0.09 | -0.02 | -0.02 |
| | 25th perc. of height (m) | -0.29 | -0.32 | 0.04 | 0.04 | -0.16 | -0.17 | 0.08 | 0.09 | -0.10 | -0.10 | -0.01 | 0.00 |
| | 75th perc. of height (m) | -0.10 | -0.11 | 0.00 | 0.00 | -0.09 | -0.10 | 0.01 | 0.01 | -0.06 | -0.07 | -0.02 | -0.02 |
| | 95th perc. of height (m) | -0.05 | -0.06 | -0.01 | -0.01 | -0.05 | -0.05 | -0.01 | 0.00 | -0.03 | -0.04 | -0.02 | -0.02 |
| | 99th perc. of height (m) | -0.04 | -0.05 | -0.01 | -0.01 | -0.04 | -0.04 | -0.01 | -0.01 | -0.02 | -0.03 | -0.02 | -0.02 |
| | SD of height (m) | 0.12 | 0.08 | -0.04 | -0.04 | 0.06 | 0.03 | -0.06 | -0.05 | -0.01 | -0.03 | -0.04 | -0.04 |
| | CV of height | 0.53 | 0.58 | -0.10 | -0.12 | 0.37 | 0.42 | -0.09 | -0.12 | 0.08 | 0.09 | 0.00 | 0.00 |
| | Robust CV of height | 0.51 | 0.53 | -0.10 | -0.10 | 0.32 | 0.34 | -0.12 | -0.12 | 0.10 | 0.10 | -0.01 | -0.01 |
| | IQR of height (m) | 0.21 | 0.14 | -0.03 | -0.03 | 0.06 | 0.02 | -0.06 | -0.05 | 0.01 | -0.02 | -0.03 | -0.04 |
| | Canopy cover at 2 m (%) | -0.42 | -0.44 | 0.06 | 0.06 | -0.33 | -0.34 | 0.09 | 0.09 | -0.06 | -0.06 | 0.00 | 0.00 |
| | Canopy cover at 10 m (%) | -0.17 | -0.17 | 0.02 | 0.02 | -0.11 | -0.11 | 0.04 | 0.04 | -0.07 | -0.07 | -0.01 | -0.01 |
| **Horizontal** | Rumple index | 1.06 | 1.16 | -0.11 | -0.12 | 0.39 | 0.43 | -0.33 | -0.36 | 0.27 | 0.29 | -0.12 | -0.13 |
| | Normalized rumple index | 1.95 | 1.86 | -0.36 | -0.38 | 0.53 | 0.51 | -0.72 | -0.72 | 0.24 | 0.23 | -0.31 | -0.31 |
| | Moran's I | -1.98 | -2.38 | 0.34 | 0.42 | -0.77 | -0.79 | 0.83 | 0.97 | -0.43 | -0.50 | 0.34 | 0.39 |
| | clusters < 25$^{th}$ perc., number | 1.69 | 1.79 | -0.23 | -0.27 | 0.31 | 0.35 | -0.33 | -0.32 | -0.01 | -0.01 | -0.20 | -0.21 |
| | clusters < 25$^{th}$ perc., size ($m^2$) | -0.81 | -0.97 | 0.04 | 0.27 | 0.18 | -0.20 | 0.24 | 0.31 | -0.11 | -0.26 | 0.06 | 0.11 |
| | clusters > 75$^{th}$ perc., number | 1.15 | 1.37 | -0.27 | -0.32 | 0.73 | 0.97 | -0.52 | -0.55 | 0.32 | 0.40 | -0.24 | -0.27 |
| | clusters > 75$^{th}$ perc., size ($m^2$) | -0.57 | -0.68 | 0.19 | 0.31 | -0.22 | -0.25 | 0.55 | 0.75 | -0.25 | -0.35 | 0.19 | 0.21 |
| | gaps < 10 m, number | 1.11 | 1.27 | -0.01 | -0.01 | 0.43 | 0.50 | -0.08 | -0.09 | 0.28 | 0.32 | -0.02 | -0.02 |
| | gaps < 10 m, size ($m^2$) | -0.16 | -0.11 | 0.00 | -0.01 | -0.01 | 0.00 | 0.00 | 0.00 | -0.08 | -0.01 | 0.00 | 0.02 |
| | gaps < 10 m, lambda | | 0.02 | | -0.02 | | -0.14 | | 0.02 | | -0.04 | | -0.03 |
| | gaps < 50% mean height, number | 1.66 | 2.07 | 0.04 | 0.05 | 0.52 | 0.64 | -0.09 | -0.10 | 0.31 | 0.38 | -0.03 | -0.04 |
| | gaps < 50% mean height, size ($m^2$) | -0.15 | -0.20 | -0.04 | -0.06 | 0.02 | -0.03 | -0.01 | -0.01 | -0.03 | -0.05 | 0.00 | 0.01 |
| | gaps < 50% mean height, lambda | | 0.19 | | 0.03 | | -0.08 | | 0.06 | | -0.03 | | 0.03 |

**Table S8: Relative bias across sites for forest structure metrics (100 / 1000 m).** Shown is the relative bias of key forest structure metrics across all nine TERN sites, separated by algorithm and normalized by the standard deviation across the nine sites. Minimum size of gaps/clusters is 25 $m^2$. Combinations of metrics and algorithms with strong bias (rMEs >= 10% or <= -10%) are shown in red.

| **rME** **within sites** | **Metric** | **CHM$_{highest}$** | | **CHM$_{tin}$** | | **CHM$_{pitfree}$** | | **CHM$_{spikefree}$** | | **CHM$_{tspikefree}$** | | **CHM$_{lspikefree}$** | |
|---|---|---|---|---|---|---|---|---|---|---|---|---|---|
| | | **100 m** | **1000 m** | **100 m** | **1000 m** | **100 m** | **1000 m** | **100 m** | **1000 m** | **100 m** | **1000 m** | **100 m** | **1000 m** |
| **Vertical** | Mean height (m) | -1.03 | -1.82 | 0.11 | 0.19 | -0.88 | -1.56 | 0.16 | 0.30 | -0.35 | -0.64 | -0.07 | -0.12 |
| | Median of height (m) | -1.01 | -1.87 | 0.38 | 1.18 | -0.78 | -1.52 | 0.31 | 0.97 | -0.32 | -0.53 | -0.06 | -0.09 |
| | 25th perc. of height (m) | -1.07 | -2.12 | 0.51 | 2.74 | -0.73 | -1.43 | 0.51 | 23.70 | -0.29 | -0.48 | 0.08 | 0.20 |
| | 75th perc. of height (m) | -0.63 | -1.13 | 0.01 | 0.02 | -0.79 | -1.36 | 0.03 | 0.07 | -0.30 | -0.56 | -0.10 | -0.17 |
| | 95th perc. of height (m) | -0.35 | -0.74 | -0.09 | -0.16 | -0.45 | -0.80 | -0.04 | -0.07 | -0.18 | -0.40 | -0.09 | -0.17 |
| | 99th perc. of height (m) | -0.26 | -0.59 | -0.09 | -0.18 | -0.34 | -0.66 | -0.05 | -0.09 | -0.13 | -0.31 | -0.08 | -0.17 |
| | SD of height (m) | 0.21 | 0.33 | -0.19 | -0.34 | -0.23 | -0.40 | -0.19 | -0.33 | -0.16 | -0.34 | -0.14 | -0.28 |
| | CV of height | 1.51 | 2.93 | -0.31 | -0.68 | 1.12 | 2.33 | -0.34 | -0.75 | 0.26 | 0.52 | -0.02 | -0.08 |
| | Robust CV of height | 1.45 | 2.56 | -0.40 | -0.74 | 1.06 | 2.01 | -0.45 | -0.81 | 0.29 | 0.52 | -0.04 | -0.09 |
| | IQR of height (m) | 0.28 | 0.42 | -0.09 | -0.13 | -0.34 | -0.78 | -0.12 | -0.17 | -0.13 | -0.33 | -0.11 | -0.28 |
| | Canopy cover at 2 m (%) | -1.46 | -3.73 | 0.25 | 0.55 | -1.16 | -2.70 | 0.36 | 0.85 | -0.22 | -0.52 | 0.02 | 0.09 |
| | Canopy cover at 10 m (%) | -0.76 | -1.64 | 0.05 | 0.07 | -0.48 | -1.08 | 0.15 | 0.28 | -0.29 | -0.58 | -0.03 | -0.06 |
| **Horizontal** | Rumple index | 3.26 | 5.84 | -0.43 | -0.77 | 0.92 | 1.77 | -1.15 | -2.03 | 0.57 | 0.98 | -0.57 | -0.99 |
| | Normalized rumple index | 4.80 | 8.34 | -0.69 | -1.22 | 1.62 | 3.23 | -1.48 | -2.71 | 0.61 | 1.09 | -0.77 | -1.45 |
| | Moran's I | -3.22 | -6.13 | 0.60 | 1.20 | -1.53 | -2.90 | 1.48 | 3.04 | -0.75 | -1.53 | 0.66 | 1.25 |
| | clusters < 25$^{th}$ perc., number | 4.04 | 10.19 | -0.36 | -0.89 | 0.93 | 2.34 | -0.67 | -1.11 | 0.29 | 0.77 | -0.30 | -0.60 |
| | clusters < 25$^{th}$ perc., size ($m^2$) | -1.29 | -2.75 | 0.15 | 0.75 | 0.51 | 0.84 | 0.53 | 1.10 | -0.05 | -0.33 | 0.27 | 0.69 |
| | clusters > 75$^{th}$ perc., number | 1.72 | 5.38 | -0.40 | -0.81 | 0.89 | 1.87 | -0.82 | -2.10 | 0.53 | 1.46 | -0.40 | -1.06 |
| | clusters > 75$^{th}$ perc., size ($m^2$) | -0.91 | -2.48 | 0.35 | 0.85 | -0.38 | -0.76 | 1.04 | 2.70 | -0.38 | -0.91 | 0.38 | 1.06 |
| | gaps < 10 m, number | 2.88 | 15.30 | -0.02 | -0.01 | 0.72 | 1.69 | -0.14 | -0.32 | 0.47 | 1.04 | -0.05 | -0.11 |
| | gaps < 10 m, size ($m^2$) | -1.14 | -11.72 | -0.24 | -11.20 | 0.09 | 0.04 | 0.01 | -0.06 | -0.05 | -0.21 | 0.04 | 0.10 |
| | gaps < 10 m, lambda | | 0.30 | | -0.01 | | -0.34 | | 0.14 | | -0.04 | | -0.12 |
| | gaps < 50% mean height, number | 2.72 | 6.12 | 0.09 | 0.23 | 0.94 | 2.33 | -0.12 | -0.23 | 0.51 | 1.13 | -0.06 | -0.13 |
| | gaps < 50% mean height, size ($m^2$) | -0.23 | -0.91 | -0.11 | -0.37 | 0.05 | -0.19 | -0.05 | -0.05 | -0.06 | -0.33 | 0.01 | 0.06 |
| | gaps < 50% mean height, lambda | | 0.54 | | 0.10 | | -0.10 | | 0.18 | | -0.06 | | 0.05 |

**Table S9: Average relative bias within sites for forest structure metrics (100 / 1000 m).** Shown is the average site-level bias of key forest structure metrics separated by algorithm and normalized by site-level standard deviation. Minimum size of gaps/clusters is 25 $m^2$. Combinations of metrics and algorithms with strong bias (rMEs >= 10% or <= -10%) are shown in red.

| $R2_{rankswapped}$ aross sites | Metric | $CHM_{highest}$ | | $CHM_{tin}$ | | $CHM_{pitfree}$ | | $CHM_{spikefree}$ | | $CHM_{tspikefree}$ | | $CHM_{lspikefree}$ | |
|---|---|---|---|---|---|---|---|---|---|---|---|---|---|
| | | 100 m | 1000 m | 100 m | 1000 m | 100 m | 1000 m | 100 m | 1000 m | 100 m | 1000 m | 100 m | 1000 m |
| **Vertical** | Mean height (m) | 0.93 | 0.91 | 1 | 1 | 0.94 | 0.92 | 1 | 1 | 0.99 | 0.99 | 1 | 1 |
| | Median of height (m) | 0.86 | 0.85 | 0.98 | 0.97 | 0.91 | 0.9 | 0.98 | 0.98 | 0.99 | 0.99 | 1 | 1 |
| | 25th perc. of height (m) | 0.60 | 0.59 | 0.94 | 0.93 | 0.79 | 0.78 | 0.94 | 0.94 | 0.98 | 0.98 | 0.98 | 0.98 |
| | 75th perc. of height (m) | 0.95 | 0.94 | 0.99 | 0.99 | 0.94 | 0.93 | 1 | 1 | 0.99 | 0.98 | 1 | 1 |
| | 95th perc. of height (m) | 0.99 | 0.98 | 1 | 1 | 0.99 | 0.98 | 1 | 1 | 1 | 0.99 | 1 | 1 |
| | 99th perc. of height (m) | 0.99 | 0.99 | 1 | 1 | 0.99 | 0.99 | 1 | 1 | 1 | 1 | 1 | 1 |
| | SD of height (m) | 0.93 | 0.97 | 1 | 0.99 | 0.96 | 0.97 | 0.99 | 0.99 | 0.99 | 0.99 | 0.99 | 0.99 |
| | CV of height | 0.71 | 0.68 | 0.98 | 0.97 | 0.87 | 0.86 | 0.98 | 0.97 | 0.99 | 0.98 | 1 | 1 |
| | Robust CV of height | 0.71 | 0.68 | 0.98 | 0.97 | 0.87 | 0.86 | 0.98 | 0.97 | 0.99 | 0.98 | 1 | 1 |
| | IQR of height (m) | 0.68 | 0.78 | 0.99 | 0.98 | 0.81 | 0.84 | 0.98 | 0.98 | 0.95 | 0.96 | 0.99 | 0.98 |
| | Canopy cover at 2 m (%) | 0.80 | 0.82 | 0.99 | 0.99 | 0.84 | 0.82 | 0.98 | 0.98 | 0.99 | 0.99 | 1 | 0.99 |
| | Canopy cover at 10 m (%) | 0.93 | 0.94 | 1 | 1 | 0.97 | 0.96 | 0.99 | 0.99 | 0.99 | 0.99 | 1 | 1 |
| **Horizontal** | Rumple index | 0.51 | 0.48 | 0.98 | 0.96 | 0.82 | 0.81 | 0.83 | 0.81 | 0.93 | 0.93 | 0.95 | 0.93 |
| | Normalized rumple index | 0.08 | 0.12 | 0.78 | 0.80 | 0.82 | 0.84 | 0.45 | 0.52 | 0.88 | 0.89 | 0.87 | 0.87 |
| | Moran's I | 0.01 | 0.01 | 0.82 | 0.71 | 0.77 | 0.74 | 0.35 | 0.31 | 0.76 | 0.74 | 0.82 | 0.80 |
| | clusters < 25th perc., number | 0.04 | 0.03 | 0.79 | 0.85 | 0.49 | 0.59 | 0.79 | 0.83 | 0.79 | 0.83 | 0.87 | 0.92 |
| | clusters < 25th perc., size ($m^2$) | 0.01 | 0.01 | 0.69 | 0.84 | 0.34 | 0.40 | 0.61 | 0.75 | 0.64 | 0.79 | 0.72 | 0.88 |
| | clusters > 75th perc., number | 0.31 | 0.32 | 0.83 | 0.87 | 0.7 | 0.79 | 0.61 | 0.63 | 0.76 | 0.79 | 0.82 | 0.92 |
| | clusters > 75th perc., size ($m^2$) | 0.24 | 0.28 | 0.69 | 0.84 | 0.59 | 0.74 | 0.51 | 0.61 | 0.62 | 0.75 | 0.68 | 0.88 |
| | gaps < 10 m, number | 0.63 | 0.64 | 0.97 | 0.99 | 0.78 | 0.83 | 0.94 | 0.96 | 0.91 | 0.92 | 0.96 | 0.99 |
| | gaps < 10 m, size ($m^2$) | 0.88 | 0.91 | 0.95 | 1 | 0.89 | 0.97 | 0.95 | 0.99 | 0.93 | 0.98 | 0.95 | 0.99 |
| | gaps < 10 m, lambda | | 0.88 | | 0.94 | | 0.85 | | 0.80 | | 0.79 | | 0.96 |
| | gaps < 50% mean height, number | 0.19 | 0.14 | 0.93 | 0.97 | 0.59 | 0.63 | 0.90 | 0.94 | 0.8 | 0.86 | 0.91 | 0.99 |
| | gaps < 50% mean height, size ($m^2$) | 0.73 | 0.64 | 0.89 | 0.96 | 0.79 | 0.84 | 0.89 | 0.96 | 0.88 | 0.94 | 0.89 | 0.98 |
| | gaps < 50% mean height, lambda | | 0.85 | | 0.97 | | 0.90 | | 0.96 | | 0.97 | | 0.99 |

**Table S10: Rank-consistency across sites for forest structure metrics (100 / 1000 m).** Shown is $R2_{Spearman}$ (when swapping 50% of grid cells between 2 and 16 pulses $m^{-2}$) of key forest structure metrics across all nine TERN sites, separated by algorithm and normalized by standard deviation across the sites. Minimum size of gaps/clusters is 25 $m^2$. Combinations of metrics and algorithms that do not preserve the ranking of grid cells well (rank consistency $R^2 < 0.9$) are shown in red.

| $R2_{rankswapped}$ within sites | Metric | $CHM_{highest}$ | | $CHM_{tin}$ | | $CHM_{pitfree}$ | | $CHM_{spikefree}$ | | $CHM_{tspikefree}$ | | $CHM_{lspikefree}$ | |
|---|---|---|---|---|---|---|---|---|---|---|---|---|---|
| | | 100 m | 1000 m | 100 m | 1000 m | 100 m | 1000 m | 100 m | 1000 m | 100 m | 1000 m | 100 m | 1000 m |
| **Vertical** | Mean height (m) | 0.33 | 0.18 | 0.97 | 0.91 | 0.52 | 0.32 | 0.95 | 0.86 | 0.82 | 0.61 | 0.98 | 0.95 |
| | Median of height (m) | 0.34 | 0.29 | 0.76 | 0.64 | 0.47 | 0.37 | 0.80 | 0.69 | 0.77 | 0.58 | 0.96 | 0.93 |
| | 25th perc. of height (m) | 0.29 | 0.29 | 0.64 | 0.52 | 0.35 | 0.30 | 0.66 | 0.51 | 0.75 | 0.54 | 0.87 | 0.75 |
| | 75th perc. of height (m) | 0.60 | 0.39 | 0.96 | 0.92 | 0.65 | 0.55 | 0.99 | 0.97 | 0.85 | 0.67 | 0.97 | 0.94 |
| | 95th perc. of height (m) | 0.80 | 0.53 | 0.98 | 0.95 | 0.79 | 0.63 | 0.99 | 0.98 | 0.93 | 0.80 | 0.98 | 0.93 |
| | 99th perc. of height (m) | 0.86 | 0.59 | 0.97 | 0.92 | 0.83 | 0.69 | 0.99 | 0.97 | 0.96 | 0.85 | 0.98 | 0.93 |
| | SD of height (m) | 0.72 | 0.49 | 0.93 | 0.82 | 0.74 | 0.58 | 0.93 | 0.82 | 0.90 | 0.76 | 0.96 | 0.88 |
| | CV of height | 0.21 | 0.19 | 0.77 | 0.54 | 0.40 | 0.28 | 0.75 | 0.48 | 0.86 | 0.65 | 0.98 | 0.94 |
| | Robust CV of height | 0.21 | 0.19 | 0.77 | 0.53 | 0.40 | 0.28 | 0.75 | 0.48 | 0.86 | 0.66 | 0.98 | 0.94 |
| | IQR of height (m) | 0.40 | 0.25 | 0.90 | 0.84 | 0.56 | 0.47 | 0.90 | 0.83 | 0.83 | 0.66 | 0.94 | 0.81 |
| | Canopy cover at 2 m (%) | 0.10 | 0.23 | 0.80 | 0.61 | 0.22 | 0.18 | 0.65 | 0.42 | 0.82 | 0.65 | 0.91 | 0.80 |
| | Canopy cover at 10 m (%) | 0.35 | 0.25 | 0.90 | 0.88 | 0.54 | 0.30 | 0.84 | 0.75 | 0.75 | 0.62 | 0.93 | 0.91 |
| **Horizontal** | Rumple index | 0.10 | 0.21 | 0.75 | 0.54 | 0.40 | 0.33 | 0.22 | 0.12 | 0.63 | 0.43 | 0.63 | 0.43 |
| | Normalized rumple index | 0.18 | 0.27 | 0.52 | 0.33 | 0.27 | 0.19 | 0.10 | 0.12 | 0.56 | 0.38 | 0.45 | 0.25 |
| | Moran's I | 0.14 | 0.27 | 0.57 | 0.34 | 0.29 | 0.08 | 0.11 | 0.13 | 0.45 | 0.18 | 0.46 | 0.25 |
| | clusters < $25^{th}$ perc., number | 0.25 | 0.39 | 0.48 | 0.51 | 0.10 | 0.22 | 0.33 | 0.31 | 0.37 | 0.28 | 0.43 | 0.36 |
| | clusters < $25^{th}$ perc., size ($m^2$) | 0.17 | 0.27 | 0.45 | 0.50 | 0.16 | 0.20 | 0.28 | 0.25 | 0.30 | 0.27 | 0.41 | 0.34 |
| | clusters > $75^{th}$ perc., number | 0.11 | 0.14 | 0.62 | 0.53 | 0.39 | 0.25 | 0.26 | 0.10 | 0.45 | 0.31 | 0.55 | 0.51 |
| | clusters > $75^{th}$ perc., size ($m^2$) | 0.11 | 0.16 | 0.49 | 0.49 | 0.35 | 0.23 | 0.23 | 0.08 | 0.41 | 0.28 | 0.49 | 0.45 |
| | gaps < 10 m, number | 0.26 | 0.17 | 0.70 | 0.83 | 0.32 | 0.44 | 0.60 | 0.53 | 0.48 | 0.29 | 0.71 | 0.84 |
| | gaps < 10 m, size ($m^2$) | 0.50 | 0.25 | 0.73 | 0.87 | 0.47 | 0.48 | 0.74 | 0.83 | 0.68 | 0.72 | 0.75 | 0.84 |
| | gaps < 10 m, $\lambda$ | | 0.51 | | 0.78 | | 0.46 | | 0.60 | | 0.56 | | 0.79 |
| | gaps < 50% mean height, number | 0.17 | 0.23 | 0.71 | 0.79 | 0.20 | 0.28 | 0.66 | 0.68 | 0.48 | 0.47 | 0.70 | 0.91 |
| | gaps < 50% mean height, size ($m^2$) | 0.39 | 0.22 | 0.60 | 0.61 | 0.48 | 0.40 | 0.64 | 0.72 | 0.65 | 0.71 | 0.74 | 0.85 |
| | gaps < 50% mean height, $\lambda$ | | 0.52 | | 0.87 | | 0.64 | | 0.81 | | 0.84 | | 0.93 |

**Table S11: Average rank-consistency within sites for forest structure metrics (100 / 1000 m).** Shown is the average site-level $R2_{Spearman}$ (when swapping 50% of grid cells between 2 and 16 pulses $m^{-2}$) of key forest structure metrics separated by algorithm and normalized by site-level standard deviation. Minimum size of gaps/clusters is 25 $m^2$. Combinations of metrics and algorithms that do not preserve the ranking of grid cells well (rank consistency $R^2 < 0.9$) are shown in red.

| $R2_{rankswapped}$ within sites HIGH-QUALITY SCANS | Metric | $CHM_{highest}$ | | $CHM_{tin}$ | | $CHM_{pitfree}$ | | $CHM_{spikefree}$ | | $CHM_{tspikefree}$ | | $CHM_{lspikefree}$ | |
|---|---|---|---|---|---|---|---|---|---|---|---|---|---|
| | | 100 m | 1000 m | 100 m | 1000 m | 100 m | 1000 m | 100 m | 1000 m | 100 m | 1000 m | 100 m | 1000 m |
| **Vertical** | Mean height (m) | 0.92 | 0.79 | 1 | 0.98 | 0.90 | 0.78 | 1 | 0.98 | 1 | 0.99 | 1 | 1 |
| | Median of height (m) | 0.90 | 0.73 | 0.95 | 0.87 | 0.82 | 0.66 | 0.96 | 0.91 | 0.99 | 0.98 | 0.99 | 0.99 |
| | 25th perc. of height (m) | 0.71 | 0.47 | 0.90 | 0.88 | 0.67 | 0.54 | 0.92 | 0.86 | 0.96 | 0.86 | 0.97 | 0.96 |
| | 75th perc. of height (m) | 0.97 | 0.93 | 0.99 | 0.98 | 0.93 | 0.86 | 1 | 0.99 | 1 | 0.99 | 1 | 1 |
| | 95th perc. of height (m) | 0.99 | 0.95 | 1 | 0.99 | 0.97 | 0.93 | 1 | 0.99 | 1 | 1 | 1 | 0.99 |
| | 99th perc. of height (m) | 0.99 | 0.96 | 0.99 | 0.99 | 0.98 | 0.94 | 1 | 0.99 | 1 | 1 | 1 | 0.99 |
| | SD of height (m) | 0.98 | 0.94 | 0.99 | 0.98 | 0.98 | 0.96 | 1 | 0.98 | 1 | 1 | 1 | 0.99 |
| | CV of height | 0.86 | 0.62 | 0.97 | 0.90 | 0.84 | 0.66 | 0.98 | 0.91 | 1 | 0.99 | 1 | 0.99 |
| | Robust CV of height | 0.86 | 0.63 | 0.97 | 0.90 | 0.84 | 0.68 | 0.98 | 0.91 | 1 | 0.99 | 1 | 0.99 |
| | IQR of height (m) | 0.94 | 0.84 | 0.99 | 0.97 | 0.94 | 0.90 | 0.99 | 0.98 | 1 | 0.99 | 1 | 0.99 |
| | Canopy cover at 2 m (%) | 0.76 | 0.51 | 0.97 | 0.93 | 0.80 | 0.58 | 0.97 | 0.90 | 0.98 | 0.99 | 0.98 | 0.97 |
| | Canopy cover at 10 m (%) | 0.91 | 0.84 | 0.94 | 0.97 | 0.94 | 0.90 | 0.97 | 0.96 | 0.95 | 0.97 | 0.97 | 0.97 |
| **Horizontal** | Rumple index | 0.73 | 0.50 | 0.99 | 0.95 | 0.86 | 0.72 | 0.88 | 0.68 | 0.99 | 0.98 | 0.95 | 0.87 |
| | Normalized rumple index | 0.61 | 0.41 | 0.97 | 0.89 | 0.77 | 0.62 | 0.81 | 0.60 | 0.98 | 0.98 | 0.92 | 0.80 |
| | Moran's I | 0.80 | 0.50 | 0.97 | 0.93 | 0.84 | 0.73 | 0.79 | 0.57 | 0.97 | 0.97 | 0.93 | 0.80 |
| | clusters < $25^{th}$ perc., number | 0.53 | 0.26 | 0.79 | 0.83 | 0.62 | 0.55 | 0.75 | 0.83 | 0.72 | 0.88 | 0.77 | 0.89 |
| | clusters < $25^{th}$ perc., size ($m^2$) | 0.51 | 0.35 | 0.70 | 0.73 | 0.56 | 0.52 | 0.64 | 0.79 | 0.58 | 0.66 | 0.69 | 0.82 |
| | clusters > $75^{th}$ perc., number | 0.76 | 0.73 | 0.82 | 0.92 | 0.75 | 0.71 | 0.78 | 0.74 | 0.76 | 0.97 | 0.81 | 0.81 |
| | clusters > $75^{th}$ perc., size ($m^2$) | 0.73 | 0.66 | 0.69 | 0.88 | 0.71 | 0.68 | 0.70 | 0.70 | 0.67 | 0.84 | 0.74 | 0.75 |
| | gaps < 10 m, number | 0.63 | 0.58 | 0.77 | 0.90 | 0.76 | 0.72 | 0.79 | 0.79 | 0.75 | 0.91 | 0.82 | 0.86 |
| | gaps < 10 m, size ($m^2$) | 0.82 | 0.82 | 0.76 | 0.91 | 0.81 | 0.89 | 0.85 | 0.92 | 0.79 | 0.96 | 0.86 | 0.85 |
| | gaps < 10 m, $\lambda$ | | 0.87 | | 0.82 | | 0.87 | | 0.93 | | 0.90 | | 0.90 |
| | gaps < 50% mean height, number | 0.71 | 0.58 | 0.80 | 0.94 | 0.74 | 0.74 | 0.82 | 0.93 | 0.75 | 0.95 | 0.82 | 0.98 |
| | gaps < 50% mean height, size ($m^2$) | 0.81 | 0.83 | 0.73 | 0.82 | 0.76 | 0.74 | 0.78 | 0.86 | 0.75 | 0.91 | 0.84 | 0.90 |
| | gaps < 50% mean height, $\lambda$ | | 0.91 | | 0.91 | | 0.96 | | 0.94 | | 0.92 | | 0.96 |

**Table S12: Average rank-consistency within sites for high-quality scans (100 / 1000 m).** Same as preceding table, but swapping 50% of grid cells between 8 and 16 pulses $m^{-2}$ instead of 2 and 16. Minimum size of gaps/clusters is 25 $m^2$. Combinations of metrics and algorithms that do not preserve the ranking of grid cells well (rank consistency $R^2 < 0.9$) are shown in red.

| $R2_{rankswapped}$ within sites WITHIN-SCAN PD | Metric | $CHM_{highest}$ | | $CHM_{tin}$ | | $CHM_{pitfree}$ | | $CHM_{spikefree}$ | | $CHM_{tspikefree}$ | | $CHM_{lspikefree}$ | |
|---|---|---|---|---|---|---|---|---|---|---|---|---|---|
| | | 100 m | 1000 m | 100 m | 1000 m | 100 m | 1000 m | 100 m | 1000 m | 100 m | 1000 m | 100 m | 1000 m |
| **Vertical** | Mean height (m) | 0.98 | 0.96 | 1 | 1 | 0.98 | 0.97 | 0.98 | 0.96 | 0.99 | 0.98 | 1 | 0.99 |
| | Median of height (m) | 0.97 | 0.95 | 0.99 | 0.99 | 0.97 | 0.95 | 0.96 | 0.95 | 0.99 | 0.98 | 0.99 | 0.99 |
| | 25th perc. of height (m) | 0.91 | 0.88 | 0.95 | 0.98 | 0.93 | 0.94 | 0.95 | 0.94 | 0.97 | 0.98 | 0.98 | 0.99 |
| | 75th perc. of height (m) | 0.99 | 0.98 | 1 | 1 | 0.97 | 0.98 | 0.98 | 0.97 | 0.99 | 0.99 | 1 | 1 |
| | 95th perc. of height (m) | 1 | 0.99 | 1 | 1 | 0.98 | 0.99 | 0.99 | 0.98 | 1 | 0.99 | 1 | 0.99 |
| | 99th perc. of height (m) | 1 | 0.99 | 0.99 | 1 | 0.99 | 0.99 | 0.99 | 0.99 | 0.99 | 0.99 | 1 | 1 |
| | SD of height (m) | 0.99 | 0.99 | 1 | 0.99 | 0.98 | 0.98 | 0.99 | 0.98 | 1 | 0.99 | 1 | 1 |
| | CV of height | 0.96 | 0.94 | 0.99 | 0.99 | 0.97 | 0.94 | 0.97 | 0.95 | 0.99 | 0.99 | 0.99 | 0.99 |
| | Robust CV of height | 0.96 | 0.95 | 0.99 | 0.99 | 0.97 | 0.94 | 0.97 | 0.95 | 0.99 | 0.99 | 0.99 | 0.99 |
| | IQR of height (m) | 0.96 | 0.95 | 0.99 | 0.99 | 0.97 | 0.98 | 0.98 | 0.97 | 0.99 | 0.98 | 0.99 | 0.99 |
| | Canopy cover at 2 m (%) | 0.92 | 0.91 | 0.98 | 0.99 | 0.94 | 0.92 | 0.96 | 0.93 | 0.97 | 0.99 | 0.96 | 0.98 |
| | Canopy cover at 10 m (%) | 0.93 | 0.95 | 0.92 | 0.97 | 0.94 | 0.95 | 0.93 | 0.95 | 0.94 | 0.97 | 0.95 | 0.97 |
| **Horizontal** | Rumple index | 0.86 | 0.77 | 0.99 | 0.99 | 0.96 | 0.91 | 0.91 | 0.85 | 0.97 | 0.94 | 0.98 | 0.98 |
| | Normalized rumple index | 0.80 | 0.72 | 0.97 | 0.98 | 0.92 | 0.90 | 0.89 | 0.82 | 0.95 | 0.95 | 0.97 | 0.98 |
| | Moran's I | 0.84 | 0.70 | 0.96 | 0.95 | 0.91 | 0.92 | 0.86 | 0.76 | 0.94 | 0.89 | 0.97 | 0.98 |
| | clusters < $25^{th}$ perc., number | 0.60 | 0.57 | 0.78 | 0.88 | 0.71 | 0.81 | 0.72 | 0.86 | 0.71 | 0.91 | 0.73 | 0.93 |
| | clusters < $25^{th}$ perc., size ($m^2$) | 0.49 | 0.55 | 0.67 | 0.77 | 0.63 | 0.68 | 0.60 | 0.78 | 0.58 | 0.80 | 0.63 | 0.81 |
| | clusters > $75^{th}$ perc., number | 0.77 | 0.85 | 0.79 | 0.98 | 0.78 | 0.89 | 0.76 | 0.82 | 0.74 | 0.91 | 0.77 | 0.96 |
| | clusters > $75^{th}$ perc., size ($m^2$) | 0.69 | 0.76 | 0.65 | 0.93 | 0.71 | 0.72 | 0.67 | 0.72 | 0.64 | 0.69 | 0.69 | 0.82 |
| | gaps < 10 m, number | 0.58 | 0.74 | 0.73 | 0.87 | 0.72 | 0.83 | 0.74 | 0.83 | 0.73 | 0.86 | 0.75 | 0.88 |
| | gaps < 10 m, size ($m^2$) | 0.81 | 0.87 | 0.76 | 0.90 | 0.82 | 0.88 | 0.82 | 0.91 | 0.80 | 0.93 | 0.82 | 0.91 |
| | gaps < 10 m, $\lambda$ | | 0.91 | | 0.81 | | 0.90 | | 0.88 | | 0.88 | | 0.89 |
| | gaps < 50% mean height, number | 0.73 | 0.81 | 0.76 | 0.88 | 0.72 | 0.90 | 0.77 | 0.91 | 0.73 | 0.89 | 0.76 | 0.94 |
| | gaps < 50% mean height, size ($m^2$) | 0.77 | 0.85 | 0.70 | 0.89 | 0.77 | 0.87 | 0.75 | 0.83 | 0.74 | 0.86 | 0.78 | 0.91 |
| | gaps < 50% mean height, $\lambda$ | | 0.92 | | 0.94 | | 0.96 | | 0.92 | | 0.94 | | 0.95 |

**Table S13: Average rank-consistency under within-scan heterogeneity (100 / 1000 m).** Same as preceding two tables, but swapping 50% of grid cells between two scans of 4 pulses $m^{-2}$, one with within-scan variation in pulse density, the other one without. Minimum size of gaps/clusters is 25 $m^2$. Combinations of metrics and algorithms that do not preserve the ranking of grid cells well (rank consistency $R^2 < 0.9$) are shown in red.

| Algorithm | A | b | AGB (25 ha) | RMSE (25 ha) | AGB (2 $m^{-2}$) | AGB (16 $m^{-2}$) | RMSE (pd) | $R2_{rankswapped}$ |
|---|---|---|---|---|---|---|---|---|
| highest | 2.321 | 1.606 | 427.97 | 41.62 | 438.46 | 484.67 | 49.15 | 0.67 |
| tin | 4.69 | 1.452 | 427.95 | 40.47 | 482.27 | 487.92 | 16.56 | 0.96 |
| pitfree | 3.45 | 1.493 | 427.96 | 42.59 | 449.2 | 479.52 | 34.24 | 0.82 |
| spikefree | 3.504 | 1.519 | 427.96 | 41 | 486.92 | 486.97 | 16.13 | 0.97 |
| tspikefree | 3.458 | 1.512 | 427.97 | 41.48 | 462.98 | 484.34 | 26.30 | 0.90 |
| lspikefree | 3.118 | 1.535 | 427.97 | 41.94 | 477.19 | 483.3 | 16.11 | 0.96 |

**Table S14: Summary statistics for biomass estimation at Robson Creek at 1 ha scale.** Shown are summary statistics for above ground biomass (AGB, in Mg $ha^{-1}$) at Robson Creek for 100 m x 100 m grid cells. Grey statistics provide model parameters for the model described in section S2 and fitted to 25 1 ha plots, as well as mean predicted AGB across the 25 1 ha plots and its root mean squared error (RMSE, also in Mg $ha^{-1}$). AGB estimates in black colour indicate whole-site estimates of biomass at either 2 or 16 pulses $m^{-2}$, as well as the RMSE, and $R2_{rankswapped}$, which shows the consistency of AGB under random swapping of grid cells between 2 and 16 pulses $m^{-2}$.

### C. Figures

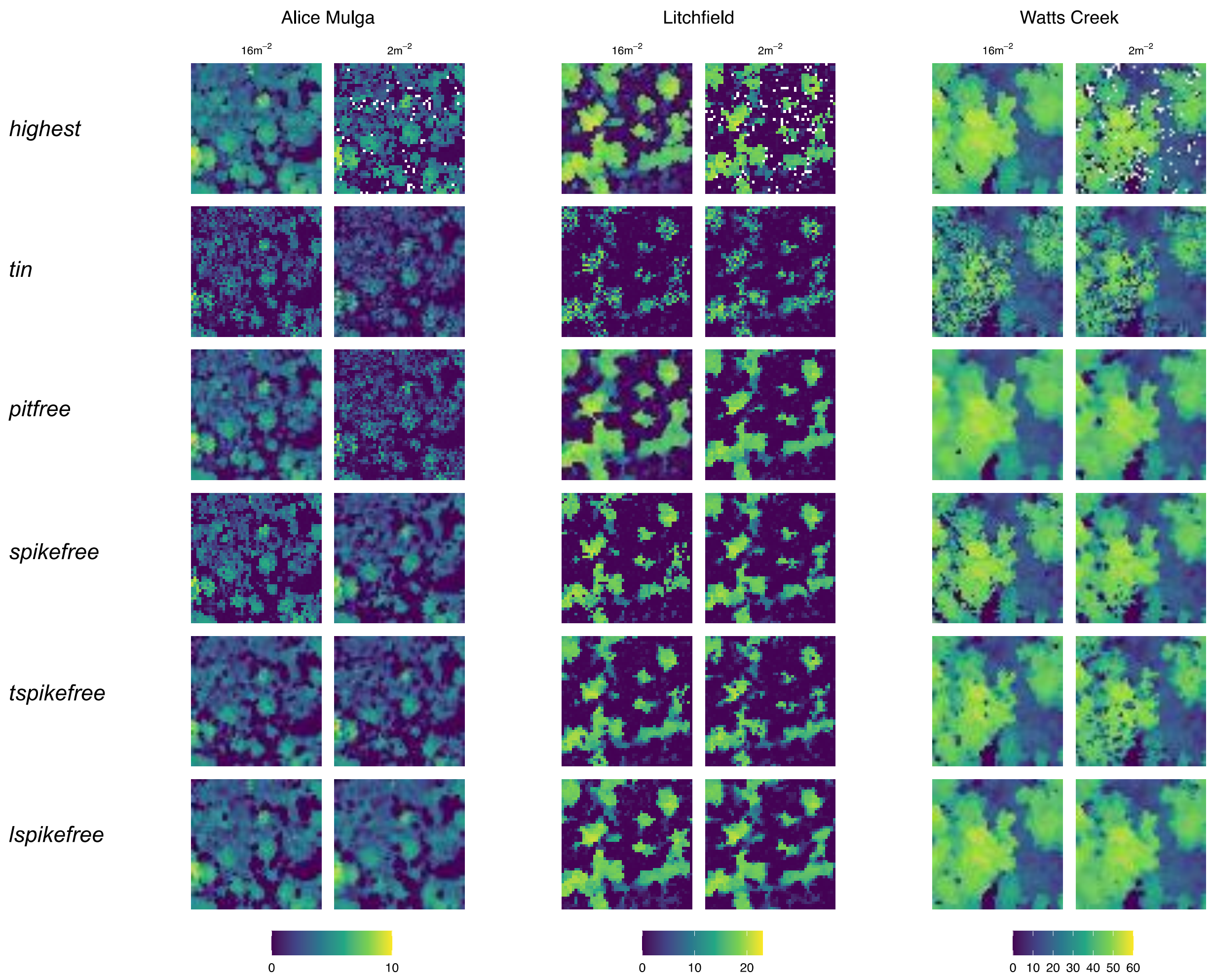

**Figure S1: CHM degradation between high and low pulse density at three sample sites**. Shown are changes in canopy height (colour bars, in m) between high (16 $m^{-2}$) and low (2 $m^{-2}$) pulse densities over an extent of 50 m x 50 m at three sample sites. Each row corresponds to a different CHM algorithm.

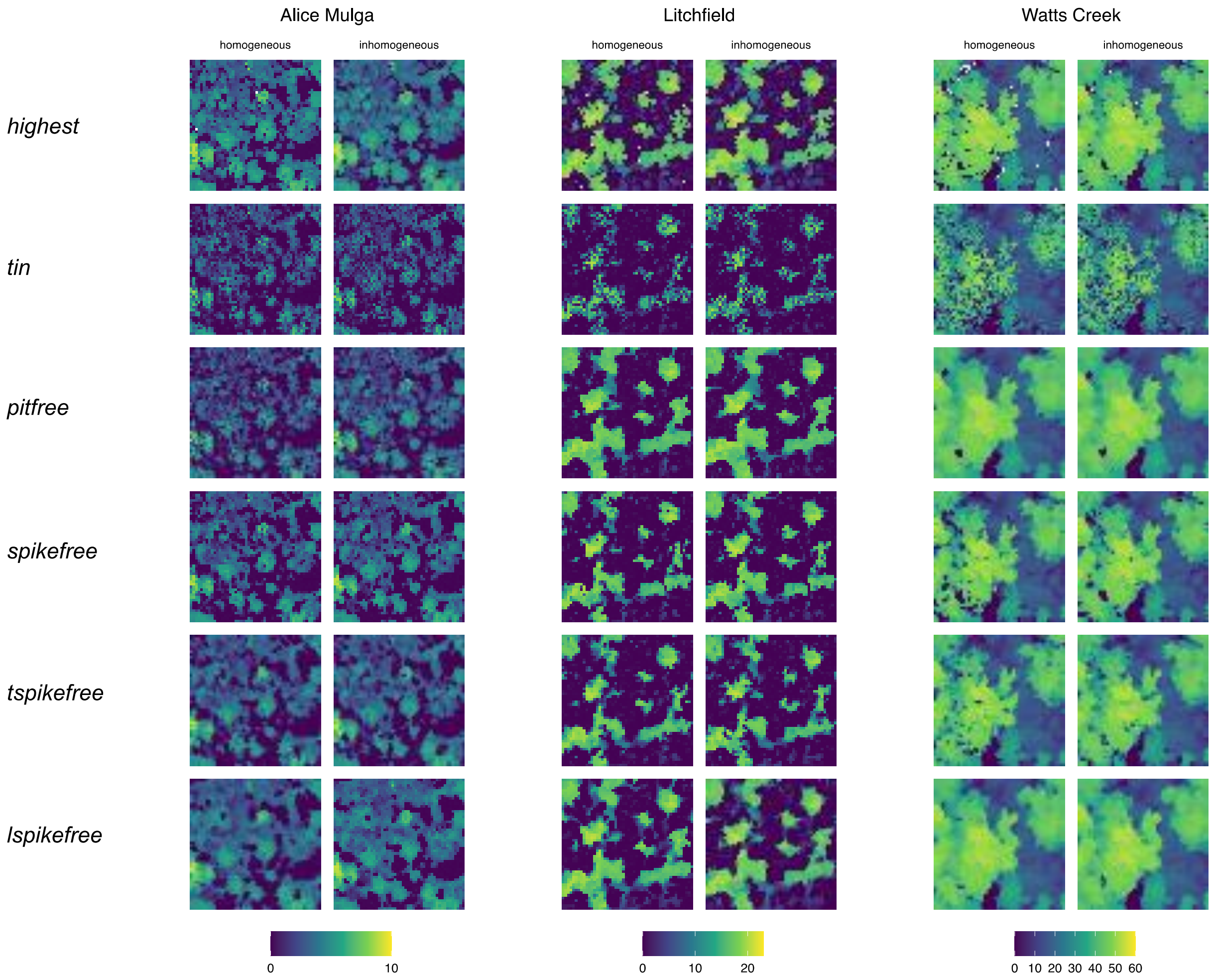

**Figure S2: CHM degradation between homogeneous and inhomogeneous pulse density**. Shown are changes in canopy height (colour bars, in m) between point clouds that nominally have the same average pulse density (4 $m^{-2}$), but where one has been homogenized, and the other not (retaining original pulse density patterns), over an extent of 50 m x 50 m at three sample sites. Each row corresponds to a different CHM algorithm. Note that in this example, we picked plots in overlap areas of flightlines to also showcase scan angle effects (Figure S4). This means, that pulse densities are higher in the inhomogeneous point cloud due to better sampling.

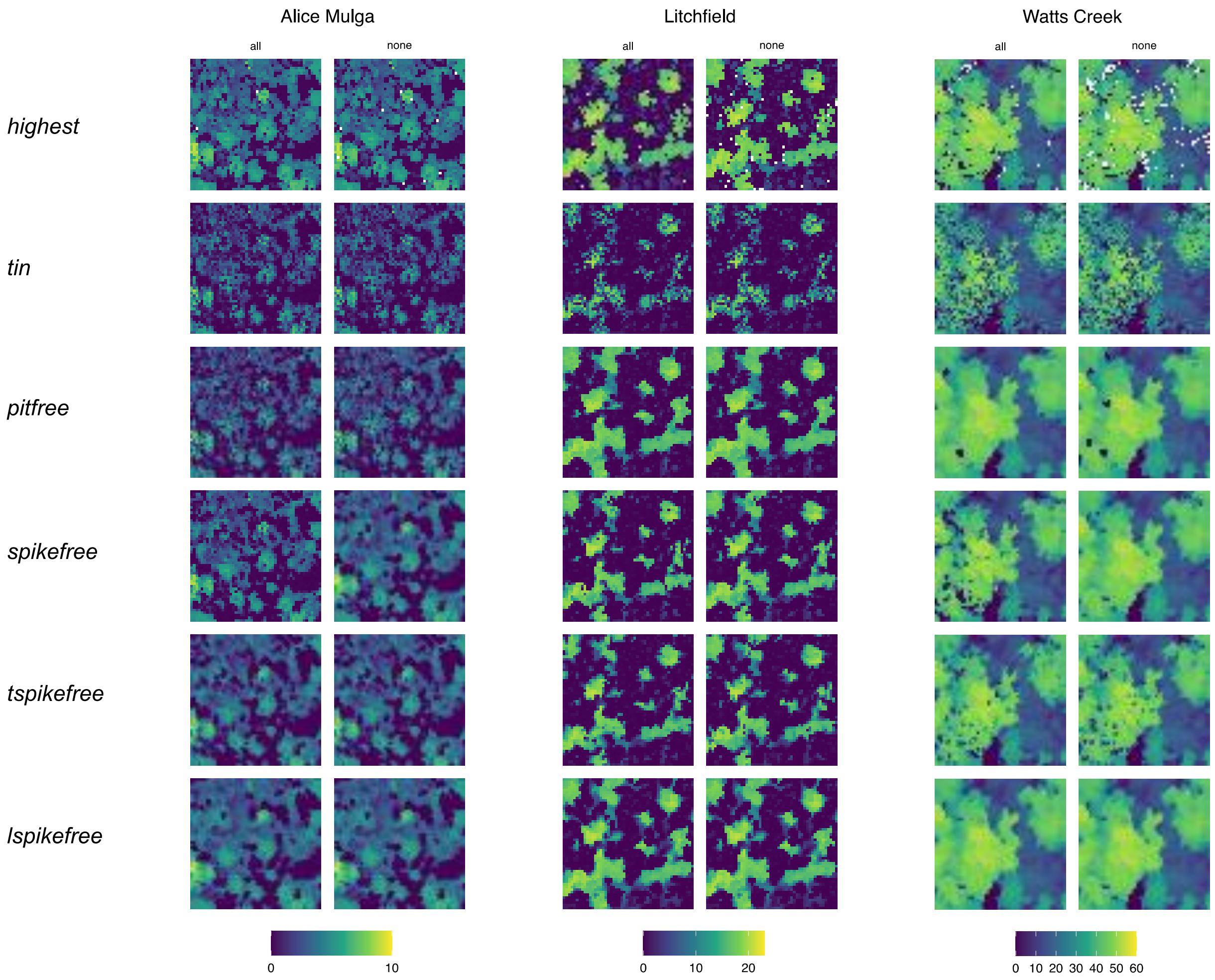

**Figure S3: CHM degradation between high and no laser penetration into the canopy**. Shown are changes in canopy height (colour bars, in m) between point clouds with full laser penetration ("all" returns) and those with no higher-order returns ("none") over an extent of 50 m x 50 m at three sample sites. Each row corresponds to a different CHM algorithm.

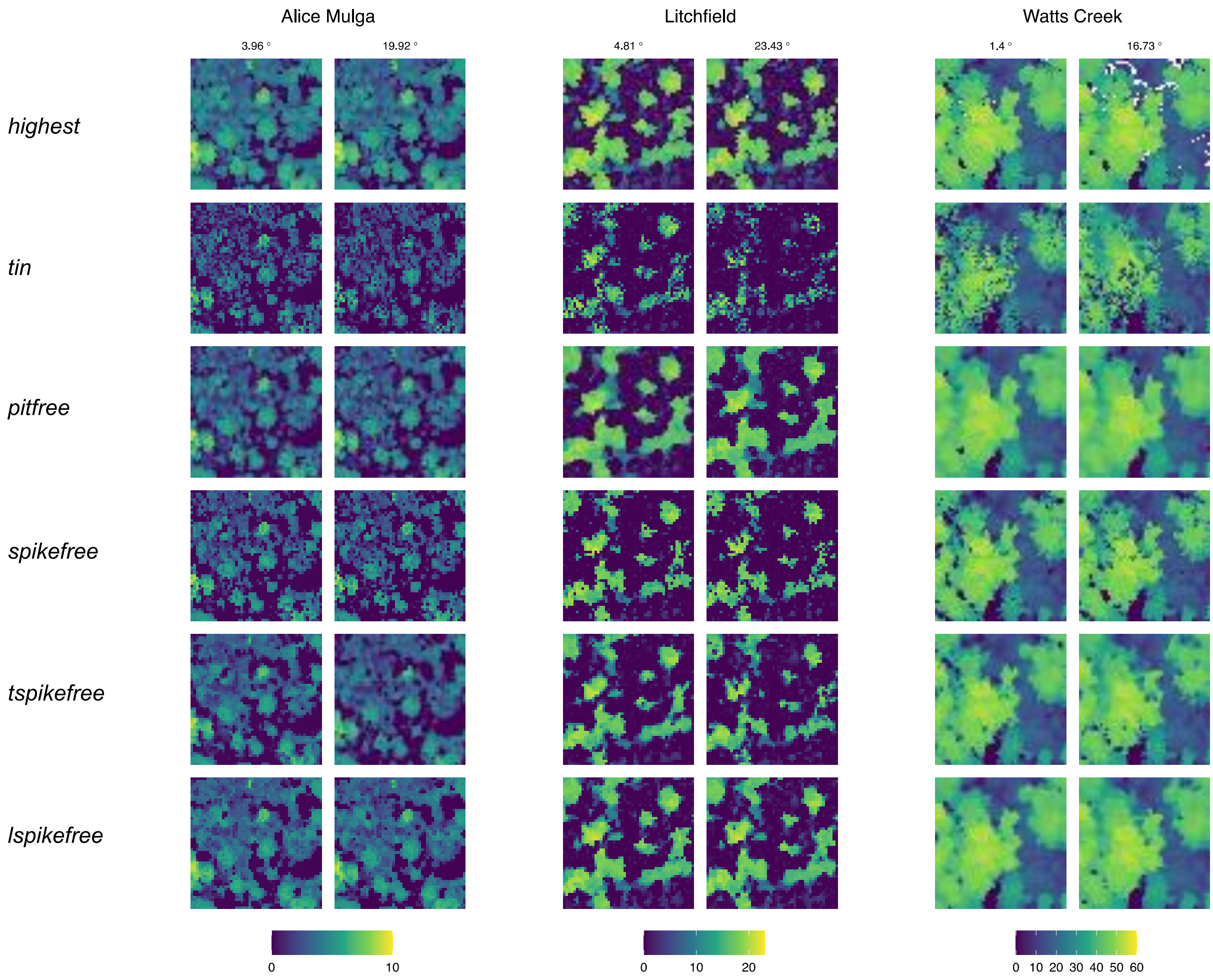

**Figure S4: CHM degradation between low and high scan angles**. Shown are changes in canopy height (colour bars, in m) between low and high scan angles from two overlapping flight lines over an extent of 50 m x 50 m at three sample sites. Each row corresponds to a different CHM algorithm. Note how $CHM_{tin}$, generally one of the most robust algorithms, is susceptible to scan angle differences (degradation of large tree crowns at Litchfield / an enlarged tree crown at Watts Creek).

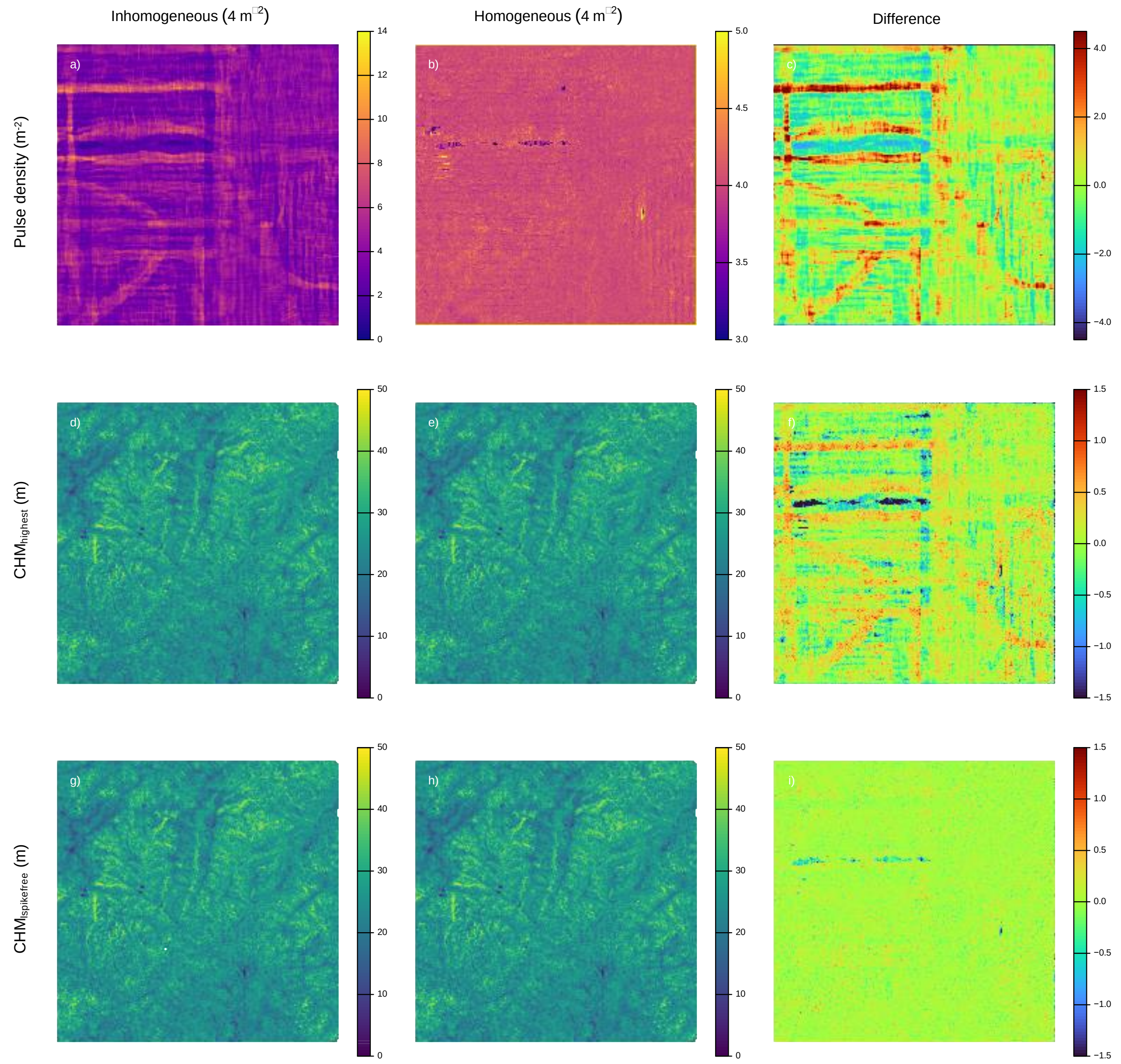

**Figure S5: Effects of inhomogeneous pulse densities within scans on canopy height estimates.** Shown are the effects of inhomogeneous downsampling of pulse density (thinning every $n^{th}$ pulse to a target density of 4 $m^{-2}$, column), homogeneous downsampling (thinning randomly within 25 m x 25 m cells to a target density of 4 $m^{-2}$, second column) and the difference between the two (third column) on canopy height models at Robson Creek, a tropical rain forest site. The first row shows the pulse density rasters as well as their differences, the second row shows the respective canopy height models and their differences based on the $CHM_{highest}$ algorithm (highest return per 1 $m^2$) and the third row shows the same for $CHM_{lspikefree}$, i.e., an algorithm that adapts to local pulse density variation. All rasters have been aggregated at 25 m resolution to remove small-scale noise, and the CHM difference rasters have been clamped between -1.5 and +1.5 to better visualize deviation patterns. As can be seen from the difference between panels f) and i), $CHM_{lspikefree}$ successfully compensates for the inhomogeneous pulse density patterns in g), whereas $CHM_{highest}$ under- and overestimates canopy height in line with local pulse density variation (0.5-1.5 m bias along flight lines). Note that Robson Creek was chosen due to its many, irregularly overlapping flightlines, but is likely to be the least sensitive to pulse density variation and thus should provide a lower bound on biases.

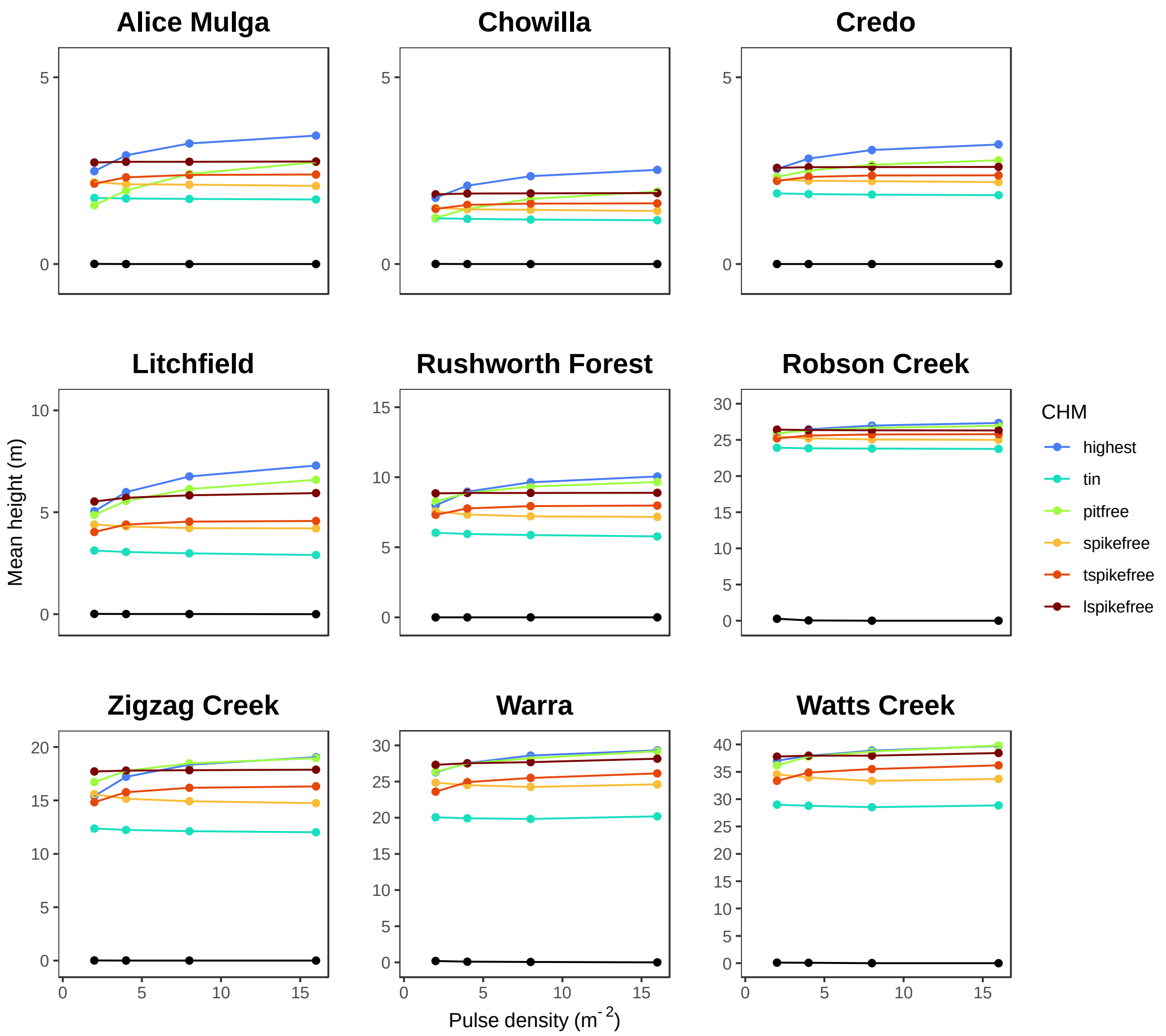

**Figure S6: Effects of pulse density on mean canopy height estimates across all nine SuperSites.** Same as Figure 2 in main text, included here for completeness. Shown are changes in mean canopy height with pulse density, separated into effects on topography (DTM) and canopy surface models (DSMs/CHMs, 6 algorithms). All rasters have been produced at 1 $m^2$ resolution and have been normalized by a common reference DTM. Axes are rescaled to the mean height of each site, and effects below 2 $m^{-2}$ are shown in lighter colours to indicate that we would not expect algorithms to perform well beyond this threshold due to sampling limitation. Note how there is little effect on DTMs in open systems such as woodlands and savannas (Alice Mulga, Chowilla, Credo, Litchfield), and even some closed-canopy forests (Zigzag Creek), but a noticeable effect in the densest and topographically most complex forest at Robson Creek. In contrast, pulse density variation has the smallest impacts on canopy height at Robson Creek, and some of the open systems experience the strongest changes in relative terms.

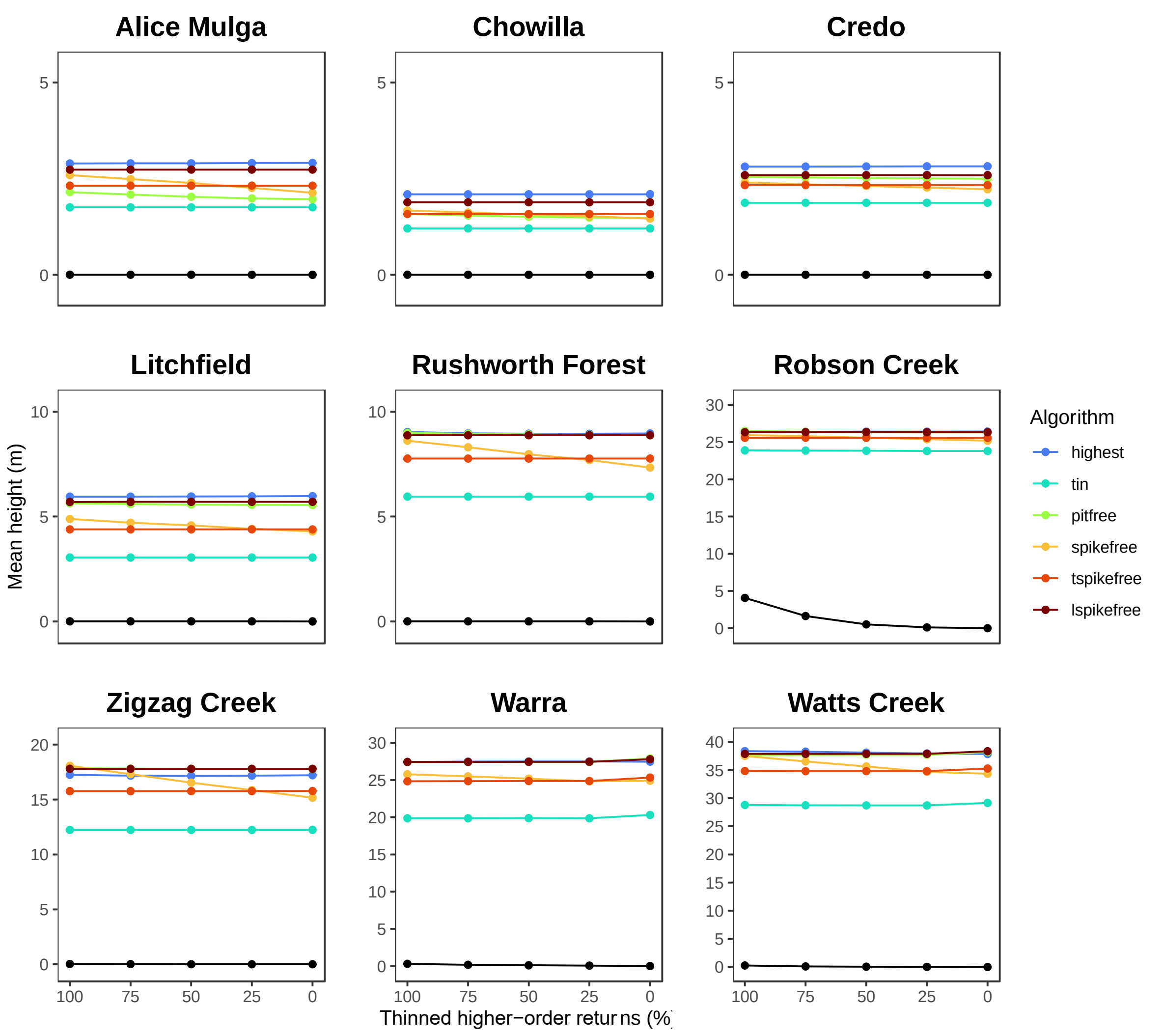

**Figure S7: Effects of laser penetration on mean canopy height estimates across all nine SuperSites.** Shown are changes in mean canopy height with an increasing percentage of thinned higher-order returns (2nd, 3rd, etc.), separated into effects on topography (DTM) and canopy surface models (DSMs/CHMs, 6 algorithms). All rasters have been produced at 1 $m^2$ resolution and have been normalized by a common reference DTM. Axes are rescaled to the mean height of each site. There is little effect on CHMs across algorithms (with the exception of $CHM_{spikefree}$), which is in accordance with the idea that surface models should not depend strongly on sub-canopy returns (or not at all, when only first returns are used as in $CHM_{tin}$ and $CHM_{lspikefree}$). Note, however the strong effect on the DTM in the densest and topographically most complex forest at Robson Creek, with biases of up to 5 m (which would induce corresponding biases in the CHM of -5 m).

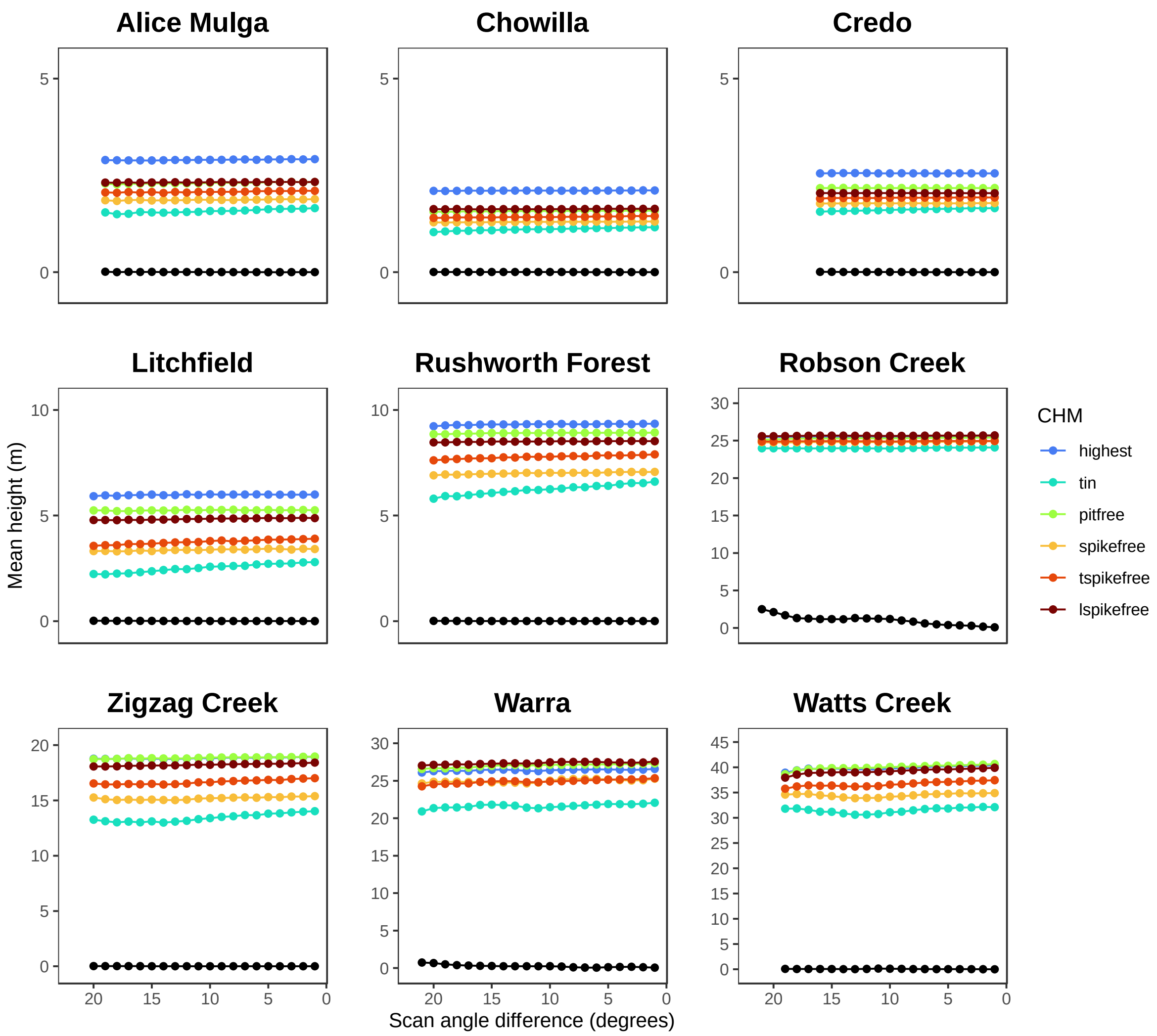

**Figure S8: Effects of scan angle on mean canopy height estimates across all nine SuperSites.** Shown are changes in mean canopy height when scan angles are increasing compared to a reference flightline. Effects are separated into effects on topography (DTM) and canopy surface models (DSMs/CHMs, 6 algorithms). All rasters have been produced at 1 $m^2$ resolution and have been normalized by a common reference DTM. Axes are rescaled to the mean height of each site. Effects on canopy surfaces are generally small, with a few more notable declines (e.g., $CHM_{tin}$ Rushworth Forest and Litchfield), but, as in Figures S6 and S7, there is a high stability in the CHM, but clear effect on the DTM in the densest and topographically most complex forest at Robson Creek.

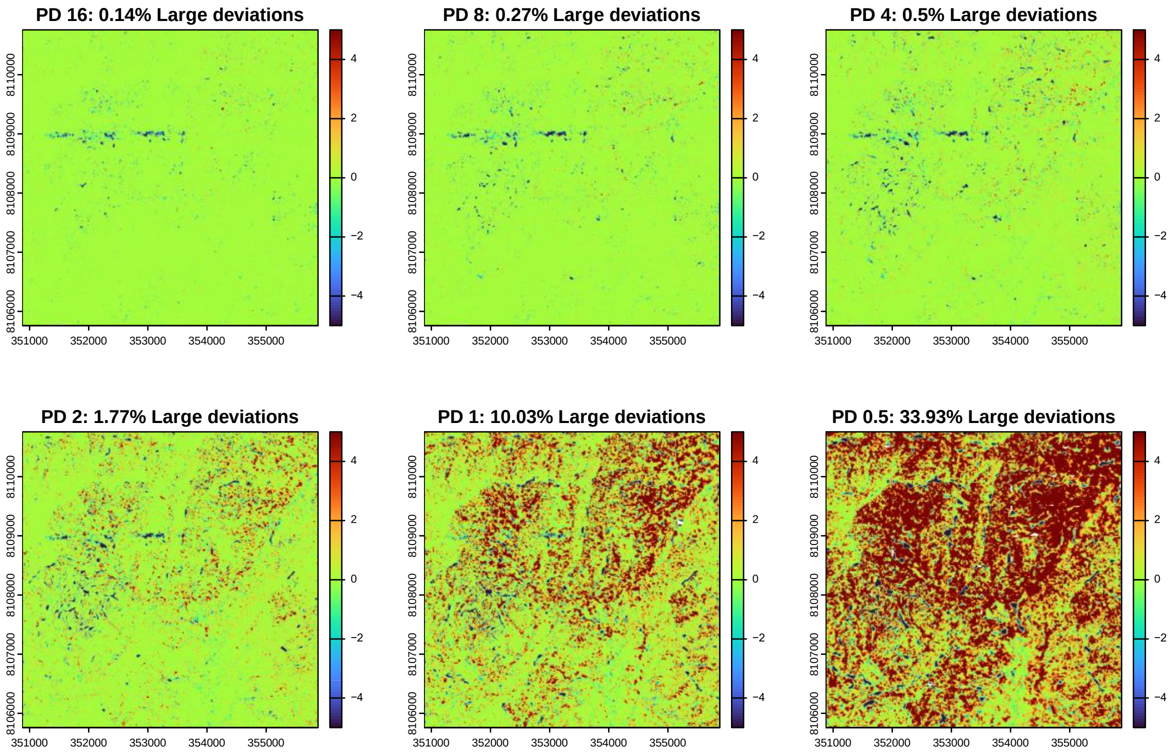

**Figure S9: Degradation of topography with pulse density at Robson Creek**. Shown are changes in topography from high (16 $m^{-2}$) to low (0.5 $m^{-2}$) pulse densities over the entire Robson Creek site. Colour scales indicate upward (red) and downward (blue) bias from a reference topographic model obtained at the scan's original pulse densities (~30 $m^{-2}$). Values are clamped between -5 m and 5 m to keep small scale changes visible. The narrow strip with strong downward deviations across pulse densities around y = 8109000 m has already very low pulse densities (~ 2 $m^{-2}$) in the original scan, so topography there is inherently difficult to reconstruct (cf. Figure S5).

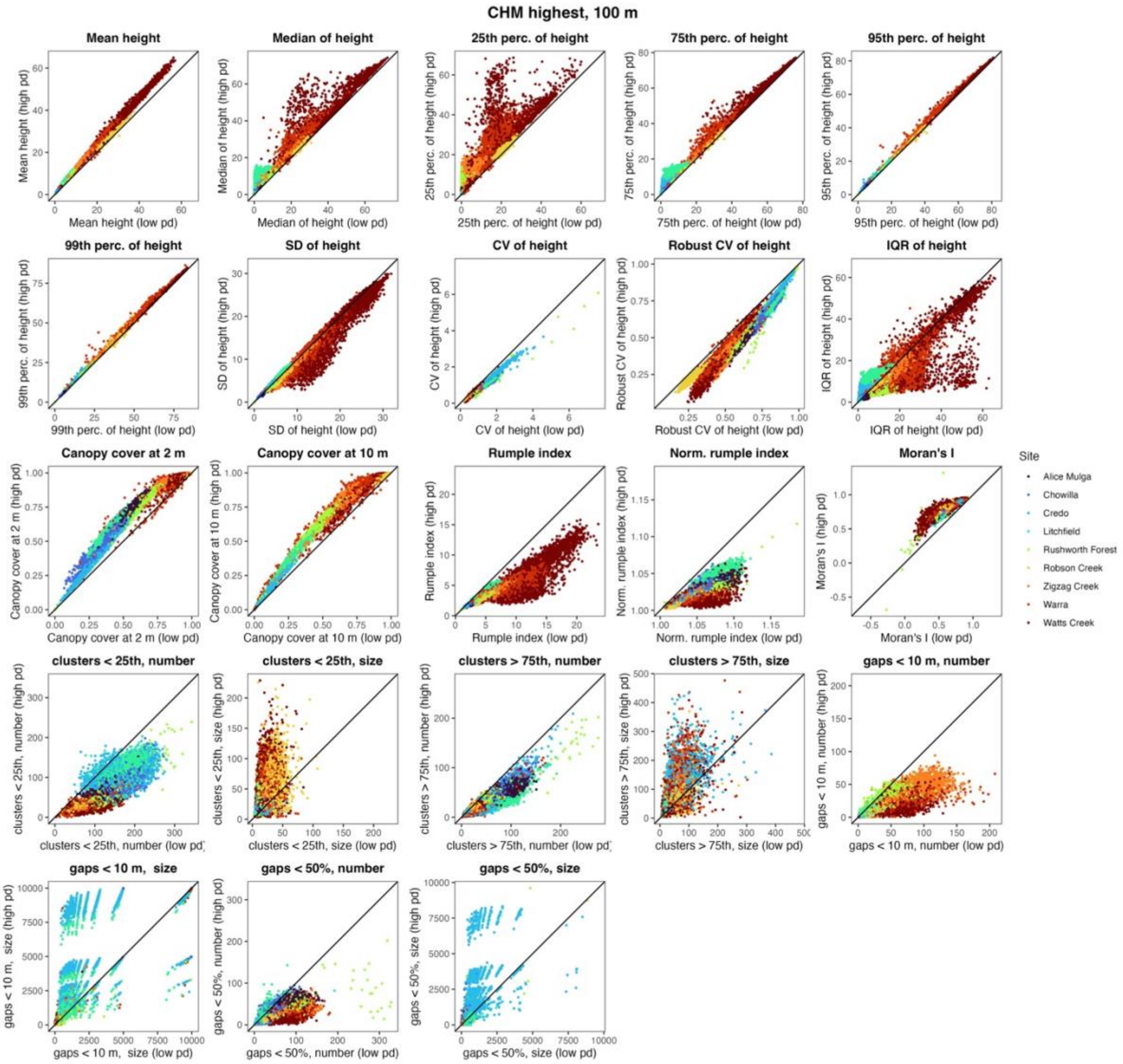

**Figure S10: Derived metrics for $CHM_{highest}$ at 100 m scale.** Shown are derived metrics for $CHM_{highest}$ at 100 m and across all sites together, compared between 2 pulses $m^{-2}$ (low pd) and 16 pulses $m^{-2}$ (high pd). Data points are coloured by site.

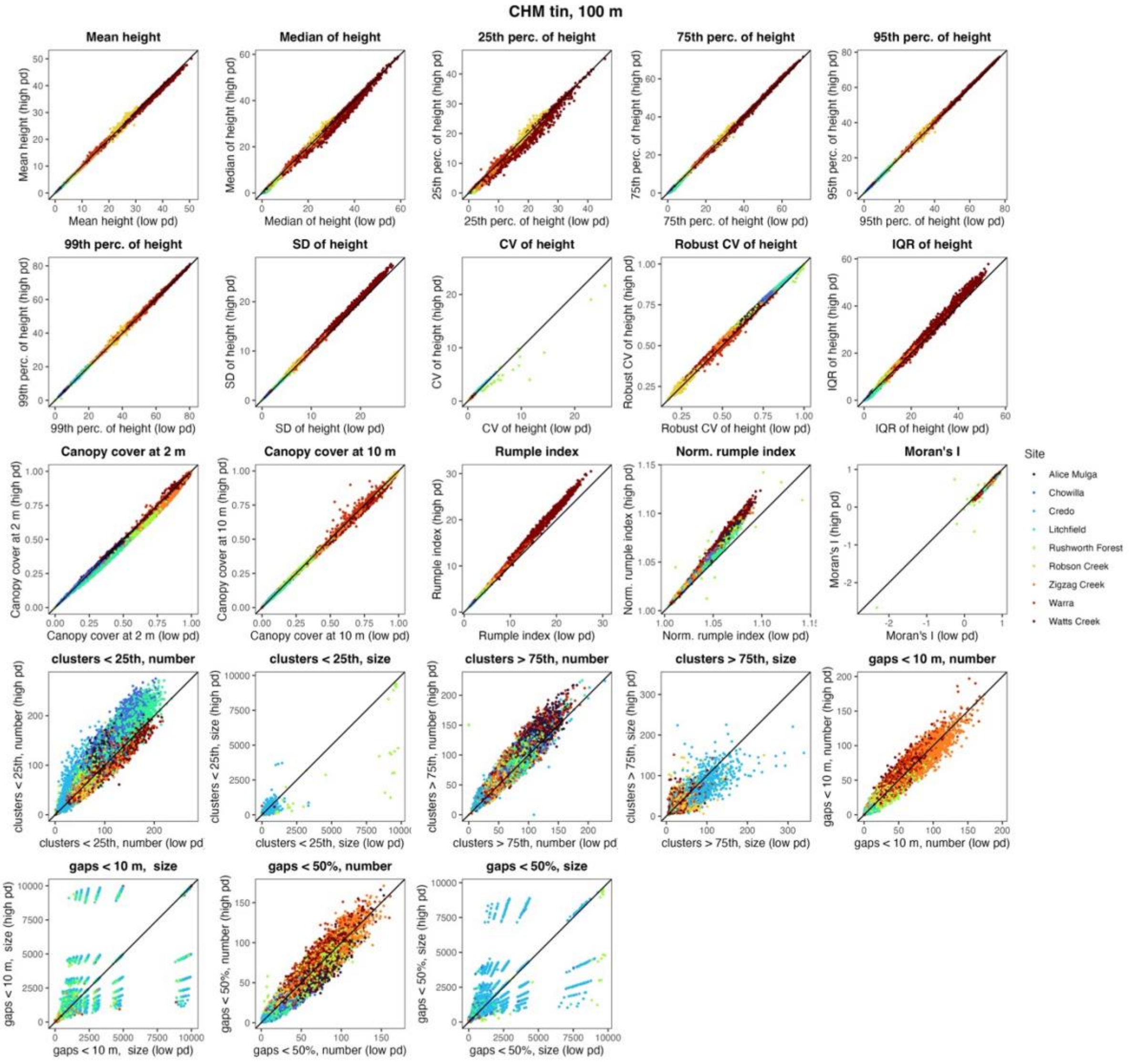

**Figure S11: Derived metrics for $CHM_{tin}$ at 100 m scale.** Shown are derived metrics for $CHM_{tin}$ at 100 m and across all sites together, compared between 2 pulses $m^{-2}$ (low pd) and 16 pulses $m^{-2}$ (high pd). Data points are coloured by site.

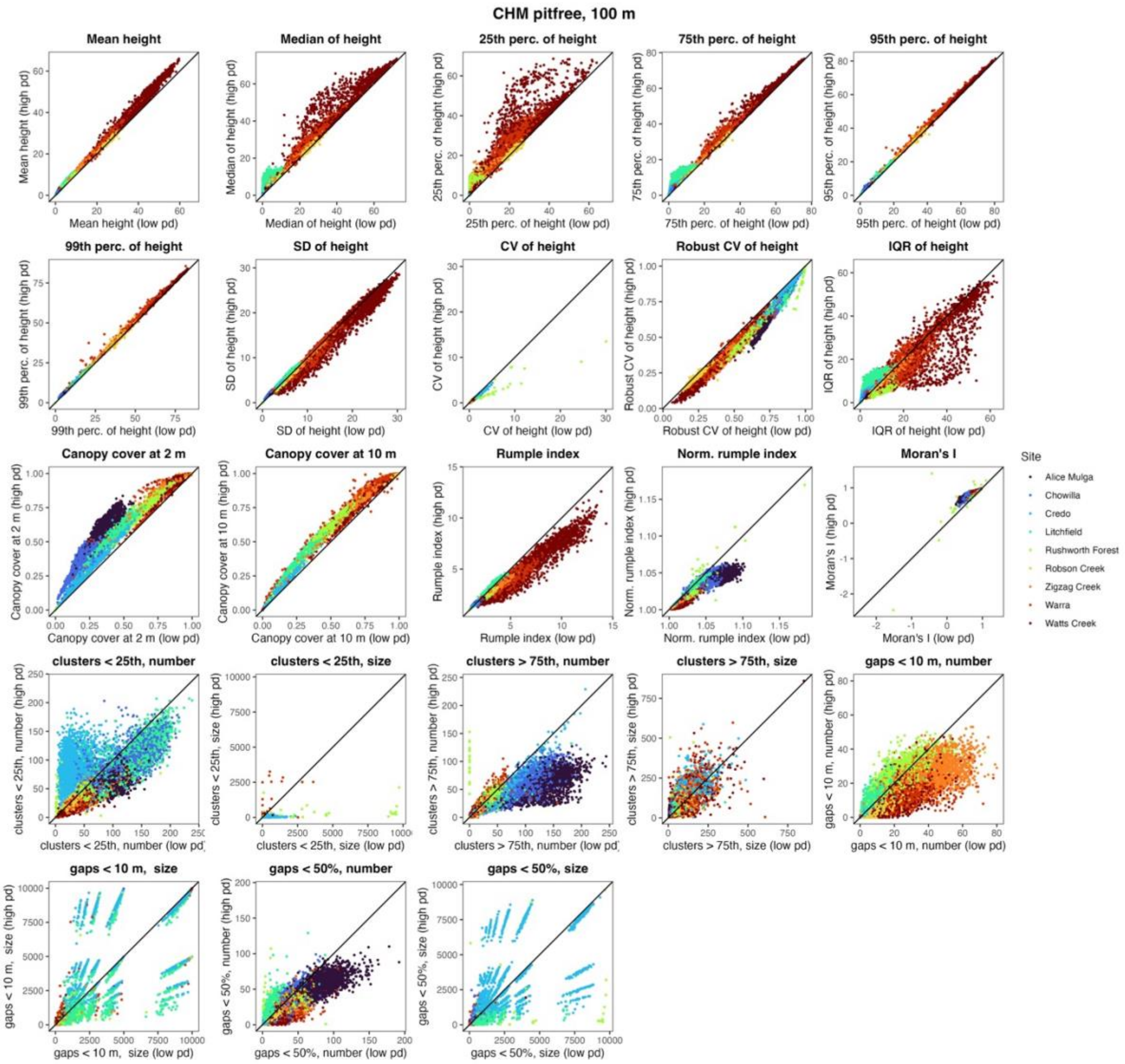

**Figure S12: Derived metrics for $CHM_{pitfree}$ at 100 m scale.** Shown are derived metrics for $CHM_{pitfree}$ at 100 m and across all sites together, compared between 2 pulses $m^{-2}$ (low pd) and 16 pulses $m^{-2}$ (high pd). Data points are coloured by site.

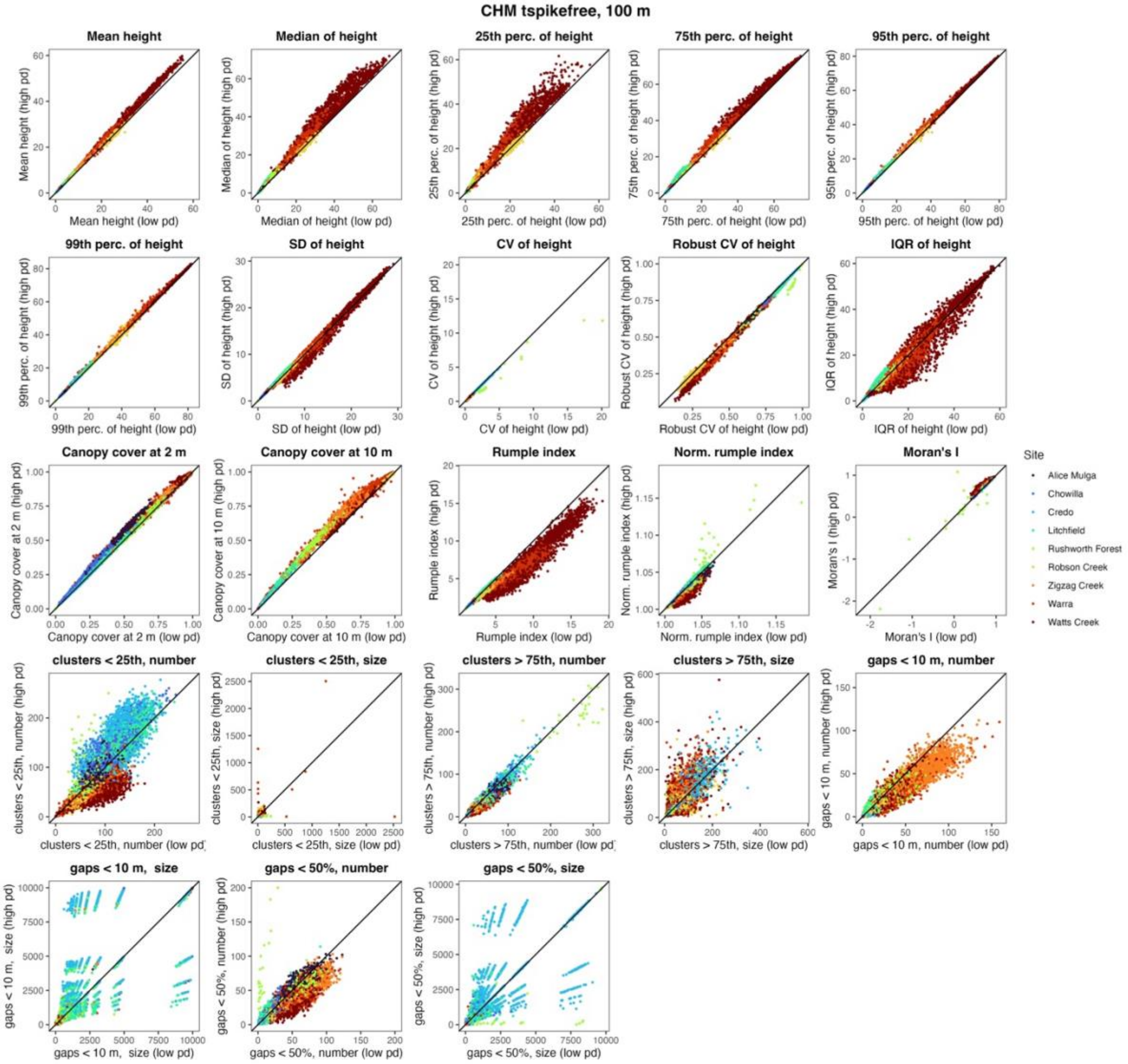

**Figure S14: Derived metrics for $CHM_{tspikefree}$ at 100 m scale.** Shown are derived metrics for $CHM_{tspikefree}$ at 100 m and across all sites together, compared between 2 pulses $m^{-2}$ (low pd) and 16 pulses $m^{-2}$ (high pd). Data points are coloured by site.

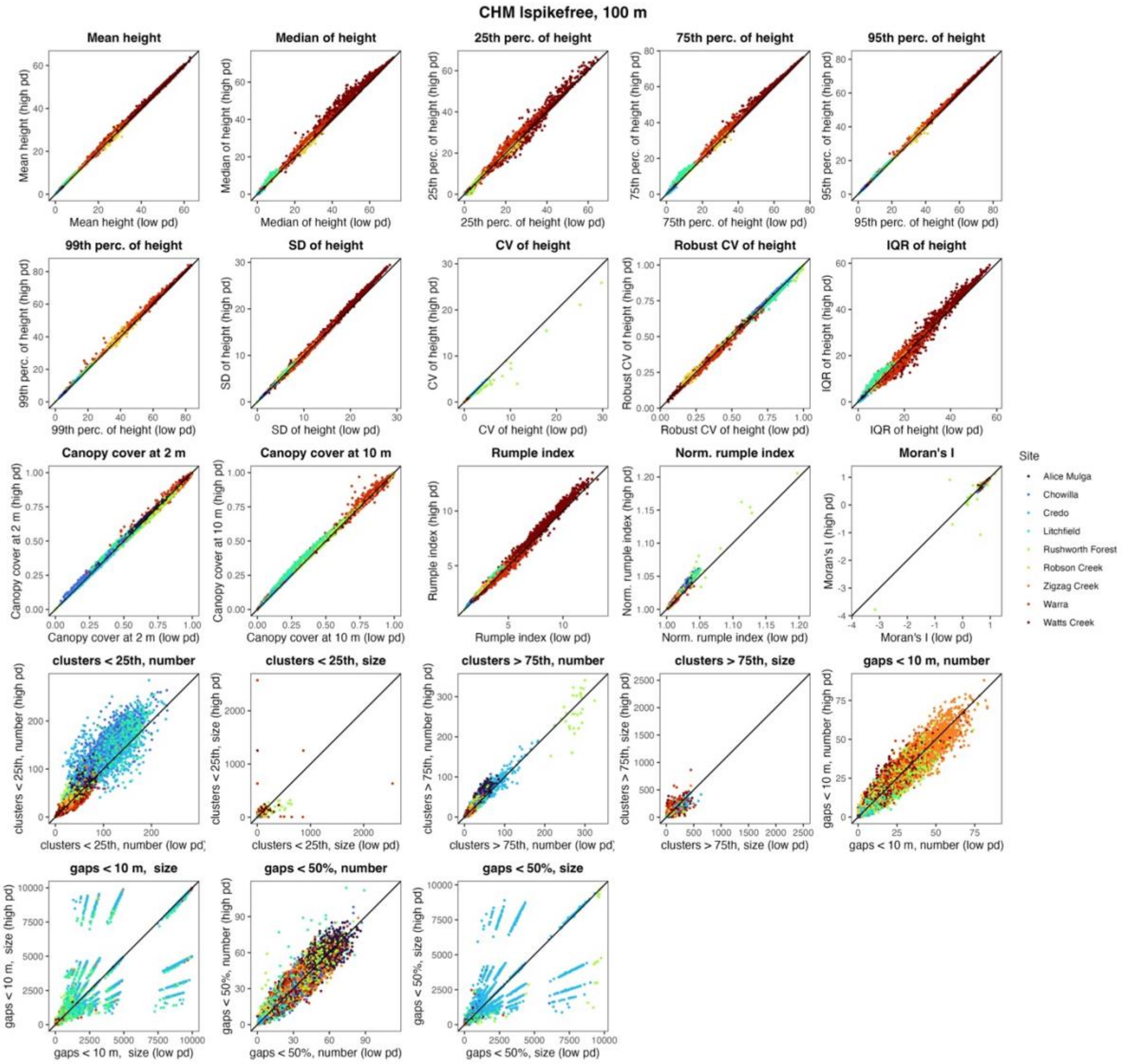

**Figure S15: Derived metrics for $CHM_{lspikefree}$ at 100 m scale.** Shown are derived metrics for $CHM_{lspikefree}$ at 100 m and across all sites together, compared between 2 pulses $m^{-2}$ (low pd) and 16 pulses $m^{-2}$ (high pd). Data points are coloured by site.

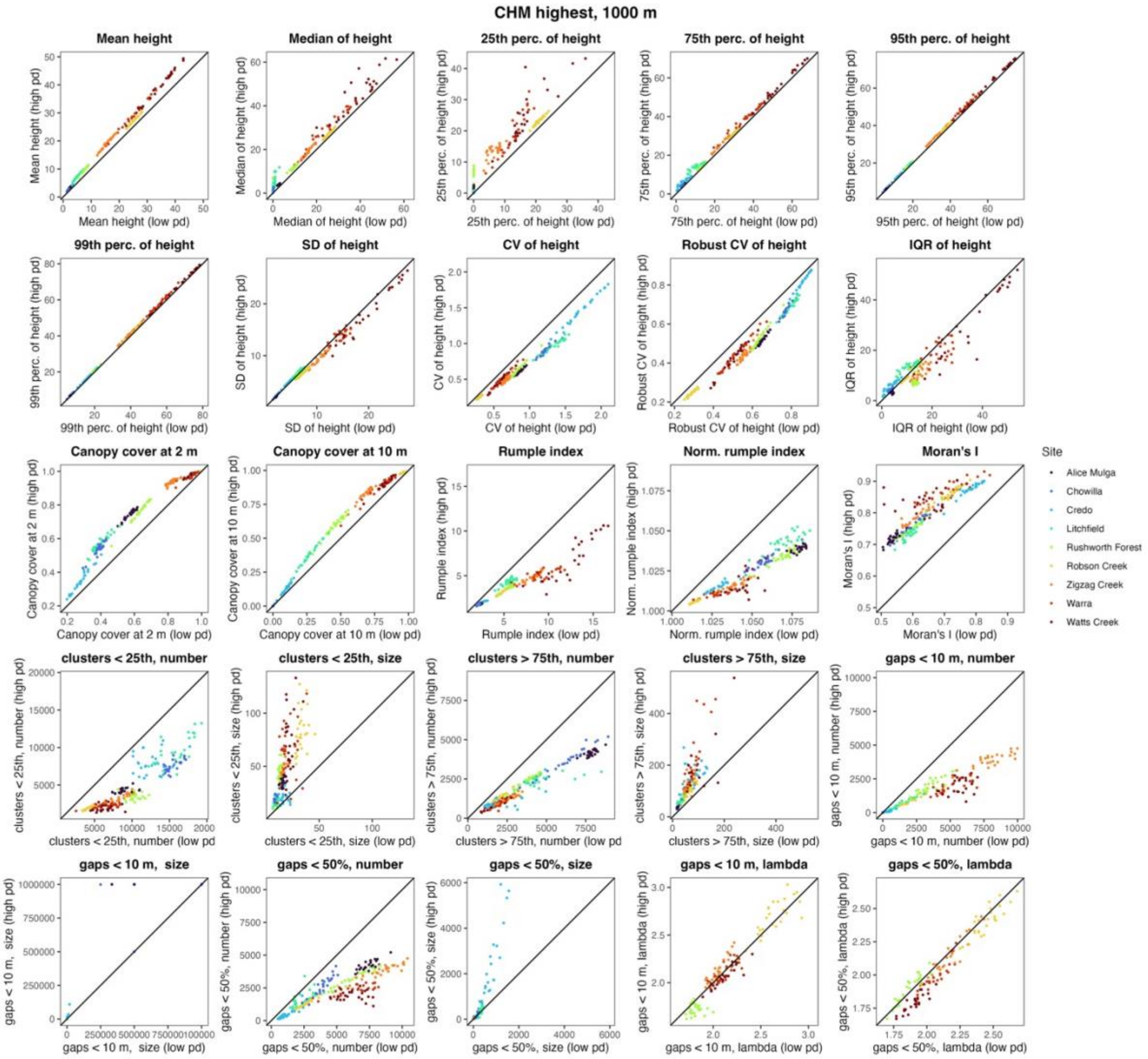

**Figure S16: Derived metrics for $CHM_{highest}$ at 1000 m scale.** Shown are derived metrics for $CHM_{highest}$ at 1000 m and across all sites together, compared between 2 pulses $m^{-2}$ (low pd) and 16 pulses $m^{-2}$ (high pd). Data points are coloured by site.

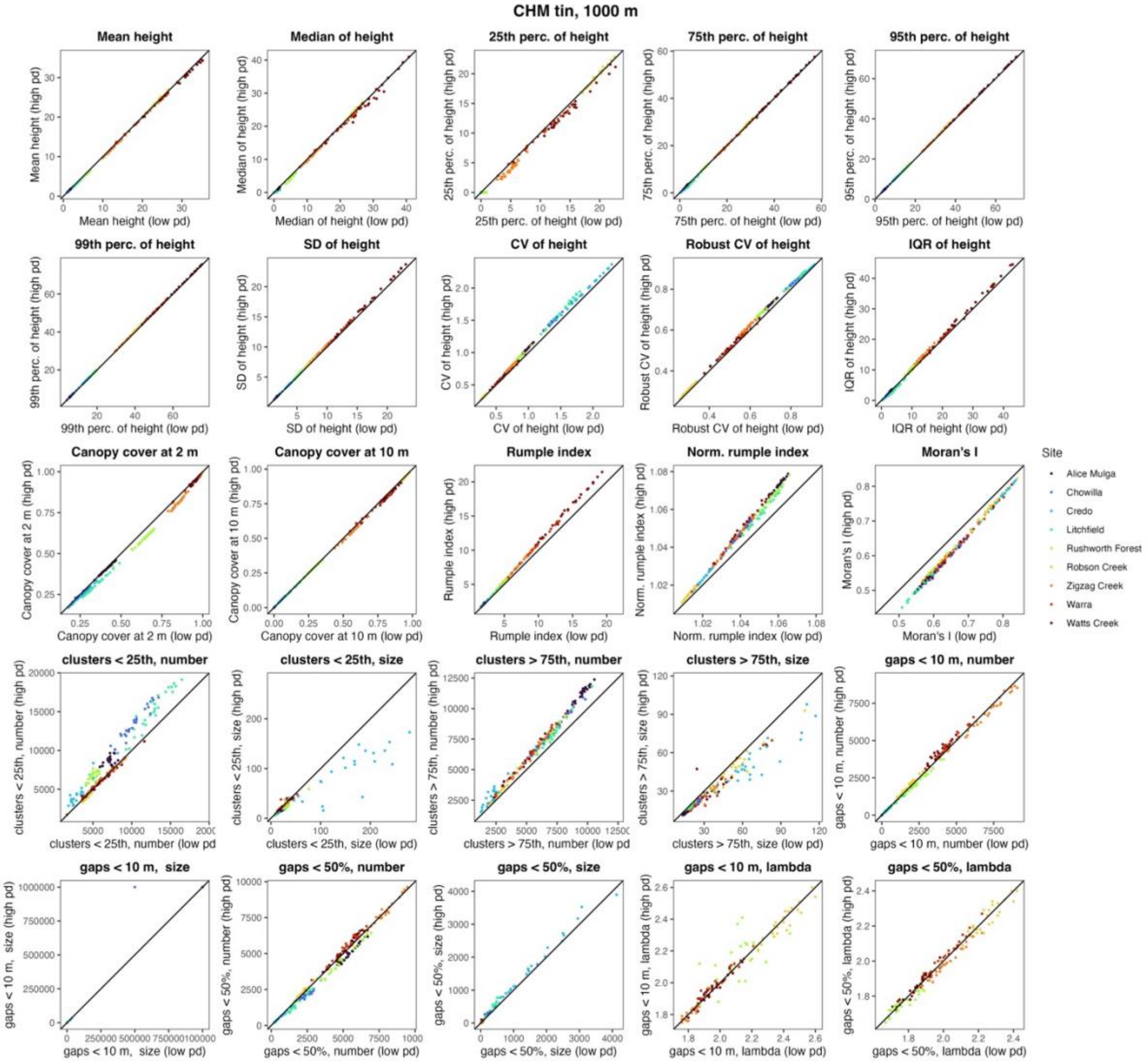

**Figure S17: Derived metrics for $CHM_{tin}$ at 1000 m scale.** Shown are derived metrics for $CHM_{tin}$ at 1000 m and across all sites together, compared between 2 pulses $m^{-2}$ (low pd) and 16 pulses $m^{-2}$ (high pd). Data points are coloured by site.

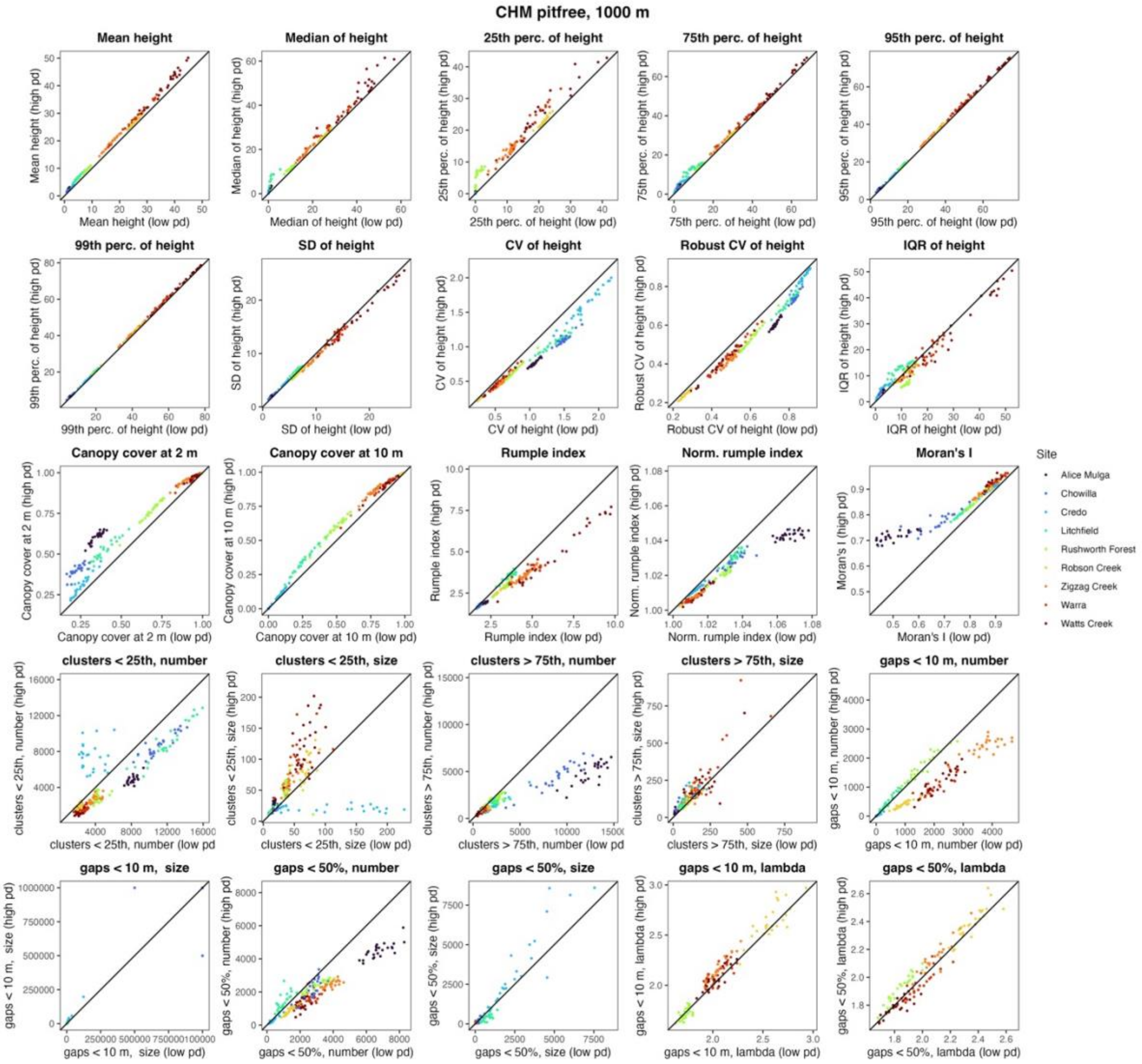

**Figure S18: Derived metrics for $CHM_{pitfree}$ at 1000 m scale.** Shown are derived metrics for $CHM_{pitfree}$ at 1000 m and across all sites together, compared between 2 pulses m$^{-2}$ (low pd) and 16 pulses m$^{-2}$ (high pd). Data points are coloured by site.

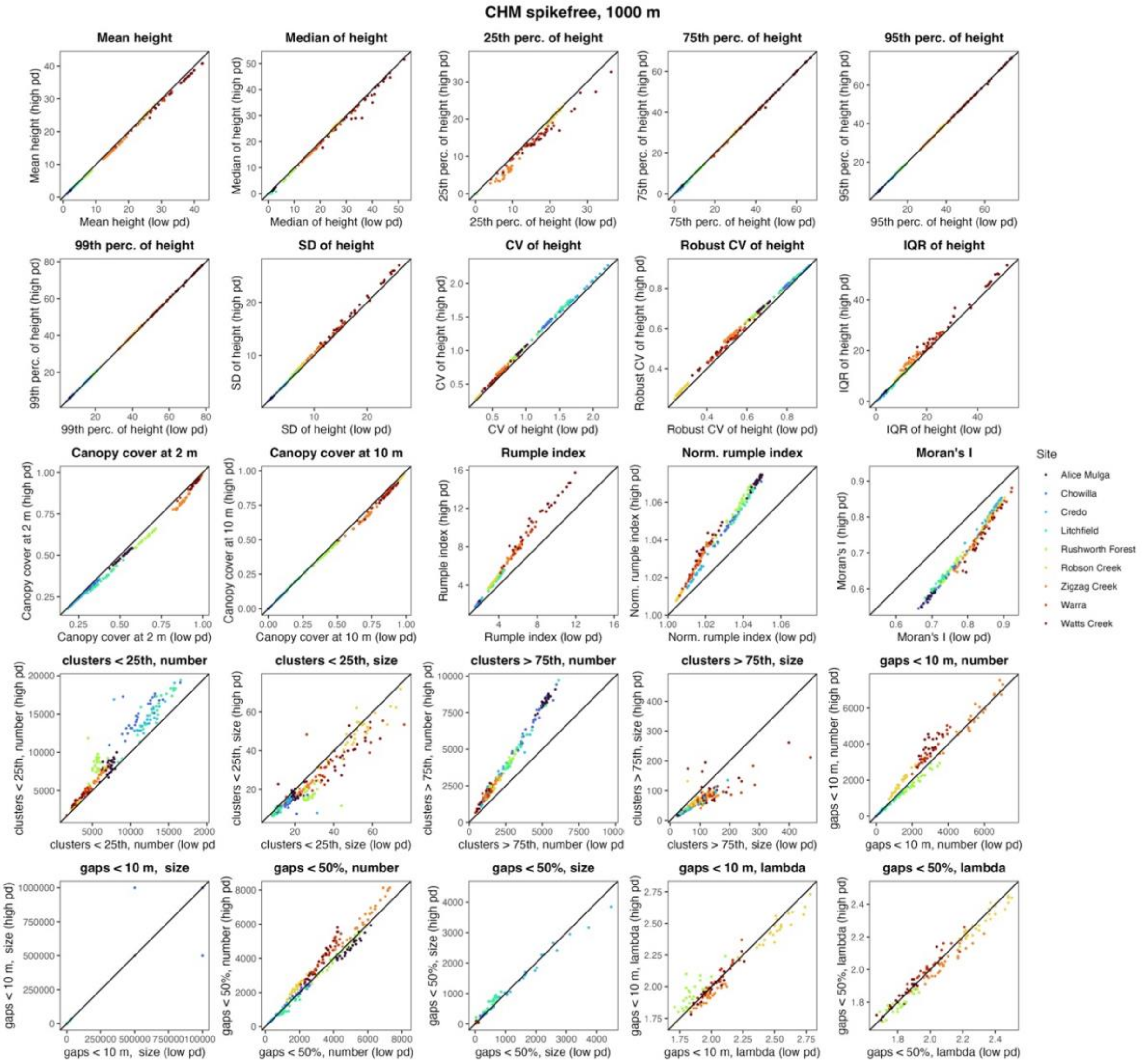

**Figure S19: Derived metrics for $CHM_{spikefree}$ at 1000 m scale.** Shown are derived metrics for $CHM_{spikefree}$ at 1000 m and across all sites together, compared between 2 pulses $m^{-2}$ (low pd) and 16 pulses $m^{-2}$ (high pd). Data points are coloured by site.

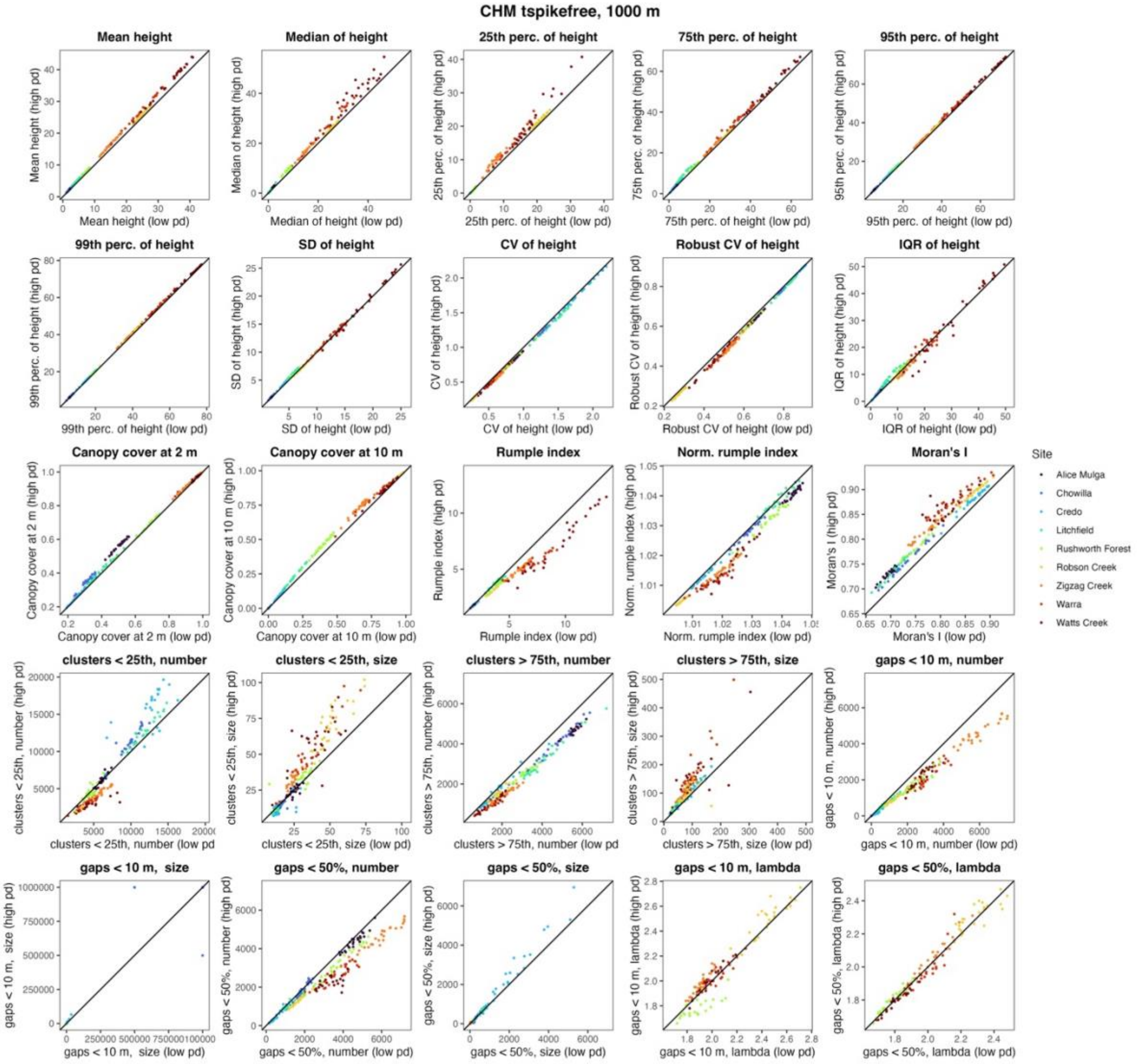

**Figure S20: Derived metrics for $\mathrm{CHM}_{\mathrm{tspikefree}}$ at 1000 m scale.** Shown are derived metrics for $\mathrm{CHM}_{\mathrm{tspikefree}}$ at 1000 m and across all sites together, compared between 2 pulses $m^{-2}$ (low pd) and 16 pulses $m^{-2}$ (high pd). Data points are coloured by site.

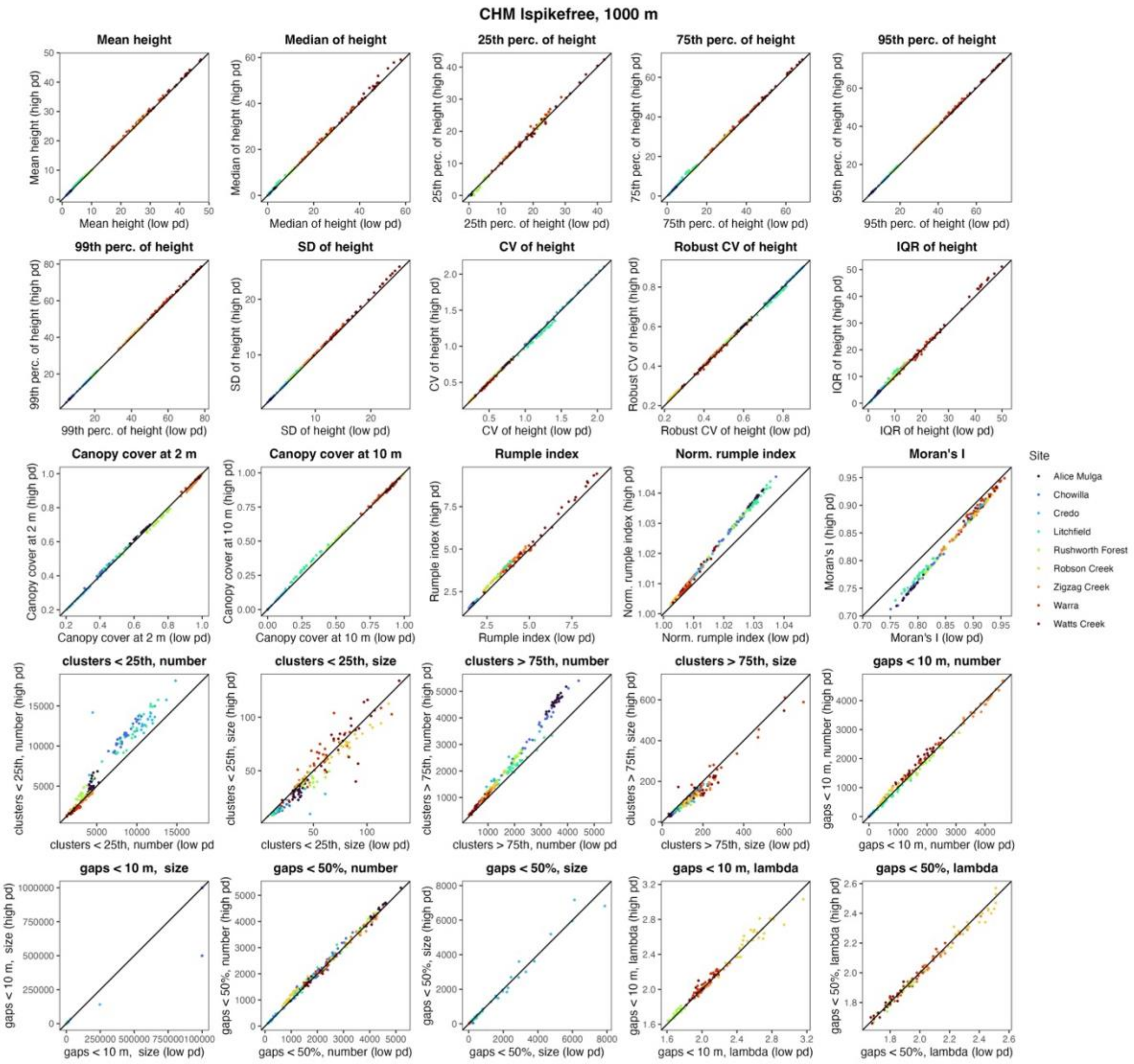

**Figure S21: Derived metrics for $CHM_{lspikefree}$ at 1000 m scale.** Shown are derived metrics for $CHM_{lspikefree}$ at 1000 m and across all sites together, compared between 2 pulses $m^{-2}$ (low pd) and 16 pulses $m^{-2}$ (high pd). Data points are coloured by site.

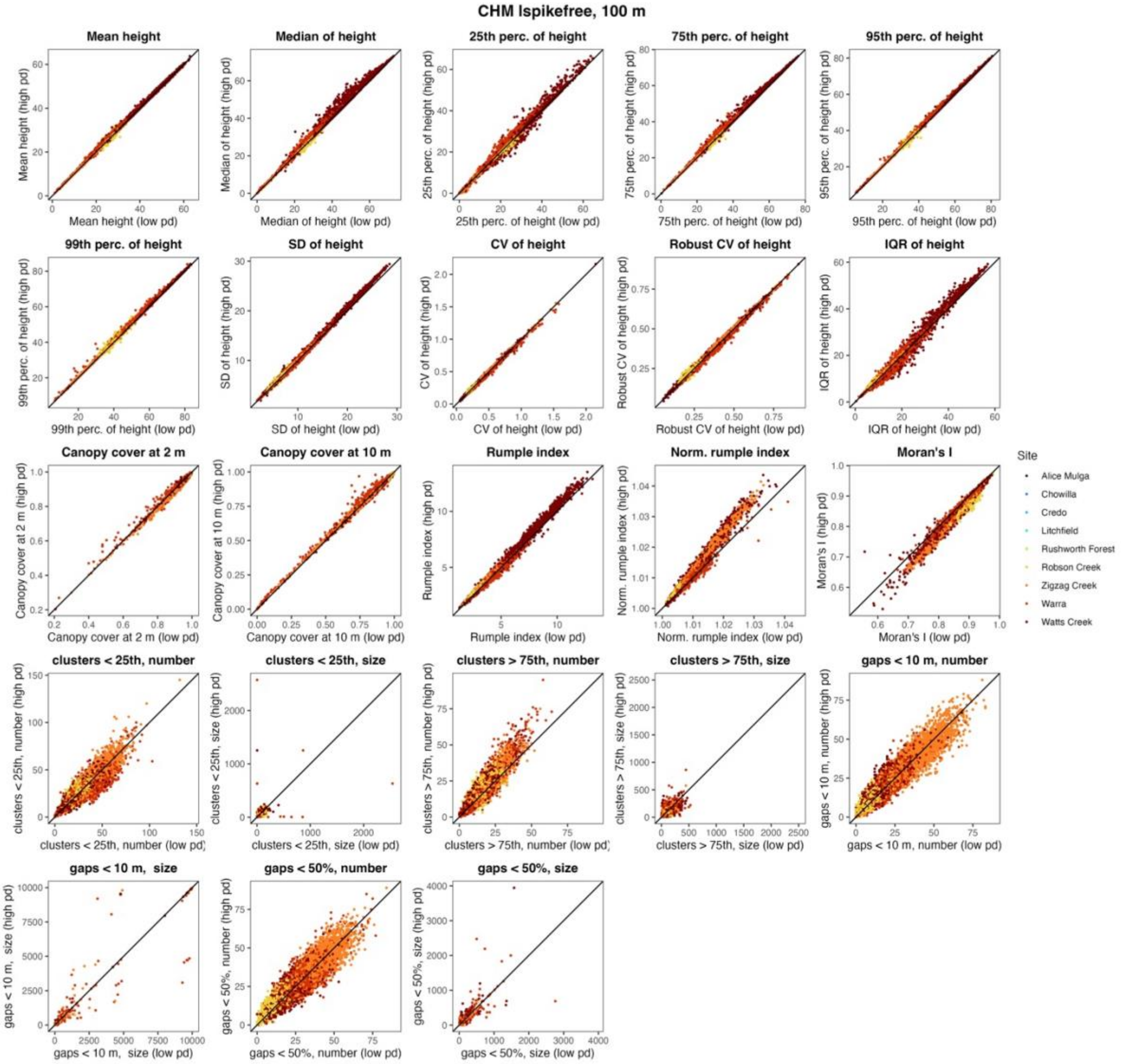

**Figure S22: Derived metrics for $CHM_{lspikefree}$ at 100 m scale, for closed-canopy forests only.** Same as Figure S15, but restricted to 4 sites (Robson Creek, Zigzag Creek, Warra, Watts Creek). Shown are derived metrics for $CHM_{lspikefree}$ at 100 m and across all sites together, compared between 2 pulses $m^{-2}$ (low pd) and 16 pulses $m^{-2}$ (high pd). Data points are coloured by site.

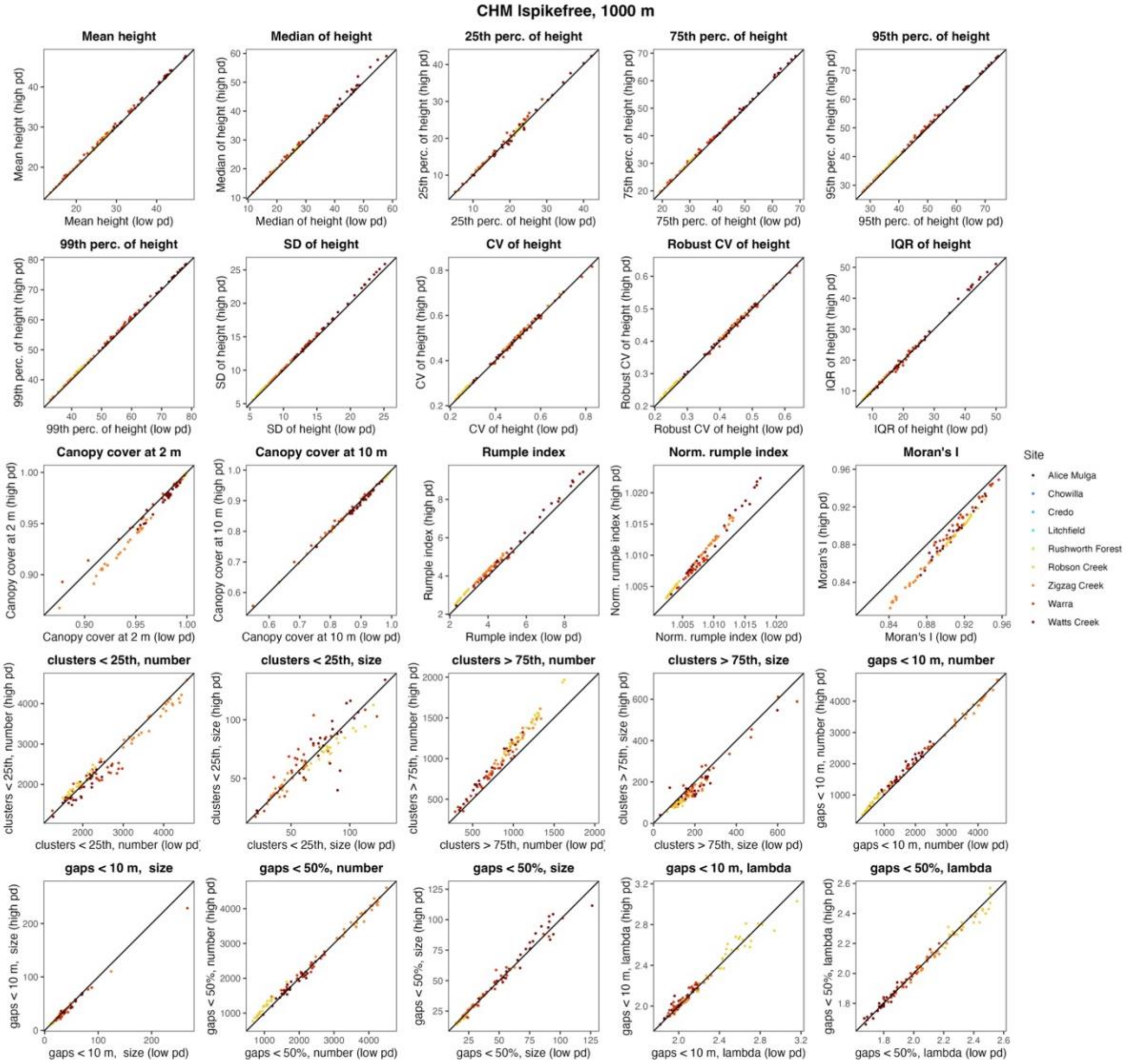

**Figure S23: Derived metrics for $CHM_{lspikefree}$ at 1000 m scale, for closed-canopy forests only.** Same as Figure S21, but restricted to 4 sites (Robson Creek, Zigzag Creek, Warra, Watts Creek). Shown are derived metrics for $CHM_{lspikefree}$ at 1000 m and across all sites together, compared between 2 pulses $m^{-2}$ (low pd) and 16 pulses $m^{-2}$ (high pd). Data points are coloured by site.
